# Diverse Diterpenoid Phytoalexins Shape Wheat Chemical Defenses

**DOI:** 10.64898/2026.05.16.720699

**Authors:** Rajesh Chandra Misra, Amr El-Demerdash, Amir Ronen, Oria Teena, Charlotte Owen, Benjamin D. Scott, Andrew Steed, Barrie Wilkinson, Paul Nicholson, Guy Polturak, Anne Osbourn

## Abstract

Cereal crops rely on a wide array of specialized metabolites to defend themselves against microbial pathogens. Here, using a combination of genome-wide analysis, heterologous pathway reconstruction, structural elucidation and *in planta* validation, we identify two pathogen-induced diterpenoid pathways in wheat that produce diterpenoids: the new glycosylated diterpenes aspisoside A and aspisoside B, and the diterpene alcohols scutenol A and scutenol B. The pathogen-responsive nature of these pathways, together with their antimicrobial activity are consistent with a likely defensive role in wheat. These compounds are produced by two biosynthetic gene clusters, each encoding a discrete pathway, so revealing organizational separation of wheat diterpenoid-based chemical defenses. The aspisoside-producing gene cluster is syntenic with diterpenoid phytoalexin-producing clusters in rice and barley, for momilactone and hordedane biosynthesis, respectively, yet gives rise to structurally distinct phytoalexins in wheat. The scutenol cluster is syntenic with currently uncharacterized predicted biosynthetic gene clusters in barley, oat, and *Brachypodium*. These findings establish diterpene glycosides as a previously unrecognized component of wheat defense chemistry and provide new insights into the chemical diversification of defense-related biosynthetic gene clusters within the Poaceae.

## INTRODUCTION

Plant natural products form a core component of plant defense, mediating interactions with pathogens, herbivores, and other environmental challenges, and encompass enormous chemical diversity^1–4^. Despite decades of research, our understanding of the kinds of chemicals that plants are capable of producing remains fragmented. Inevitably, many more plant defense metabolites await discovery, and for many of the known compounds, the biosynthetic pathways remain poorly characterized.

Among plant defense metabolites, diterpenoids represent a particularly important class in the grass family (Poaceae), forming a diverse arsenal of inducible chemical defenses in cereal crops such as rice, maize, and barley. Defense-related diterpenes have been most extensively studied in rice, where a suite of diterpene synthases (diTPS) generates different scaffolds that undergo further tailoring reactions to produce distinct types of compounds, such as momilactones, oryzalides, oryzalexins, and phytocassanes (**Fig. 1A**)^5–7^. Notably, momilactones, oryzalides and phytocassanes are produced in rice by biosynthetic gene clusters (BGCs), as are several other groups of defense-related compounds in grasses, including triterpenes, alkaloids, benzoxazinoids, and cyanogenic glucosides^8^. Maize similarly produces various types of labdane-related diterpenes, including the dolabralexin and kauralexin groups (**Fig. 1A**), the latter also showing antifungal and antifeedant activities^9–11^. Another group of diterpenoid phytoalexins, hordedanes (**Fig. 1A**), were also recently described in barley. Interestingly, hordedanes have been shown to inhibit the growth of the fungal pathogen *Fusarium graminearum* but enhance colonization by another fungal pathogen, *Bipolaris sorokiniana*. Notably, hordedane biosynthesis is encoded by a BGC homologous to the momilactone-producing cluster in rice^12^. In wheat, however, our understanding of diterpene metabolism is limited.

**Figure 1.**
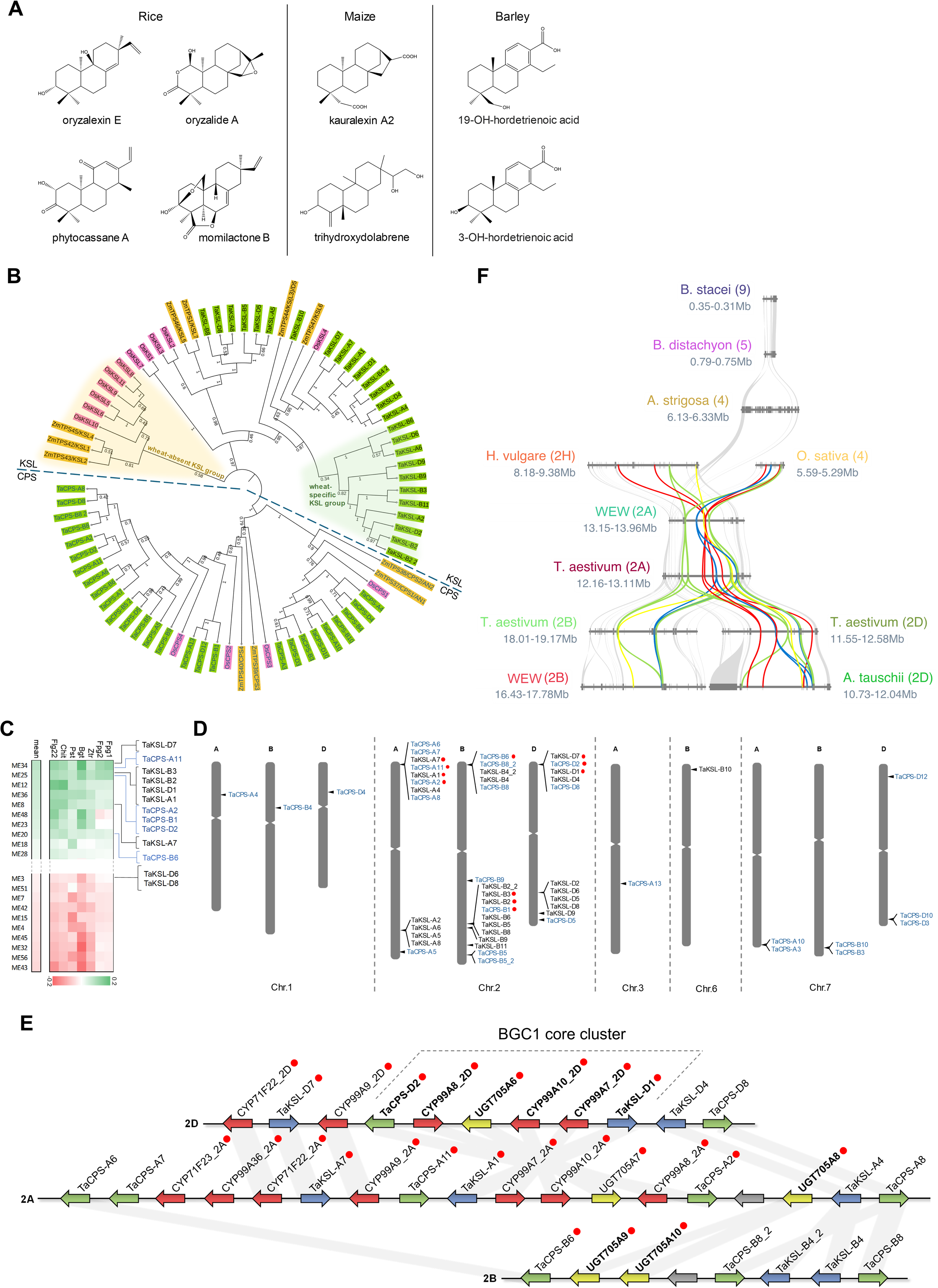
Genome-wide investigation of candidate wheat diterpene synthases reveals pathogen-induced diTPS genes co-localized in two biosynthetic gene clusters on chromosome 2. **A,** diterpenoid phytoalexins previously reported in rice, maize and barley. Representative molecules of larger compound groups are shown. **B,** phylogeny of all putative class I (KSL) and class II (CPS) diTPS genes identified in wheat. The maximum likelihood tree is comprised of diTPS genes from wheat (green), maize (yellow), and rice (pink). **C,** assignment of identified wheat CPS and KSL genes to gene expression modules of a biotic stress-related WGCNA. Gene expression modules are ordered from the most highly upregulated (top) to the most highly downregulated (bottom), following pathogen infection or treatment with chitin or flagellin. Only the top 10 and the bottom 10 modules are shown. The full figure can be found in **Supplementary Fig. 3.** Fpg1, *Fusarium pseudograminearum*; Fpg2, *Fusarium pseudograminearum*; Ztr*, Zymoseptoria tritici*; Bgt*, Blumeria graminis f. sp. tritici*; Pst*, Puccinia striiformis f. sp. Tritici*; Chit, chitin; Flg22, flagellin peptide. **D,** localization of all identified diTPS genes on the hexaploid wheat genome. Red dots indicate pathogen-induced genes, based on WGCNA data. E, BGC1, found on chromosome 2 of the A, B and D subgenomes. Red dots indicate pathogen-induced genes, based on WGCNA data. Gray lines connect homoeologous genes. Genes that have been functionally analyzed in the current study are marked in bold. F, microsynteny analysis of the BGC1 genomic region in group 2 chromosomes of hexaploid wheat (*Triticum aestivum*), and syntenic regions in rice (*Oryza sativa*), barley (*Hordeum vulgare*), *Triticum turgidum ssp. dicoccoides* (WEW; wild emmer wheat), black oat (*Avena strigosa*), *Aegilops tauschii*, and *Brachypodium* species (*B. distachyon* and *B. stacei*). Color coding of homologous genes is according to gene family: green, CPS; blue, KSL; red, CYP; yellow, UGT; gray, all other gene families.

The potential of wheat to produce diterpene phytoalexins was first demonstrated through functional characterization of several copalyl diphosphate synthase (CPS) and kaurene synthase-like (KSL) diterpene synthases, some of which exhibit UV-induced expression^13^. Four of the characterized wheat CPS enzymes were shown to catalyze the production of normal or *ent* stereoisomers of CPP, whereas five of the wheat KSL enzymes converted normal-, *ent*-, or *syn*- CPP into a range of different diterpene products^13–15^. More recently, we reported the occurrence of two putative BGCs at two different loci on chromosome 2 of hexaploid wheat, named BGC1(2A/2D) and BGC2(2B), but it remained unknown whether these clusters indeed encode biosynthetic pathways, what their metabolic products are, and how they might contribute to plant defense in wheat^16^.

Here, to gain a more comprehensive understanding of diterpenoid metabolism in bread wheat, we carried out a genome-wide investigation to identify all predicted diTPS genes. We identified 13 CPS and 11 KSL homoeologous gene groups, of which four CPS and four KSL groups were strongly induced in response to pathogen challenge. Intriguingly, all diTPS genes that were found to be pathogen-induced are co-localized in the two putative gene clusters, BGC1 and BGC2, which also comprise genes from the cytochrome P450 and UDP-dependent glycosyltransferase (UGT) families. We report the full functional characterization of the two BGCs and show by transient expression in *Nicotiana benthamiana* that they encode biosynthetic pathways for new wheat diterpenes. Specifically, BGC1 encodes a pathway for production of two normal-CPP-derived diterpene glycosides, here named aspisoside A and B, which are produced from a shared aglycone by two different UGTs. Both UGTs reside in the BGC1 cluster and belong to the monocot-specific UGT705 family. We further show that BGC2 produces *ent*-CPP-derived diterpenoids, here named scutenol A and B. Aspisoside A and B and scutenol A and B, along with other pathway products, were recombinantly produced in *N. benthamiana* in milligram quantities for structural assignment by NMR and for investigations of bioactivity. *In vitro* assays revealed that aspisosides A and B had antifungal activity, while scutenol A was antibacterial. The pathogen-induced expression of the diterpene-producing BGCs, coupled with the antimicrobial activity of their products, are suggestive of a role for these metabolites in defence in wheat. Notably, BGC1 is syntenic with BGCs that produce defensive diterpenoids in other cereal crops, including momilactones in rice and hordedanes in barley, but uniquely produces diterpene glycosides. BGC2 is syntenic with currently uncharacterized putative BGCs in other grasses. Together, these findings advance our understanding of wheat chemical defenses and uncover another layer in the evolutionary history of diterpene chemical diversification in the grasses.

## RESULTS

### Hexaploid wheat contains an extensive diTPS gene family

The first step in canonical diterpene biosynthesis involves the cyclization of the linear precursor geranylgeranyl pyrophosphate (GGPP) by diterpene synthases (diTPS) to form the cyclic hydrocarbon skeleton of diterpenoids. The initial step involves a protonation-triggered cyclization of GGPP (**1**) by class II diTPS, primarily facilitated by copalyl diphosphate synthase (CPS), resulting in the formation of a bicyclic labdadienyl or copalyl diphosphate (CPP) with a defined stereochemistry (*ent*, *syn*, *syn-ent*, or (+)/normal). Kaurene synthase-like (KSL) enzymes, belonging to class I diTPS, subsequently catalyze the cyclization and rearrangement of copalyl diphosphate (CPP) into a diverse array of diterpene scaffolds^17^. To carry out a systematic investigation of the capacity of bread wheat to produce diterpenes, we therefore first performed a genome-wide search to map all diterpene synthases (diTPS) in the hexaploid wheat genome (IWGSC refseq v1.1)^18^. An initial search for proteins containing a ‘terpene synthase’ domain (pfam domain pf01397; https://www.ebi.ac.uk/interpro/entry/pfam/PF01397/) retrieved 168 sequences. A maximum likelihood phylogenetic tree of all the predicted wheat TPS proteins indicated clear CPS and KSL branches containing 27 and 26 proteins, respectively, which were identified based on the presence of known wheat CPS and KSL genes (**Supplementary Fig. 1**). Based on further phylogenetic analysis, gene location on the wheat genome, and homoeology assignments in Ensembl Plants (http://plants.ensembl.org), we defined 13 CPS groups (each group containing 1-4 genes that include homoeologs or highly similar tandem duplicates), and 11 KSL groups (**Supplementary Fig. 1, Supplementary Data 1**). Four of the 27 predicted CPS proteins are atypically short (<550 amino acids) and are unlikely to be full-length sequences (**Supplementary Data 1**). These may represent pseudogenes or misannotations arising from ambiguities in genome assembly at their respective loci.

The wheat diTPS gene family is substantially larger than those identified in maize (4 CPS, 7 KSL genes)^19^ and rice (4 CPS, 11 KSL genes)^6,19^, even without taking into account the polyploidy-derived homoeolog diversity in wheat. For example, a single rice or maize CPS gene may have 3 orthologs in wheat, each positioned on a different subgenome (A, B, D). Therefore, wheat potentially harbours the capacity to produce a highly diverse array of as yet undiscovered diterpenoids. Phylogenetic comparisons with the full complement of rice and maize CPS and KSL protein sequences reveals a wheat-specific group of KSLs, and conversely a group comprised of maize and rice KSLs which includes the dolabralexin-related ZmTPS45/KSL4^10,20^ and OsKSL6/OsKSL10 involved respectively in oryzalide/oryzalexin biosynthesis^21–23^, but does not include any wheat sequences. The CPS proteins are divided into two main groups that contain sequences from all three species (**Fig. 1B; Supplementary Fig. 2**).

### Pathogen-induced diterpene synthases are co-localized in two biosynthetic gene clusters

We next analyzed gene expression data from a previously-generated weighted gene co-expression network analysis (WGCNA)^16,24^ to identify diTPS genes that are present in pathogen-induced gene expression modules, and are thus candidates for biosynthesis of diterpenoid phytoalexins. Five CPS genes from 4 groups (TaCPS1, TaCPS2, TaCPS6, TaCPS11), and 6 KSL genes from 4 groups (TaKSL1, TaKSL2, TaKSL3, TaKSL7) were found in pathogen-induced gene expression modules. (**Fig. 1C**, **Supplementary Fig. 3**). Notably, all of the induced genes were co-localized in two predicted biosynthetic gene clusters found on chromosome 2- one in the subtelomeric region of the short arm, and the other on the long arm (**Fig. 1D**). These loci respectively correspond with two putative gene clusters that we previously identified, BGC1 and BGC2^16^, and with the Fusarium-Responsive Gene Clusters FRGC 8/15 and FRGC 13 identified by Perochon and colleagues^25^. Interestingly, three KSL genes that are downregulated following pathogen infection (TaKSL-D5, TaKSL-D6, TaKSL-D8) are also co-localized on the D copy of BGC2 (**Fig. 1D**, **Supplementary Fig. 3**). Also notable was that the CPS and KSL genes from BGC1 showed pathogen-induced expression only in the copies of the A and D sub-genomes (with the exception of TaCPS-B6), while the CPS and KSL genes from BGC2 only showed pathogen-induced expression in the B genome copy (**Fig. 1D**). Further analysis of BGC1 revealed three homoeologous variants of a complex gene cluster, which in addition to the aforementioned diTPS genes also contains 12 cytochrome P450s from the diterpenoid-associated families CYP71 and CYP99^26^, and five UDP-dependent glycosyltransferases (UGTs). The 2A chromosome copy of the BGC is the largest, containing homoeologs of all Chr.2D cluster genes, and several additional CPS and CYP genes uniquely found in the Chr.2A copy, namely TaCPS-A6, TaCPS-A7, CYP71F23_2A, CYP99A36_2A and TaCPS-A11 (**Fig. 1E**).

### BGC1 is found on all wheat subgenomes and is syntenic with phytoalexin-producing gene clusters in rice and barley

Microsynteny analyses show that wheat BGC1 is homologous to and syntenic with the gene cluster that produces the diterpenoid phytoalexins, momilactones, in rice and with a more recently reported cluster that produces a structurally distinct group of diterpenoid phytoalexins, hordedanes (**Fig. 1F**)^12^. A wheat-focused microsynteny analysis further shows that BGC1 is generally conserved between the A and D copies, but that the B copy is largely truncated, and lacks any CYP450 genes. Correspondingly, the homologous BGCs in Chr.2A of wild emmer wheat (WEW; donor of the A and B genomes) and in *Aegilops tauschii* (donor of the D genome) both contain CPS, KSL, CYP and UGT genes, while the cluster on WEW Chr.2B also lacks any CYP450s. The syntenic regions in other grass genomes that we analyzed, namely diploid black oat (*Avena strigosa*), *Brachypodium distachyon* and *Brachypodium stacei* do not appear to contain any elements of the diterpene BGC found in wheat, rice and barley (**Fig. 1F**).

We next focused on the smaller of the pathogen-induced BGCs, BGC1(2D). This cluster is comprised of 11 genes with annotations relevant to diterpenoid biosynthesis, including three KSL and two CPS-type diterpene synthases, five cytochrome P450s and one UGT (**Fig. 1E**). TaKSL-D1 was used as bait in a gene co-expression analysis, with gene expression data obtained from wheat-expression.com^24^. Interestingly, the genes that were found to be most highly co-expressed with TaKSL-D1 were five of its directly flanking genes, including TaCPS-D2, three CYP99 genes CYP99A8_2D, CYP99A10_2D, CYP99A7_2D, and the glycosyltransferase UGT705A6 (**Supplementary Table S1**). We thus hypothesized that these six genes may form a ‘core cluster’ that encodes a shared biosynthetic pathway. This six-gene core cluster is also conserved in the A genome copy of BGC1, but in inverted orientation (**Fig. 1E**). Co-expression analysis using TaKSL-A1 similarly found that it is highly co-expressed with the five directly flanking genes, and an additional glycosyltransferase, UGT705A8 (**Supplementary Table S1**).

### The BGC1 core-cluster encodes a pathway for biosynthesis of two diterpene glycosides

To investigate the enzymatic function of the putative core cluster genes in BGC1(2D) and UGT705A8 from Chr.2A, the genes were cloned into a pEAQ-HT expression vector^27^, and transformed into *Agrobacterium tumefaciens* (GV3101) for functional analysis *via* transient co-expression (‘agroinfiltration’) in *Nicotiana benthamiana*^28^. The seven candidate genes (TaCPS-D2, TaKSL-D1, CYP99A8_2D, CYP99A10_2D, CYP99A7_2D, UGT705A6 and UGT705A8) were transiently co-infiltrated as a pool into *N. benthamiana* leaves. To facilitate the functional characterization of the candidate genes by increasing supply of the precursor GGPP (**1**), the pool of co-infiltrated genes additionally included the upstream MEP pathway gene geranylgeranyl diphosphate synthase from *Taxus canadensis* (TcGGPPS), and the truncated 3-hydroxy-3-methylglutaryl-coenzyme A reductase from *Avena strigosa* (AstHMGR), a rate-limiting enzyme in the mevalonate (MVA) pathway^29,30^. CPS and KSL genes were cloned without their predicted signal peptides to drive cytosolic rather than plastidial localization, for increased diterpene production^12,30,31^. Liquid chromatography-mass spectrometry-charged aerosol detection (LC-MS-CAD) analysis of infiltrated leaves indicated the formation of two peaks with retention times 7.05 and 7.64 min, when all genes are co-expressed together (**Fig. 2A**). To assess which of the genes are essential to the observed diterpene biosynthetic pathway, we carried out a ‘drop-out’ experiment by systematically removing one gene at a time during the heterologous reconstruction of the pathway in *N. benthamiana*. With the exception of CYP99A10_2D, all CPS/KSL/CYP genes appeared to be essential for formation of both putative diterpene glycosides. UGT705A6 and UGT705A8 were respectively required for producing the Rt 7.64 and Rt 7.05 compounds, thus indicating that each UGT specifically catalyzes the formation of one of the two compounds (**Fig. 2A**).

**Figure 2.**
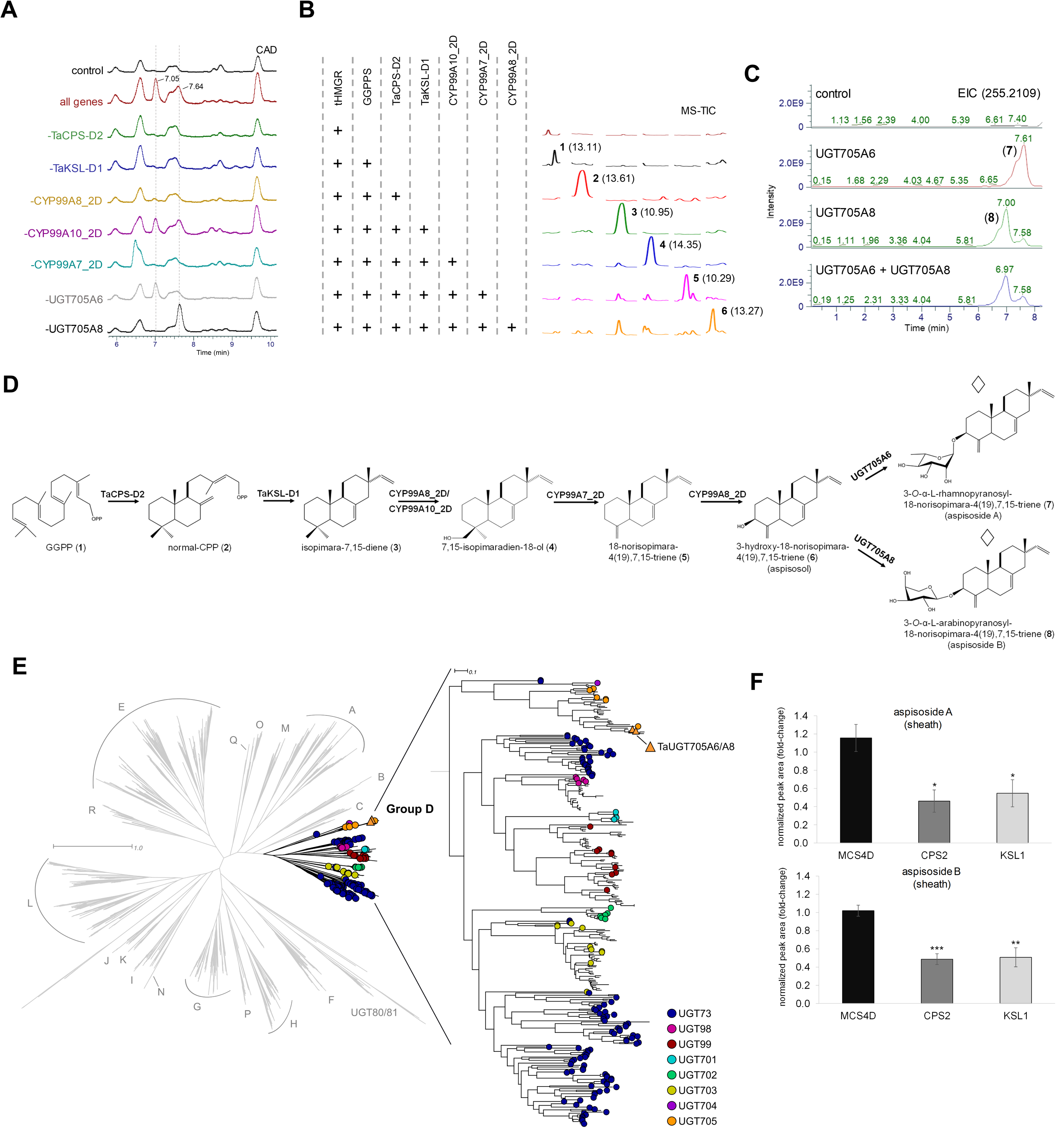
Gene cluster BGC1(2D) encodes a biosynthetic pathway for normal CPP-derived diterpene glycosides aspisoside A and aspisoside B. **A**, HPLC-CAD chromatograms of extracts from combinatorial transient expression experiments in *N. benthamiana* of BGC1(2D) genes. pEAQ-HT-DEST1 empty vector is used as a control. All genes- tHMGR, TcGGPPS, TaCPS-D2, TaKSL-D1. CYP99A7_2D, CYP99A8_2D, CYP99A10_2D, UGT705A6, UGT705A8. Remaining chromatograms are of drop-out experiments comprised of expression of all genes except one. Expression of all genes resulted in formation of two major peaks in the CAD chromatograms, with retention times 7.05 and 7.64. **B**, LC-MS analysis of leaf extracts from *N. benthamiana* combinatorial transient gene expression experiments. TIC, total ion chromatogram. Peak numbering corresponds with compound structures presented in panel D and **Supplementary Data S2**. Peak retention times are shown in brackets. +, indicates presence of gene in the expressed gene combination. **C**, LC-MS chromatograms of extracts from combinatorial transient expression experiments in *N. benthamiana* of BGC1(2D) genes. EIC, extracted ion chromatogram of a common fragment ion of aspisoside A and aspisoside B (*m/z* = 255.2109, [M + H]^+^). Chromatograms shown are of co-expression experiments of aspisosol-forming genes together with UGT705A6, UGT705A8, both UGTs, or none (control). **D**, proposed pathway for aspisosides biosynthesis from GGPP. Structures marked with a diamond sign were assigned by NMR analyses. **E**, relative quantification of aspisoside A and B in blade extracts from wheat plants following BSMV-mediated virus induced gene silencing of TaCPS-D2 and TaKSL-D1. Relative quantification (fold-change) values indicate means of four biological replicates ± SEM. Asterisks denote the statistical significance of a two-tailed *t*-test. \**P* < 0.05, \*\**P* < 0.01, \*\*\**P* < 0.001. **F**, Phylogenetic analysis of aspisoside-forming UGTs places them in group D family UGT705. 844 UGTs were mined from the *T. aestivum* genome using UDPGT Pfam PF00201 (https://www.ebi.ac.uk/interpro/entry/pfam/PF00201/). These were aligned with 246 reference UGTs from^33^ and 966 additional plant UGTs from the UGT Nomenclature Committee (https://labs.wsu.edu/ugt/) using MAFFT^60^ with a maximum of 1000 iterations, and a phylogeny generated from alignments with RaXML^61^ using the PROTGAMMAAUTO model and 100 bootstraps. Major UGT family groups are labelled.

### CYP99s from the core cluster catalyze sequential oxidations of isopimara-7,15-diene

We next carried out step-by-step reconstitution of the biosynthetic pathway of the putative diterpene glycosides by combinatorial expression in *N. benthamiana*. Infiltrated leaves were extracted and analyzed by gas chromatography-mass spectrometry (GC-MS). TaCPS-D2 and TaKSL-D1 were previously determined to sequentially cyclize geranylgeraniol (**1**) (*m/z*=290.2, Rt 13.11) to normal copalol (**2**) (*m/z*=290.2, Rt 13.61) and then isopimara-7,15-diene (**3**) (*m/z*=272.2, Rt 10.95)^16^. Here we observed that the additional expression of either CYP99A8_2D or CYP99A10_2D, but not CYP99A7_2D, to the isopimaradiene scaffold-forming-genes (AstHMGR, TcGGPPS, TaCPS-D2, TaKSL-D1) yielded a new, unidentified peak (**4**) *(m/z*=288.2, Rt 14.35). The CYP99s were further co-infiltrated in pairs together with the scaffold- forming-genes; expression of A8+A10 did not yield any additional peak to **4** that was produced by each of the two CYP99s alone, while expression of A7+A10 yielded a new unidentified peak (**5**) (*m/z*=256.2, Rt 10.29), and expression of A7+A8 yielded a different unidentified peak (**6**) (*m/z*=272.2, Rt 13.27). Co-infiltration of all three CYP99s similarly resulted in the accumulation of compound **6**, which we thus considered to be the final product of the cytochrome P450-mediated oxidation steps of the pathway (**Fig. 2B, Supplementary Fig. 4**). GC-MS mass spectra for compounds **1** - **6** are presented in **Supplementary Fig. 5**. BGC1-related genes and compounds are listed in **Supplementary Data 2** and **Supplementary Data 3**, respectively.

### UGT705A6 and UGT705A8 respectively catalyze rhamnosyl- and arabinosyl-transferase activity

We next tested the two candidate UDP-glycosyltransferase (UGTs) for their ability to glycosylate compound **6**, *via* agroinfiltration in *N. benthamiana* coupled with high-resolution LC-MS/MS analysis. Co-infiltration of the genes producing compound **6** together with UGT705A6, or with UGT705A8, respectively resulted in formation of new peaks (**7**) (*m/z* = 419.2791 [M + H]^+^) or (**8**) (*m/z* = 405.2632 [M + H]^+^). Compound **7** also appeared as a minor product of UGT705A8. Combined expression of UGT705A6 and UGT705A8 resulted in the accumulation of both compound **7** and compound **8**, with the latter being a more prominent peak (**Fig. 2C; Supplementary Fig. 6**). To determine the chemical structures of the glycosylated products formed through the action of these UGTs, we carried out large-scale vacuum-mediated agroinfiltration^32^ of ∼100 *N. benthamiana* plants with the same set of genes used in the above-described co-infiltration experiments, followed by solvent extraction and purification of products **7** and **8** from 136 g dry leaf material by flash chromatography and semi-preparative HPLC, respectively yielding 3.2 mg and 6.5 mg. Through extensive 1D and 2D nuclear magnetic resonance (NMR) analyses, we structurally assigned the products as 3-*O*-α-L-rhamnopyranosyl-18-norisopimara-4(19),7,15-triene (**7**), hereinafter named aspisoside A, and 3-*O*-α-L-arabinopyranosyl-18-norisopimara-4(19),7,15-triene (**8**), hereinafter named aspisoside B (**Supplementary Data 4**). The relative stereochemistry of the sugar residues was resolved based on coupling constants calculations and analysis of coupled HSQC values for the anomeric protons. Notably, these structural assignments also infer the structures of their precursors, and combined with the above-described LC-MS analyses allow us to propose a full biosynthetic pathway for aspisoside A and B: following the sequential cyclization of GGPP (**1**) to normal CPP (**2**) and then isopimara-7,15-diene (**3**), CYP99A8_2D and CYP99A10_2D redundantly catalyze 18-hydroxylation to give 7,15-isopimaradien-18-ol (**4**). Methanol elimination catalyzed by CYP99A7_2D then gives 18-norisopimara-4(19),7,15-triene (**5**), followed by 3-hydroxylation by CYP99A8_2D, yielding 3-hydroxy-18-norisopimara-4(19),7,15-triene (**6**) (hereinafter named aspisosol). Finally, the aspisosol aglycone undergoes glycosylation at the 3-*O* position with a rhamnose or arabinose sugar to afford aspisoside A (**7**), or aspisoside B (**8**), respectively catalzyed by UGT705A6 or UGT705A8 (**Fig. 2D**).

### BGC1(2B) contains two additional active glycosyltransferases

The B genome copy of BGC1 is substantially smaller compared to the A and D copies, and includes seven putative diterpenoid-associated genes, of which only three were found to be pathogen-induced in the WGCNA-derived data (**Fig. 1E**). These included two UGTs, UGT705A9 and UGT705A10, that are homoeologs of UGT705A8, and the diTPS gene TaCPS-B6. Given their pathogen-induced expression, we proceeded to functionally analyze these by transient expression in *N. benthamiana*. Co-infiltration of the aspisosol-forming genes (AFG) together with UGT705A9 or with UGT705A10 led to the formation of aspisoside B, but not aspisoside A. Both enzymes are thus arabinosyltransferases that produce the same product as their A genome homoeolog, UGT705A8 (**Supplementary Fig. 7**). TaCPS-B6, however, was found to be enzymatically inactive, possibly explained by altered splicing resulting in the inclusion of an 18 amino acids-long segment in its protein sequence, which is not found in sequences of related CPS enzymes (**Supplementary Figs. 8, 9, 10**). Extensive phylogenetic analysis of all wheat UDP-dependent glycosyltransferases places the BGC1 UGTs in Group D, associated with triterpenoid and flavonoid glycosylation^33^, and more specifically in the UGT705 family, a monocot-specific clade^34^ that currently includes only three functionally characterized UGTs involved in glycosylation of the flavonoid tricin in rice^35^ (**Fig. 2E**).

### Viral Induced Gene Silencing of TaCPS-D2 and TaKSL-D1 decreases aspisoside accumulation

To further investigate the roles of TaCPS-D2 and TaKSL-D1 in biosynthesis of aspisosides in wheat, we carried out viral induced gene silencing (VIGS) using the Barley Stripe Mosaic Virus (BSMV) system^36,37^. TaCPS-D2 and TaKSL-D1 were cloned into pCa-γbLIC in antisense orientation, resulting in the constructs pCa-γb::TaCPS-2D and pCa-γb::TaKSL-D1, respectively whereas pCa-γb::mcs4D (containing a non-coding DNA sequence) was used as a negative control. The construct pCa-γb::PDS, for silencing the wheat phytoene desaturase gene was used as positive control for viral infection, due to its expected visible bleaching phenotype (**Supplementary Fig. 11**). To investigate the consequences of gene silencing on diterpene biosynthesis, relative aspisoside production was assessed using LC-MS. Analysis of TaCPS-D2- or TaKSL-D1- silenced wheat blade tissues revealed a ∼50% to 60% reduction in aspisoside A and aspisoside B content. This decrease supports an essential role of TaCPS2_2D and TaKSL1_1D in the biosynthesis of the two glycosylated diterpenoids (**Fig. 2F**). Successful silencing of TaCPS-D2 and TaKSL-D1 in the sampled tissue was further assessed using quantitative reverse-transcription PCR (qRT-PCR). Significant reduction of the transcript level was respectively detected for TaCPS-D2 (∼80%) and TaKSL-D1 (∼90%) genes in pCa-γb::TaCPS-D2 and pCa-γb::TaKSL-D1 inoculated plants, relative to the values from pCa-γb::mcs4D-infected control tissue (**Supplementary Fig. 11**).

### Aspisoside production is induced by infection with fungal pathogens

Fungal pathogen infection in cereals such as rice, wheat, and maize induces specific biosynthetic pathways that produce antimicrobial compounds, primarily flavonoids, terpenoids (diterpenes and triterpenes) and other phenolic compounds (*e.g*., phenylamides) which act as phytoalexins and are often encoded by pathogen-induced biosynthetic gene clusters^6,16,38–40^. As noted above, analysis of available transcriptomic data from pathogen infection experiments of wheat reveals a pathogen-induced expression pattern for BGC1 genes (**Supplementary Fig. 12**). To further investigate the pathogen-responsive expression of BGC1 genes, 3-week-old cv. Chinese Spring wheat plants were infected with the fungal pathogen *Fusarium culmorum* (isolate FC2021), by applying a thick agar-based, mycelium -containing slurry into plastic tubes placed around plant stem bases. Total RNA was extracted from blade samples 3- and 7- days post inoculation (dpi) and the expression levels of seven genes from cluster BGC1(2A/2D) was measured by quantitative real-time PCR (qRT-PCR). These experiments revealed strong induction of all seven genes in infected plants when compared with mock-inoculated plants at both timepoints, with stronger induction observed at 3 dpi (**Fig. 3A**). To examine the relative accumulation of aspisoside A (**7**) and aspisoside B (**8**) following *F. culmorum* infection in wheat, blade and sheath tissues were sampled at 7- and 14- dpi, and extracts analyzed by LC-MS. Both compounds were detected in blade and sheath extracts at both timepoints (**Supplementary Fig. 13**). Semi-quantitative analysis showed a ∼5-fold higher level of aspisoside A in *F. culmorum*-infected plants compared to mock-treated plants at 14 dpi, while no significant differences in aspisoside B levels were observed between mock and *F. culmorum*-infected plants in either tissue at either timepoint (**Supplementary Fig. 13**). To further investigate aspisoside accumulation following fungal pathogen infection, we next infected 3-week-old cv. Chinese Spring plants with a conidial suspension of *Magnaporthe oryzae* (strain BTJ4P), a well-characterized fungal pathogen known to cause blast disease in cereals^41^. Sheath and blade tissues were sampled from infected and mock-treated plants at 4- and 10- dpi, and extracts were analyzed by LC-MS. Aspisosides A and B both induced following *M. oryzae* infection and were detected both in sheath and blade tissues of infected plants at 10 dpi (**Fig. 3B**), but barely detected at 4 dpi. Notably, the peaks representing the two aspisosides were among the most prevalent peaks in the APCI total ion chromatogram of the 10 dpi *M. oryzae*-infected samples (**Supplementary Fig. 14)**. The observed peaks in the tissue extracts were identified and confirmed by comparison with purified standards (**Fig. 3B, Supplementary Fig. 14**). Taken together, our results suggest that BGC1(2A/2D) encodes a biosynthetic pathway for diterpenoid glycosides that is activated during pathogen attack.

**Figure 3.**
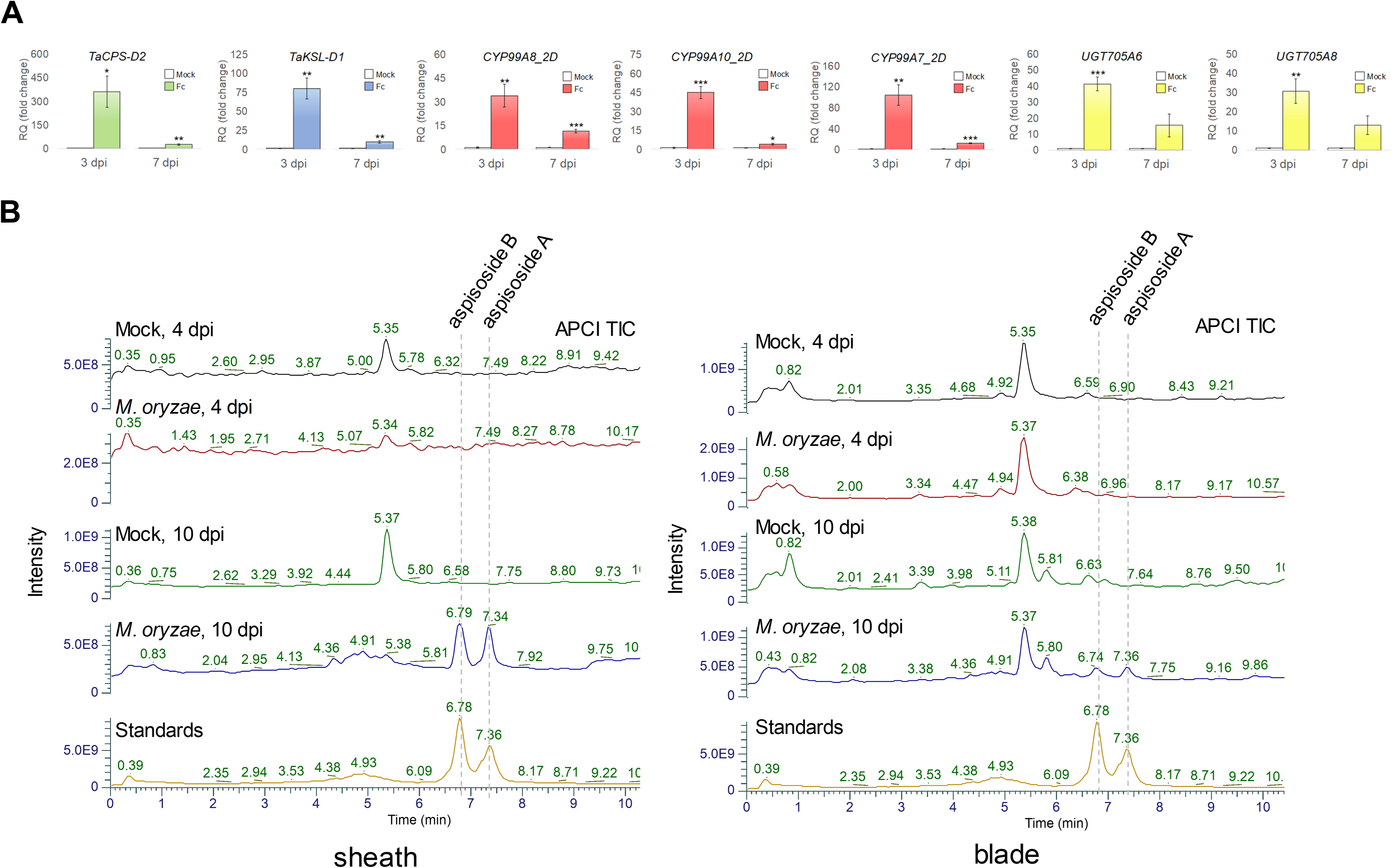
Induction of aspisoside biosynthesis in pathogen-infected wheat. **A**, quantitative real-time PCR of seven genes from cluster BGC1(2A/2D) in blade extracts from mock- or *F. culmorum*- treated wheat plants, 3- and 7-days post infection. Fc, *Fusarium culmorum*. Relative quantification (fold-change) values indicate means of three biological replicates ±SEM. Asterisks denote the statistical significance of a two-tailed *t*-test. \**P* < 0.05, \*\**P* < 0.01, \*\*\**P* < 0.001. **B**, LC-MS analysis of aspisoside A and B accumulation in sheath and blade tissues of wheat plants infected with *M. oryzae*, 4 or 10 days post infection. TIC, total ion chromatogram.

### BGC2 encodes two branching diterpenoid biosynthetic pathways

The other putative diterpene-producing biosynthetic gene cluster that we identified, BGC2(2B), is found on a different locus on chromosome 2 of the B subgenome. BGC2(2B) is comprised of likely diterpene-associated genes including one copalyl diphosphate synthase (CPS), six kaurene synthase-like genes (KSLs), and four cytochrome P450 monooxygenases (CYPs) (**Fig. 4A**). Of these, one CPS gene (TaCPS-B1), two KSL genes (TaKSL-B2 and TaKSL-B3), and two CYP genes (CYP99A39_2B and CYP701A64_2B) showed pathogen-responsive expression in the analyzed WGCNA data (**Fig. 4A**). Partial copies of the BGC2 cluster are also found in the A and D subgenomes, each comprised of four KSL genes and one CYP99. However, none of the A and D subgenome homoeologs appear to be pathogen-induced (**Fig. 4A**). TaCPS- B2 was previously shown to function as an *ent*-CPP synthase producing *ent*-CPP (**9**) from GGPP (**1**), while TaKSL-B2 was found to catalyze cyclization of *ent*-CPP (**9**) to produce *ent*-pimara-8(14),15-diene (**10**)^14–16^. Zhou and colleagues also reported another wheat kaurene synthase-like gene, TaKSL3, which exhibited only residual activity, producing two diterpenes that could not be identified due to their low accumulation levels^15^. Interestingly, TaKSL3 (hereinafter TaKSL-B3) resides in BGC2(2B), directly flanking TaKSL-B2^16^. Given the clear pathogen-responsive expression of TaKSL-B3 in the analyzed WGCNA data, we hypothesized that it may encode an active enzyme. To examine this, we cloned an N-terminal-truncated TaKSL-B3 into the pEAQ-HT expression vector for transient expression in *N. benthamiana*. GC-MS analysis of infiltrated *N. benthamiana* leaves following co-expression of TaCPS-B2 and TaKSL-B3 together with tHMGR and GGPPS resulted in the detection of a new peak (**12**) (*m/z*=272.25) (**Fig. 4B; Supplementary Fig. 15**). To gain more insights into substrate specificity and catalytic scope of the characterized TaKSL enzymes, we conducted additional co-expression experiments. Specifically, TaCPS-D2 and TaCPS-B1 were co-expressed with TaKSL-D1, TaKSL-B2 and TaKSL-B3. Neither TaKSL-B2 nor TaKSL-B3 react with normal CPP (**2**). Conversely, TaKSL-D1 from BGC1 does not react with *ent*-CPP (**9**) (data not shown). Of the four cytochrome P450s in BGC2(2B), two showed pathogen-responsive expression in the WGCNA data and in pathogen infection experiments from the Wheatomics database ^42^, namely CYP701A64_2B and CYP99A39_2B (**Fig. 4A, Supplementary Fig. 16**). Comprehensive functional analysis of the pathogen-induced genes in BGC2(2B) was thus undertaken, *via* a set of combinatorial co-infiltration experiments in *N. benthamiana*. Combined expression of TaCPS-B2, TaKSL-B2 and CYP701A64_2B resulted in production of a new peak (**11**), which we anticipated to be a hydroxylated product of *ent*-pimara-8(14),15-diene (**10**) based on its mass (*m/z*=288.25). Combined expression of TaCPS-B2, TaKSL-B3, and CYP99A39_2B led to the formation of another product (**13**) (*m/z*=288.25), which we hypothesized to be a hydroxylation product of compound **12** (**Fig. 4B; Supplementary Figs. 14, 16**). To assign structures for compounds **11** and **13,** we proceeded to produce them in *N. benthamiana* by vacuum-mediated large-scale infiltration. From a total of 75 infiltrated *N. benthamiana* plants for each gene combination, we recovered 4.3 mg of compound **11** and 95 mg of compound **13**. The structures were determined by extensive 2D NMR as *ent*-pimara-8(14),15-diene-19-ol (**11**), hereinafter named scutenol A and (Z)-biformene-3α-ol (**13**), hereinafter named scutenol B (**Fig. 4C**) (**Supplementary Data 4**). The assigned structure for **11** corresponds with the assigned structure for its precursor **10**, with the only difference between the two being the addition of a hydroxyl group at carbon number 19. CYP701A64_2B thus catalyzes 19-hydroxylation of the *ent*-pimaradiene scaffold. Assignment of the structure of **13** also allows assignment of the structure of its precursor **12** to be (Z)-biformene, and therefore to deduce that TaKLS-3B is a (Z)-biformene synthase and CYP99A39_2B is a (Z)-biformene-3-hydroxylase (**Fig. 2C**). The occurrence and distribution of scutenol A and scutenol B in wheat tissues was investigated by GC-MS analysis of blade and sheath extracts from *M. oryzae*-infected plants. These analyses revealed the presence of distinct peaks corresponding to scutenol A and scutenol B in infected tissues at 10 dpi, whereas these compounds were below the detection level in mock-treated plants. Interestingly, scutenol A was specifically found in sheath extracts, while scutenol B was uniquely identified in blades (**Fig. 4D**). Similarly to the aspisosides, scutenol A and B were not detected in any of the 4 dpi sample extracts. The peaks at 10 dpi were identified based on retention time and mass spectra comparison with scutenol A and scutenol B standards purified from *N. benthamiana* (**Fig. 4D, Supplementary Fig. 17**).

**Figure 4.**
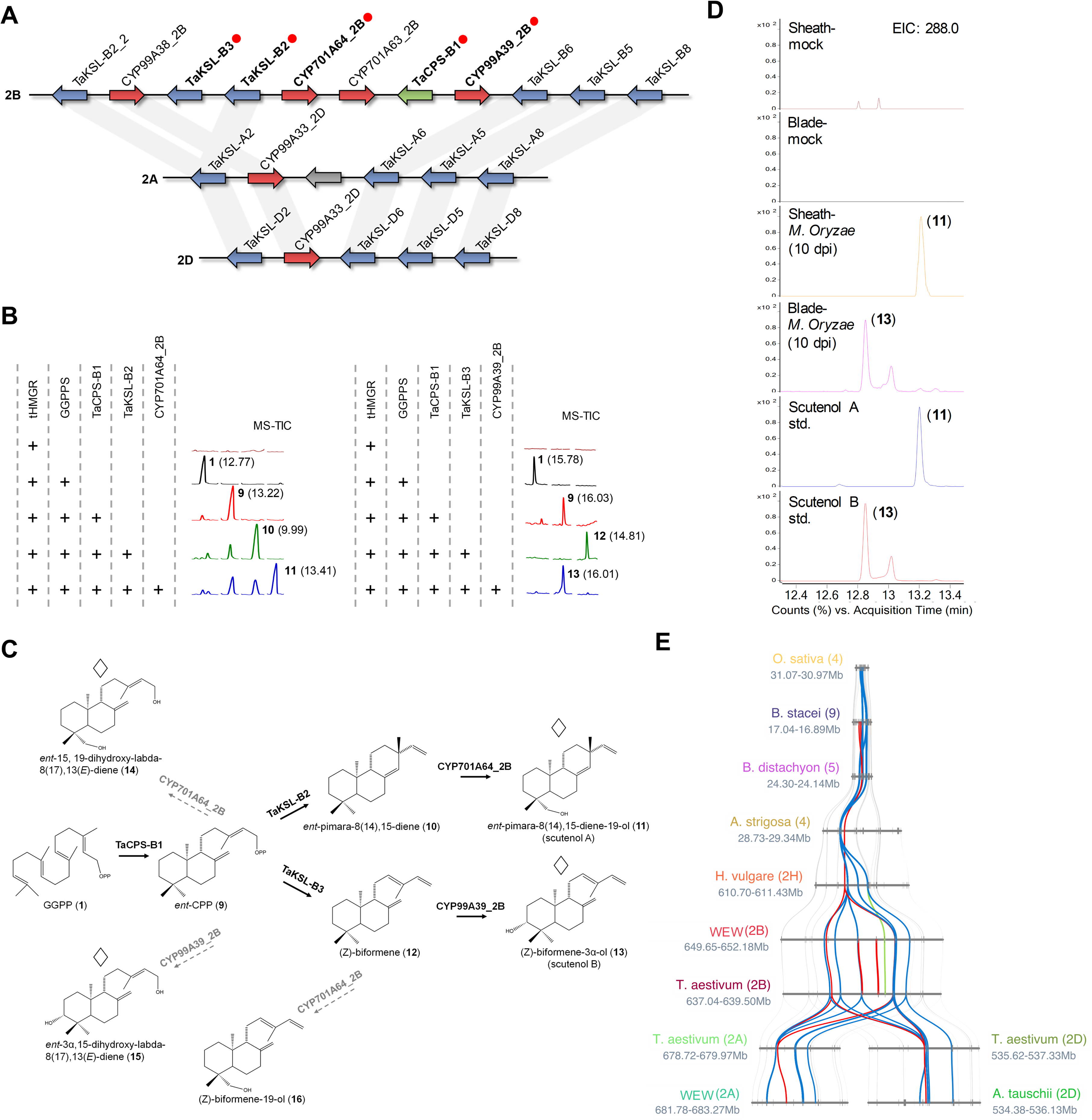
Gene cluster BGC2(2B) encodes a biosynthetic pathway for *ent*-CPP-derived diterpenoids scutenol A and scutenol B. **A**, BGC2 found on chromosome 2 of the A,B and D subgenomes. Red dots indicate pathogen-induced genes, based on WGCNA data. Gray lines connect homoeologous genes. Functionally analyzed genes are marked in bold. **B**, LC-MS analysis of leaf extracts from *N. benthamiana* combinatorial gene expression experiments. TIC, total ion chromatogram. Peak numbering corresponds with compound structures presented in panel D and **Supplementary Data 2**. Peak retention times are shown in brackets. +, presence of gene in the expressed gene combination. Left, TaKSL-B2-mediated pathway to scutenol A. Right, TaKSL-B3-mediated pathway to scutenol B. **C**, proposed pathway for scutenol biosynthesis from GGPP. Structures marked with a diamond sign were assigned by NMR analyses. **D**, GC-MS analysis of scutenol A and B accumulation in sheath and blade tissues of wheat plants infected with *M. oryzae*, 10 days post infection. EIC, extracted ion chromatogram. **E**, microsynteny analysis of BGC2 genomic region in group 2 chromosomes of hexaploid wheat (*Triticum aestivum*), and syntenic regions in rice (*Oryza sativa*), barley (*Hordeum vulgare*), *Triticum turgidum ssp. dicoccoides* (WEW; wild emmer wheat), black oat (*Avena strigosa*), *Aegilops tauschii*, and *Brachypodium* species *B. distachyon* and *B. stacei*. Color coding of homologous genes is according to gene family: green, CPS; blue, KSL; red, CYP; gray, all other gene families.

### BGC2(2B) CYPs can directly hydroxylate *ent*-CPP

Previous work demonstrated that cytochrome P450 enzymes could catalyze specific modifications of primary diterpene alcohol intermediates (*i.e*., CPS products) by adding a furan ring to form furanoditerpenoids^43^. Therefore, we sought to investigate whether CYP701A64_2B and CYP99A39_2B could potentially act directly on the TaCPS-2B product *ent*-CPP, thereby enabling production of novel diterpenoid derivatives. To test this, we co-expressed each of the two P450s with TaCPS-B2 in *N. benthamiana*. Remarkably, both CYP701A64_2B and CYP99A39_2B demonstrated activity on *ent*-CPP (**9**), giving rise to two new products, **14** (*m/z*=306.26) and **15** (*m/z*=306.26), respectively (**Fig. 4C, Supplementary Fig. 18, Supplementary Data 3**). Following large-scale vacuum-mediated infiltration, the two products were purified from freeze-dried *N. benthamiana* leaves: 0.5 mg of compound **14** was purified from 80 gr dry material, and 4 mg of compound **15** was purified from 100 gr dry material. The structures of the two compounds were determined by extensive 2D NMR as *ent*-15,19-dihydroxy-labda-8(17),13(E)-diene (**14**) and *ent*-3α,15-dihydroxy-labda-8(17),13(E)-diene (**15**) with oxidation occurring at the C19 and C3 positions, respectively (**Fig. 4C**) (**Supplementary Data 4**), coinciding with the hydroxylation positions of CYP701A64_2B and CYP99A39_2B on the respective KSL products. Such enzymatic promiscuity underscores the potential for engineering these enzymes to produce a wide array of novel diterpenoids. Additionally, minor production of **16** was detected when CYP701A64_2B was co-expressed with TaCPS-B2 and TaKSL-B3, and the product was putatively identified as (Z)-biformene-19-ol (**16**) based on the 19-hydroxylase catalytic activity of CYP701A64_2B on substrates **9** and **10**. Co-expression of CYP99A39_2B with TaCPS-B2 and TaKSL-B2 did not produce any detectable product, indicating that it may have more specific substrate preference (**Fig. 4C**). GC-MS mass spectra for pathway products and intermediates of BGC2(2B) is presented in **Supplementary Fig. 19.** BGC2(2B)-related genes and compounds are listed in **Supplementary Data 2** and **Supplementary Data 3**, respectively.

### BGC2 is interlinked with BGC1 and is syntenic with gene clusters in other grass genomes

Given that both the aspisoside and scutenol biosynthetic pathways are pathogen-induced and overlap in the tissues in which they are expressed, we hypothesized that the two pathways may potentially interact with each other. To test this, we co-expressed BGC2(2B) genes together with UGT705A6 or UGT705A8 in *N. benthamiana*. While UGT705A8 did not exhibit glycosylation activity in any of the gene combinations tested, co-infiltration of UGT705A6 together with scutenol-biosynthetic genes did indeed result in new glycosylation products of scutenol A and scutenol B, as well as of two pathway side-products, with accurate masses corresponding to the addition of a deoxyhexose (+146 Da), plausibly a rhamnosyl group. Specifically, the newly-formed compounds were identified as *ent*-pimara-8(14),15-diene-19-deoxyhexoside (**17**) (*m/z*=434.31), (Z)-biformene-3-deoxyhexoside (**18**) (*m/z*=434.31), *ent*-15-hydroxy-labda-8(17),13(E)-diene-19-deoxyhexoside (**19**) (*m/z*=452.31), and (Z)-biformene-19-deoxyhexoside (**20**) (*m/z*=434.31), based on their masses and the combination of co-infiltrated genes in each of the experiments (**Supplementary Fig. 20, Supplementary Data 3**). However, we could not detect any of these glycosides by LC-MS analysis of *M. oryzae*-infected plants, and thus the *in-planta* interplay between the pathways encoded by the two gene clusters remains to be further investigated.

Similarly to BGC1, microsynteny analysis allowed us to identify partially conserved gene clusters in additional grass species that are syntenic with BGC2, of which their metabolic products are either unknown or are chemically distinct from the wheat BGC2 products. Specifically, partially conserved clusters were found in barley, oat, Brachypodium spp., and rice (**Fig. 4E**). In the latter, the syntenic region is comprised of a tandem array of KSL genes that includes OsKS1 (kaurene synthase involved in gibberellin biosynthesis), OsKSL2 (bayerene synthase)^44^, and OsKSL3, which was defined as a pseudogene^23^. Gene phylogeny also shows that OsKSL1/2/3 group together with TaKSL5/8, which are found in the BGC2 cluster (**Supplementary Fig. 2**). Genes from the syntenic clusters in the remaining abovementioned species have not been functionally characterized to the best of our knowledge.

### Aspisosides and scutenols exhibit antifungal and antibacterial activity *in vitro*

Given the pathogen-responsive expression patterns of BGC1 and BGC2 genes and accumulation of their metabolic products following pathogen infection (*i.e*., a phytoalexin-like accumulation pattern), we aimed to evaluate the antifungal potential of aspisosides and scutenols against major fungal pathogens that threaten crop health and yields. We therefore examined their *in vitro* antifungal activity against two cereal fungal pathogens *Fusarium culmorum* and *Magnaporthe oryzae*, as well as against two generalist fungal plant pathogens, namely *Botrytis cinerea* and *Rhizoctonia solani*. Fungal growth inhibition was determined using the agar disc diffusion method as carried out previously^45^. In these assays, aspisoside A exhibited notably stronger fungal growth inhibition compared to aspisoside B, with the strongest inhibition observed against *M. oryzae*. Conversely, scutenol A and B showed little to no antifungal activity against any of the tested pathogens, indicating a limited role in direct suppression of fungal pathogens (**Fig. 5A**).

**Figure 5.**
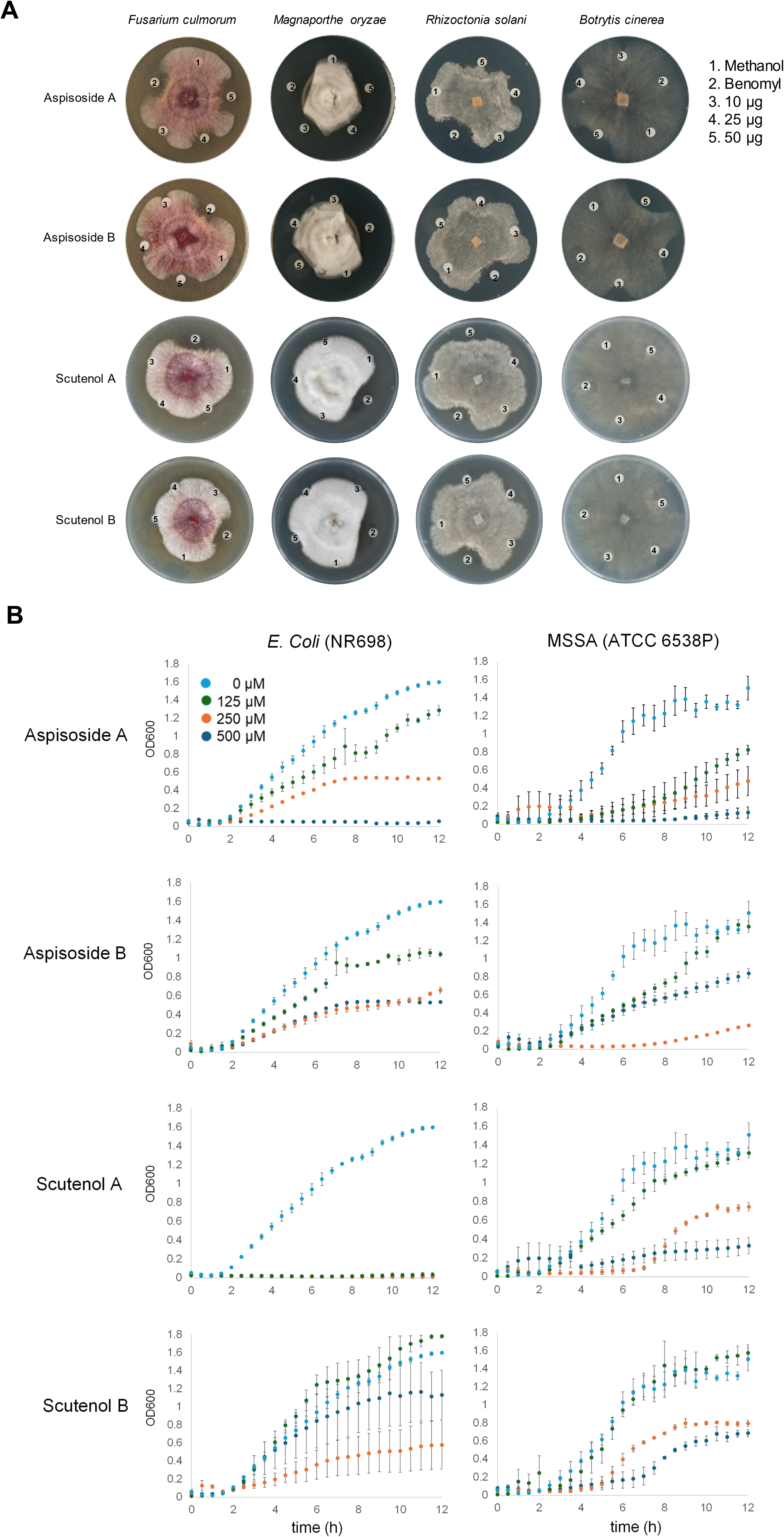
Antimicrobial activity of aspisosides and scutenols. **A**, fungal inhibition disc diffusion assays of purified compounds aspisoside A/B, and scutenol A/B, against four fungal plant pathogens. The fungicide benomyl was used as positive control. **B**, bacterial growth inhibition assays of methicillin-sensitive *Staphylococcus aureus* (MSSA) and *E. coli* strain NR698, in liquid LB media containing 125–500 µM aspisoside A or B. Mean values ±SEM are shown. Each time point respectively represents three biological replicates.

Publicly available wheat gene expression data shows that BGC1(2D) and BGC2(2B) are not only upregulated by fungal pathogens but also induced during bacterial pathogen infection and in response to flagellin, a bacterial PAMP (**Supplementary Figs. 12 and 15**). Therefore, we evaluated the potential antibacterial activity of aspisosides and scutenols using bacterial growth-inhibition assays in liquid media using 96-well plates, by monitoring OD₆₀₀ every 30 min over a 12-hour-long incubation. The assays included two bacterial species: the Gram-negative *Escherichia coli* (hyperpermeable mutant strain NR698 that is deficient in its outer membrane), and the Gram-positive *Staphylococcus aureus* (ATCC 6538P, methicillin-sensitive; MSSA). Complete growth inhibition was observed with *E. coli* NR698 in the presence of scutenol A in the lowest concentration tested (125 µM) whereas complete inhibition of growth was observed with both the *E. coli* and *S. aureus* strains in the presence of aspisoside A, albeit only at a high concentration of 500 µM. However, low or no inhibitory activity was observed either with scutenol B or aspisoside B (**Fig. 5B**). Taken together, the results suggest that both aspisosides and scutenols exhibit antifungal and/or antibacterial activity *in vitro*, thus supporting a defensive role for these compounds against microbial pathogens in wheat.

## DISCUSSION

A genome-wide search for diterpene synthases in the bread wheat genome, and analysis of their expression patterns under biotic stress conditions, have led us to identify several groups of pathogen-induced CPS and KSL genes, intriguingly all of which are co-localized in one of the two biosynthetic gene clusters in group 2 chromosomes. Combinatorial heterologous expression of BGC1 genes in *N. benthamiana* enabled us to reconstruct the biosynthetic pathway leading to production of two diterpene glycosides, aspisoside A and B. Similarly, heterologous expression assays also demonstrated the enzymatic activities of pathogen-induced genes located in BGC2 on Chr.2B. In contrast to BGC1, the genes within BGC2 encode two short and separate pathways for the *ent*-CPP derived diterpene alcohols, scutenol A and scutenol B. Promiscuous activity of the CYP99 and CYP701 genes from BGC2 also allowed the biosynthesis of three additional products in *N. benthamiana*, two of which are generated by direct activity on CPP rather than on a KSL product. Direct activity of cytochrome P450s on CPP is unusual in diterpene biosynthesis, but was also previously observed in switchgrass, where CYP71 enzymes were found to catalyze the formation of a furane ring on several different CPS products^43^.

BGCs that are syntenic to those encoding aspisoside and scutenol biosynthesis in wheat occur in several other grass species, but notably produce structurally distinct diterpenes, pointing to the exceptional chemical diversity of the cluster-encoded pathways within the Poaceae. Given the abundance of genomic data available for the grasses, and the wide occurrence of the momilactone-like BGCs (MLBGCs) within this family^46^, this cluster could represent an excellent model for studying chemical diversification in plant specialized metabolism. Functional characterization of MLBGCs in additional Poaceae species will likely provide additional intriguing examples of diversification of the clusters’ metabolic pathways through gene recruitment, gene loss, and enzyme neofunctionalization. An interesting feature in the diversification of the pathways encoded by the MLBGCs lies in their initial step, in which OsCPS4 from the momilactone cluster produces *syn*-CPP, whereas the CPS enzymes in wheat BGC1 and BGC2 respectively produce normal CPP and *ent*-CPP (**Fig. 6**). Divergent stereochemistry is thus already evident as a factor contributing to the diversity of compounds produced by the MLBGCs, alongside the different enzymatic activities of the KSL and downstream decorating genes. Stereo-divergence in biosynthesis of plant specialized metabolites was similarly observed in the case of steroidal glycoalkaloids in the Solanaceae family, the result of distinct activities of a divergent cytochrome P450 enzyme^47^. For BGC2, syntenic clusters can be found in several other grass genomes, including *B. distachyon*, oat, and barley (**Fig. 4E**), but the wheat BGC2 is the first of these clusters to be functionally characterized. Nevertheless, some diversification is inherent in the cluster itself, as it encodes two short separate pathways to scutenol A and scutenol B, initially derived from different modifications of *ent*-CPP by TaKSL-B2 and TaKSL-B3, which respectively generate *ent*-pimara-8(14),15-diene, and (Z)-biformene.

**Figure 6.**
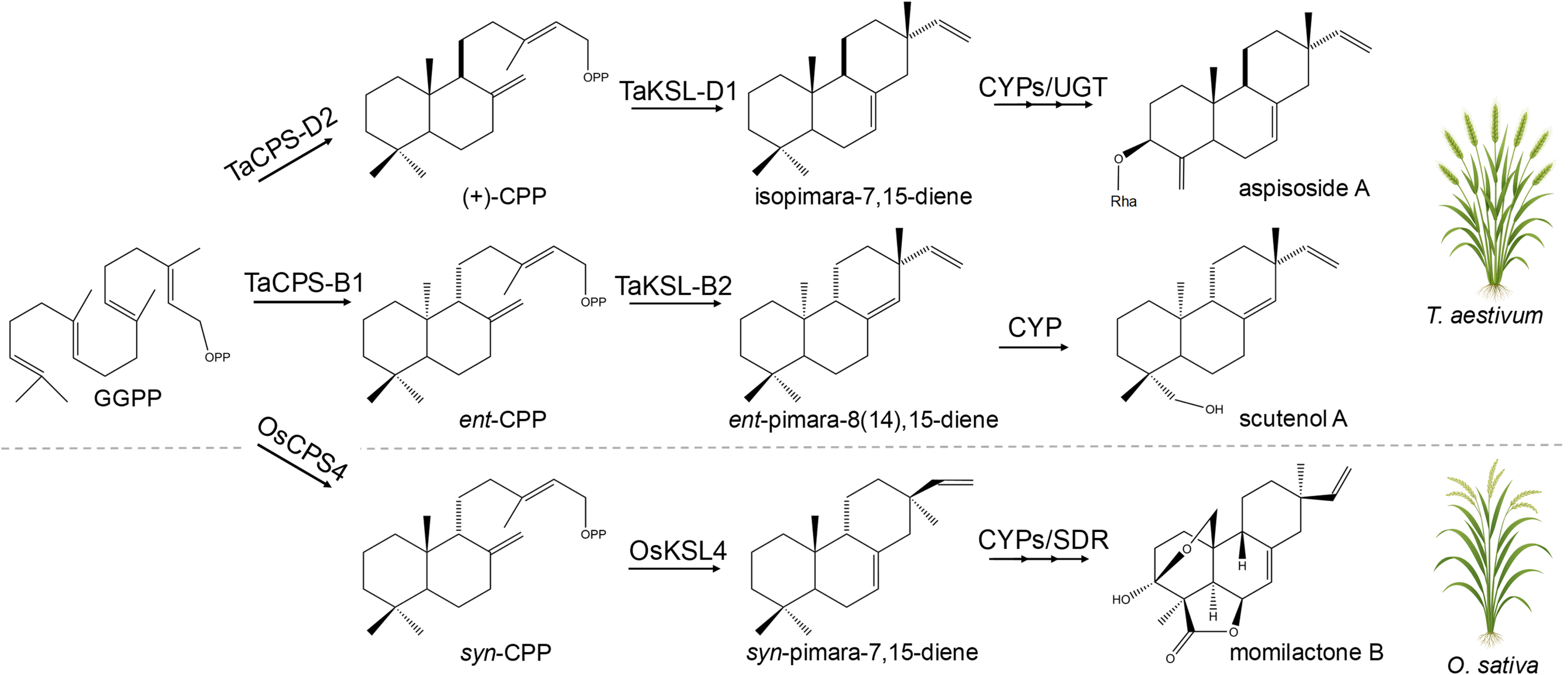
Stereo-divergence underlies chemical diversification of diterpenoid pathways encoded by the momilactone-like gene clusters in rice and wheat. Divergence of the three pathways to aspisosides, scutenols, and momilicatones is already evident in their respective first steps, in which OsCPS4 from the momilactone cluster produces *syn*-CPP, whereas the CPS enzymes in wheat BGC1 and BGC2 respectively produce normal CPP and *ent*-CPP.

The evolutionary history of these clusters within the grasses also remains enigmatic. The occurrence of homologous MLBGCs in species of the relatively distant Pooideae, Oryzoideae, Chloridoideae and Panicoideae groups was proposed to be the result of two lateral gene transfer events, first from the Pooideae to the Chloridoideae/Panicoideae group, and from Chloridoideae/Panicoideae to the Oryzoideae in a subsequent event^46^. However, this explanation is unlikely when considering the syntenic location of the wheat/barley BGCs and the rice momilactone cluster (**Fig. 1F**), as it would require the lateral gene transfer to ‘return’ the gene cluster to its original genomic location. Lateral gene transfer is indeed widespread among grasses, but tends to occur between species from closely related groups and typically to non-syntenic genomic regions^48,49^. In the future, a wider analysis of the presence/absence of MLBGCs across the Poaceae may furnish a better understanding of the evolutionary history of these clusters^40^.

Glycosylated diterpenoids in plants are relatively scarce (compared to triterpene glycosides, for example), and knowledge about their biosynthesis remains limited^50,51^. In the grass family, the occurrence of putative diterpene glycosides has been reported in barley ^12^ and switchgrass^52^, but the structures, biosynthesis, and function of these compounds are unknown. Our identification of structurally defined diterpene glycosides in wheat therefore provides the first direct chemical and biosynthetic evidence that diterpene glycosylation constitutes a *bona fide* branch of phytoalexin metabolism in grasses. The occurrence of aspisosides thus resolves earlier reports of putative diterpene glycosides in cereals by linking defined chemical structures to a genetically encoded biosynthetic pathway, and suggests that diterpene glycosylation may be more widespread in the Poaceae than previously appreciated.

The unusual activity of diterpene glycosylation catalyzed by UGT705A6 and UGT705A8 is further compounded by their preference for the uncommon sugar donors UDP-rhamnose and UDP-arabinose, respectively. The use of these sugar donors distinguishes aspisosides from the more prevalent glucose-conjugated plant metabolites and may have functional implications for diterpene phytoalexin stability, localization or transport. Phylogenetic analysis placed the aspisoside UGTs in the 705 clade, a monocot-specific subgroup of group D glycosyltransferases. The functions of only three genes from this clade are known so far, namely *OsUGT705A1*, *OsUGT705A2,* and *OsUGT705A3* from rice, which interestingly glycosylate flavonoids rather than terpenes^35^. The recruitment of UGT705 enzymes for diterpene glycosylation further highlights glycosyltransferases as potential evolutionary innovation points in grass specialized metabolism. The apparent functional divergence of UGT705 family members, from flavonoid glycosylation in rice to diterpene glycosylation in wheat, suggests that this clade retains catalytic plasticity that can be co-opted for structurally distinct classes of metabolites.

The *in vitro* antifungal and/or antibacterial activities exhibited by aspisosides and scutenols and the strong induction of their biosynthetic genes following treatment with pathogens or PAMPs, are consistent with a role for these compounds in wheat defense. How these compounds specifically contribute to plant defense (*e.g*., against which specific pathogens, in which developmental stages etc.) remains to be determined, and will require additional future investigations, possibly through pathogenicity assays with pathway mutants or with transgenic lines overproducing aspisosides or scutenols. Further to obtaining a deeper understanding of the potential defensive roles of aspisosides and scutenols *in planta*, additional investigations of these and other diterpenoids will also have to take into consideration the plausible accumulation of additional diterpenoids produced in hexaploid wheat in response to pests or pathogens, which may be generated by distinct biosynthetic pathways or ones diverging from those we describe here. Overall, considering wheat polyploidy and the extensive set of diterpene synthase candidates identified in its genome, the complexity of its diterpene metabolic network may even exceed those found in rice and maize. The findings presented here therefore likely provide only a partial glimpse of the entire wheat diterpene metabolic network.

## MATERIALS AND METHODS

### Microsynteny analysis

To perform microsynteny analysis and generate figures, a python implementation of MCScan was used (https://github.com/tanghaibao/jcvi/wiki/MCscan)^53^. FASTA and GFF3 files were retrieved from EnsemblPlants (http://plants.ensembl.org) for chromosomes 5A, 5B and 5D of *Triticum aestivum* (IWGSC), 5A and 5D of *Triticum turgidum subsp. diccocoides* (WEWSeq_v.1.0) and 5D of *Aegilops tauschii* (Aet_v4.0); from NCBI (https://www.ncbi.nlm.nih.gov/datasets/genome/) for chromosome 4 of *Oryza sativa japonica* group (IRGSP-1.0); from Phytozome (http://phytozome.jgi.doe.gov/) for chromsome 2H *Hordeum vulgare* (r1), 5 of *Brachypodium distachyon* (v3.2) and 9 of *Brachypodium stacei* (v1.1); and from (http://www.ncgr.ac.cn/oat) for chromsome 4 of *Avena strigosa*. MCScan ortholog finding and synteny assignment was run with a c-score of 0.99 and a single iteration.

### Gene cloning and synthesis

RNA from 10-day-old *Triticum aestivum* plants (Chinese Spring), infected with powdery mildew (*Blumeria graminis f. sp. tritici*), was isolated using RNeasy plant mini kit (Qiagen) according to the manufacturer’s instructions. cDNA was generated using the Super Script III First-Strand Synthesis kit (Invitrogen) using oligo(dT) primers, according to manufacturer’s protocols. Gene coding sequences of CYP99A8_2D (TraesCS2D02G029700), CYP99A10_2D (TraesCS2D02G029900), CYP99A7_2D (TraesCS2D02G030000), UGT705A6 (TraesCS2D02G029800), UGT705A9 (TraesCS2B01G040500), UGT705A10 (TraesCS2B01G040600), TaCPS-B1 (TraesCS2B02G445500; Genbank PV805598), TaKSL-B2 (TraesCS2B02G445200; PV805597), CYP701A64_2B (TraesCS2B02G445300), and CYP99A39_2B (TraesCS2B02G445600) were amplified by PCR using Phusion DNA polymerase (Thermo Fisher Scientific) or Q5 DNA polymerase (New England Biolabs) and gene specific primers from Merck Life Science (**Supplementary Table S2**). tHMGR was previously amplified from *Avena strigosa* cDNA^29^. TaCPS-B6 (TraesCS2B02G040400; PV805599) was cloned into a 3α1 plant expression vector, driven by *Solanum lycopersicum* Ubiquitin 10 promoter and terminator, using Goldenbraid cloning^54^. TaCPS-D2 (TraesCS2D02G029600), TaKSL-D1 (TraesCS2D02G030100), UGT705A8 (TraesCS2A02G028000), TaKSL-B3 (TraesCS2B02G445100; PV805596), and TcGGPPS (GenBank AF081514) were synthesized by Twist Bioscience, San Francisco, CA, USA. TaCPS-D2, TaKSL-D1, TaCPS-B1, TaKSL-B2 and TaKSL-B3 sequences were cloned with an N-terminal truncation for cytosolic expression. Synthesized and PCR-amplified genes were cloned into a pDONR207 Gateway entry vector and subcloned into a pEAQ-HT-DEST1 plasmid, using BP and LR Clonase II enzyme mixes (Thermo Fisher Scientific), respectively. Plasmid DNA was isolated using the QIAprep Spin Miniprep Kit (Qiagen). Sequence confirmation was carried out using Sanger DNA sequencing (Eurofins Genomics).

### Transient expression in *Nicotiana benthamiana via* agrobacterium-mediated infiltration

pEAQ-HT-DEST1 plasmids containing a single gene of interest were transformed into electrocompetent *Agrobacterium tumefaciens* cells (GV3101) *via* electroporation. Transformed cells were grown on LB-agar plates containing kanamycin (50 μg/mL) and rifampicin (25 μg/mL) at 28°C for 2-3 days. Single colonies were then picked for growth at 28°C in 10 mL LB media with antibiotics selection. Cultures grown overnight were centrifuged at 5,000 g for 5 min, the supernatant discarded, and the pellet resuspended in MMA buffer (10 mM MgCl_2_, 10 mM MES pH 5.6, 200 µM acetosyringone) to OD_600_ 0.2 and incubated for 2-3 h at room temperature. For small-scale transient experiments, leaves of 5-weeks-old greenhouse-grown *Nicotiana benthamiana* plants were infiltrated using a needleless syringe. Leaves for control experiments were infiltrated with agrobacteria harboring an empty pEAQ-HT-DEST1 vector. The plants were further maintained in the greenhouse after infiltration. Infiltrated leaves were harvested 5 days post infiltration (dpi) and freeze-dried prior to metabolite extraction.

### Wheat infection with fungal pathogens

Methods for infection of ‘Chinese Spring’ wheat plants with *Fusarium culmorum* were as previously reported^39^. Infection with cereal blast (*Magnaporthe oryzae*) was performed as follows. A fungal inoculum of *M. oryzae* strain BTJ4P^55^ was prepared as previously described^56^. A conidial suspension of 0.15 × 10^6^·ml^-1^ conidia was used for spray-inoculation of ten 3-week-old wheat plants (cv. Cadenza). Ten mock-treated plants were sprayed with water only. Following inoculation, plants were grown in a growth chamber (22°C, 16 h light, 8 h dark) and harvested 4- and 10-days post inoculation. 5 biological replicates of mock- or blast-infected plants were individually collected, each replicate comprised of 10 plants. Samples were freeze dried immediately after collection.

### Large-scale *N. benthamiana* agroinfiltration, extraction, purification, and NMR analyses

For large-scale *N. benthamiana* infiltration experiments, whole plants were infiltrated under vacuum, as previously described^16,29,32^. Specific methods for metabolite extraction, purification, and NMR analyses of the purified compounds are detailed in **Supplementary Data 4**.

### Metabolite extraction from *Nicotiana benthamiana* leaves

Freeze-dried *Nicotiana benthamiana* leaf tissue 5-days post agroinfiltration was collected and homogenized using two 3 mm tungsten carbide beads (Qiagen) in a TissueLyser (Geno/Grinder (Spex) at 1,200 rpm for 1 min at room temperature. Each biological replicate consisted of 3 leaves from the same experiment. For metabolite extraction, 500 μL ethyl acetate (HPLC grade; Fisher Scientific) was added to each sample (10 mg) and vortexed. The samples were further incubated for 1 h at room temperature (25°C) with intermediate shaking in a thermal block, followed by centrifugation at 18,200 *g* for 10 min for removal of cell debris. For GC-MS analysis, 100 μL supernatants were filtered on a mini column (pore size 0.22 µm, Geneflow) and transferred to 100 μL glass inserts placed in 2 mL vials. For LC-MS analysis, 200 μL of the supernatants were transferred to a new microcentrifuge tube and evaporated to dryness under N_2_. The samples were reconstituted in 50 μL methanol (HPLC grade; Fisher Scientific), filtered using mini columns (pore size 0.22 µm, Geneflow) and transferred to LC-MS vials.

### Metabolite extraction from fungus-infected wheat plants

Metabolite extraction of *F. culmorum* and *M. oryzae* infected and non-infected plants was performed similarly as described in^39^. In brief, 200 mg freeze-dried blade and sheath samples of treated (infected) and mock (non-infected) plants were extracted in 100% HPLC-grade ethyl acetate (4 mL) for 1 hr at 25°C with continuous shaking, containing 80 µg mL^−1^ digitoxin (Sigma Aldrich) as internal standard and centrifuged at 23,000 g for 10 min to remove cell debris. The supernatant was transferred to new microcentrifuge tubes and evaporated in a SpeedVac (Genevac EZ-2 Elite) at room temperature until complete dryness. Dried samples were reconstituted in 100% HPLC-grade methanol (500 µL) and filtered into LC-MS vials through a mini column (pore size 0.22 µm, Geneflow) for HPLC-MS analysis using a Thermo Scientific Q Exactive Hybrid Quadrupole-Orbitrap Mass spectrometer HPLC system. The samples were analyzed from five biological replicates, and each analyzed in three technical replicates.

### Metabolite extraction from silenced wheat tissues

At 21 days post-silencing (dps), mock (mcs4D) and silenced (PDS, TaCPS-D2, or TaKSL-D1) blade and sheath tissues were harvested, freeze-dried and grounded individually into powder with an IKA A11 hand grinder. 200 mg of blade and 150 mg of sheath tissues were extracted with 4 mL of 100% HPLC-grade ethyl acetate in 15 mL falcon tubes, containing 80 µg·mL^−1^ digitoxin (Sigma Aldrich) as internal standard. The mixture was vortexed for 1 h at 25°C and centrifuged at 4000 rpm for 15 min and the supernatant was then transferred into a new falcon tube (15 mL). The solvent was evaporated in a Genevac EZ-2 Elite centrifugal evaporator (SP). Finally, the dried samples were re-suspended in 100% HPLC-grade methanol (300 µL), filtered through a 0.22 μm filter membrane (Geneflow) and transferred to LC-MS vials for HPLC-MS/MS analysis using a Thermo Scientific Q Exactive Hybrid Quadrupole-Orbitrap Mass spectrometer HPLC system.

### GC–MS analysis of diterpenoids

Samples were analyzed using an Agilent 5977B MSD mass spectrometer (MS) connected to an Agilent 7890B gas chromatograph (GC) system equipped with a Zebron ZB-5HT (35 m length, 0.25 mm internal diameter, 0.10 μm film thickness) Inferno column (Phenomenex). One microliter sample was injected in pulsed splitless mode (16.038 psi pulse pressure) with the inlet temperature set to 250°C and the carrier gas (helium) with a flow rate of 1.2 mL/min. The GC oven temperature program was initially set at 130°C for 2 min, with subsequent increase from 130 to 250°C (8°C min^-1^), maintained for 5 min, then elevated from 250 to 310°C (10°C min^-1^) and maintained at 310°C for 3 min. A modified method was used to analyze TaKSL-B3 and downstream diterpene pathway products (**13**, **16**). In short, one microliter sample was injected in split mode (10:1; 9.3 psi pulse pressure) with the inlet temperature set to 280°C and the carrier gas (helium) with a flow rate of 1.2 mL min^-1^. The GC oven temperature program was initially set at 60°C for 2 min, with subsequent increase from 60 to 150°C (10°C min^-1^), then elevated from 150 to 250°C (20°C min^-1^) and maintained for 5 min. A full EI–MS was generated for each sample by scanning within the *m/z* range of 60 to 800 amu with a solvent delay of 15 min. For quantification, the detector was set to 7.2 scans/sec. GC-MS data was analyzed using Agilent MassHunter software.

### LC–MS analysis of diterpenoids

High-resolution mass spectrometry analysis of the metabolites was carried out on a Q Exactive instrument (Thermo Scientific). Chromatography was performed using a Kinetex 2.6 μm XB-C18 100 Å, 50 mm × 2.1 mm (Phenomenex) column kept at 30°C. Water containing 0.1% formic acid (FA) and acetonitrile containing 0.1% formic acid (FA) were used as mobile phases A and B, respectively, with a flow rate of 0.5 mL/min. A gradient elution program was applied as follows: 0–0.75 min linearly increased from 0 to 20% B, 0.75–7 min linearly increased from 20 to 60% B, 7–11.5 min linearly increased from 60 to 80% B, 11.5–15 min linearly increased from 80 to 100% B, 15– 15.5 min linearly decreased from 100 to 20% B hold for 1.5 min for re-equilibration, giving a total run time of 17 min. MS detection was performed following atmospheric pressure chemical ionization (APCI) in positive mode, in the range of 100-1500 *m/z* and a mass resolution of 70,000. Full scan in combination with data-dependent MS2 scans (Full MS/dd-MS2, Top 3) was applied. Targeted selected ion monitoring (tSIM) and parallel reaction monitoring (PRM) acquisition were performed with mass resolution set at 35,000 FWHM and a mass isolation window of 1.6 m/z and AGC of 3e6. In PRM mode, the data were acquired according to a predetermined inclusion list containing the accurate masses. All data were acquired and processed utilizing Thermo Scientific Xcalibur software.

### Relative quantification of aspisoside A and aspisoside B

Semi-quantitative LC-MS analyses of aspisosides A and B in wheat extracts were carried out using the Xcalibur software package v4.3 (Thermo Scientific). Automatic peak detection and integration was done with ICIS algorithm, applying 7 smoothing points and other default parameters, using [M + H]^+^ ion *m/z* = 419.2788 for aspisoside A, ion *m/z* = 255.2105 for aspisoside B, and ion *m/z*=375.2529 for the internal standard, digitoxin.

### RNA isolation and qRT-PCR

Total RNA was isolated from blade and sheath tissues using a RNeasy kit (Qiagen) with on-column DNAse treatment using a RNase-free DNAse set (Qiagen), according to the manufacturer’s instructions. Two μg of RNA was used to synthesize cDNA by using a High-Capacity cDNA Reverse Transcription Kit (Applied Biosystems) in a single 20 μL reaction, followed by dilution of the cDNA (1:4). qRT-PCR was performed on a CFX96 Touch Real-Time PCR instrument (Bio-Rad) in the following conditions: initial step in the thermal cycler for 3 min at 95°C, followed by PCR amplification for 40 cycles of 10 s at 95°C and 30 s at 59°C, and finally dissociation analysis to confirm the specificity of PCR products with 0.5°C ramping from 55°C to 95°C. Each 10 μL reaction was comprised of 5 μL LightCycler 480 SYBR Green I Master mix (Roche Life Science), 1 μL cDNA template, 3 μL H_2_O and 1 μL primer mix (0.5 μM each primer). Relative transcript levels were calculated according to the Pfaffl method^57^, using the housekeeping gene β-tubulin (TUBB) as reference. Three technical replicates were performed per sample, and 4 to 5 biological replicates were included for each condition. Primers were designed using Primer3 software^58^ and are listed in **Supplementary Table S2**.

### Virus-induced gene silencing in wheat

To silence the TraesCS2D02G029600 (TaCPS-D2) gene and TraesCS2D02G030100 (TaKSL-D1) gene, 225-bp and 249-bp fragments of cDNA sequence were respectively designed using si-Fi siRNA Finder software^59^ (**Supplementary Table S2**). The target sequences were PCR amplified including LIC adaptor sequences to the 5′-ends of the PCR primers, 5′-AAGGAAGTTTAA-3′ (forward primer) and 5′-AACCACCACCACCGT-3′ (reverse primer) and cloned into the BSMV RNAγ vector pCa-γbLIC^37^ *via* ligation independent cloning (LIC), in antisense orientation. The pCa- γbLIC vector was linearized with ApaI (New England Biolabs) at 25°C for 2 h, followed by inactivation of the enzyme by incubating at 65°C for 20 min. To generate complimentary sticky ends, both the gel-purified PCR products and linearized pCa-γbLIC vector were treated with T4 DNA polymerase. In brief, ∼200–250 ng of the PCR products were incubated at 22°C for 30 min in a total volume of 10 μL with 0.6 U of T4 DNA polymerase (New England Biolabs) and 500 ng of ApaI-digested pCa-γbLIC was incubated at 22 °C for 30 min in a total volume of 50 μL with 3 units of T4 DNA polymerase (New England Biolabs), followed by heat-inactivation of the T4 DNA polymerase by incubation at 75°C for 15 min. Further, 10 μL of treated PCR products (∼200–250 ng) and 2 μL of the treated pCa-γbLIC vector (20 ng) were mixed, incubated at 65°C for 2 min and then incubated for 10 min at room temperature to allow annealing of the complementary sticky ends. 2 μL of the reaction mixture was used to transform subcloning efficiency™ DH5α competent cells (Invitrogen), and recombinant cells were selected on kanamycin plates (50 µg/mL). The transformants were confirmed by sequencing the plasmid DNA with primers 2235.F (5′-GATCAACTGCCAATCGTGAGTA-3′) and 2615.R (5′- CCAATTCAGGCATCGTTTTC-3′) that flank the LIC site in the pCa-γbLIC vector.

Sequence verified recombinant pCa-γbLIC VIGS constructs and other binary vectors pCaBS-α, pCaBS-β were transformed into *Agrobacterium tumefaciens* GV3101 by electroporation. To prepare the virus inoculum, the BSMV vectors were first introduced *via* agroinfiltration into the leaves *Nicotiana benthamiana*. For agroinfiltration, Agrobacteria cultures were grown overnight in 28°C in LB media, resuspended in MMA buffer (10 mM MgCl2, 10 mM MES pH 5.6, 200 µM acetosyringone) to OD_600_ 1.5, and incubated at room temperature for 2–3 h. The agrobacteria strains carrying BSMV binary vectors pCaBS-α, pCaBS-β, and pCa-γbLIC constructs were mixed in a 1:1:1 ratio and infiltrated into fully expanded leaves of approximately 5-6 week old *N. benthamiana* plants using a needleless 1 ml syringe. A pCa-γbLIC construct targeting the wheat TaPDS gene was used a as positive control. Infiltrated *N. benthamiana* leaf tissue at 3–4 days post-agroinfiltration was harvested and grinded using a prechilled mortar and pestle in 10 mM potassium phosphate buffer pH 7 containing 1.5% w/v Celite 545 AW (Sigma-Aldrich) abrasive until a fully homogenous slurry was produced. To infect wheat plants with BSMV, one or the two consecutive youngest fully expanded leaves were firmly rubbed 3–5 times with the slurry. 5–10 min post-infection the leaves were sprayed using a fine mist of water and the plants were covered with plastic bags overnight. On the next day the plants were returned to standard growth conditions; photoperiod: 16 h, temperature: 22-25°C and relative humidity: 50-60%. Two-week-old treated plants were sprayed with a 200 µM methyl jasmonate (MeJA) + 0.01% Tween-20 solution. Four days after spraying with MeJA, wheat blade and sheath tissues were collected and stored in −80°C for further experimental analysis.

### Mycelial growth inhibition *in vitro* assays

The agar disc diffusion method^45^ was used to assay the pure compounds (aspisoside A, aspisoside B, scutenol A and scutenol B) as mycelial growth inhibitors. Mycelial discs (Ø 5 mm) of the fungi from the edges of a 6–10-day old culture were centrally placed in Petri dishes (Ø 90 × 25 mm), each containing 25 mL Potato Dextrose Agar (PDA) media and Complete Medium Agar (CMA) media (three plates per treatment and concentration, with three repeats). Next, sterile Whatman filter paper (No. 42) discs of Ø 5 mm were evenly placed onto the plates at equal distance from the advancing edge of the growing fungal mycelia for two days (*Fusarium culmorum*, *Botrytis cinerea*, and *Rhizoctonia solani*) or five days (*Magnaporthe oryzae*). The pure compounds, in methanol solution, were added to the Whatman filter paper discs. Methanol was added to an additional disc as negative control, and benomyl (0.1 mg/mL) in methanol was used as positive control. Plates were incubated at 22–23°C for three days (*Fusarium culmorum*, *Botrytis cinerea*, and *Rhizoctonia solani*) or seven days (*Magnaporthe oryzae*) and photographed.

### Antimicrobial *in vitro* activity assays

Bacterial growth inhibition effects of purified compounds aspisoside A, aspisoside B, scutenol A and scutenol B were assessed as described previously^39^. In brief, the bacterial strains *Escherichia coli* (NR698), *Bacillus subtilis* (168), and *Staphylococcus aureus* (ATCC 6538 P, methicillin-sensitive; MSSA) were initially grown in fresh liquid LB media (5 mL) at 37°C with 200 rpm shaking overnight, and diluted the next day to OD_600_ 0.001. 200 μL cultures were prepared in 96-well plates (Sterilin microtiter U well plates, Thermo Scientific). Each well contained 196 μL bacterial suspension in LB media with initial OD_600_ of 0.001, and 4 μL of each compound stock solutions in DMSO. Control wells contained 4 μL DMSO without any compound. 75 μL mineral oil (Aldrich) was added to each well to prevent evaporation. Each plate contained three or four replicate wells. The bacteria were grown at 37°C for 12 h in a FLUOstar Omega Microplate Reader (BMG Labtech), under constant shaking at 500 rpm, with OD_600_ readings taken every 30 min.

## AUTHOR CONTRIBUTIONS

G.P., R.C.M., and A.O. conceived and designed the project. G.P., R.C.M., and O.T. conducted gene cloning and vector construction, expressed heterologous genes in *N. benthamiana*, extracted and quantified metabolites from wheat and *N. benthamiana*, and performed analyses using GC-MS and LC-MS/MS. G.P. and R.C.M. carried out virus-induced gene silencing (VIGS) in wheat, conducted quantitative real-time PCR, and analyzed all experimental results. G.P., R.C.M. and B.S. designed and performed bacterial growth inhibition assays. G.P. and R.C.M. designed the *in vitro* mycelial growth inhibition assays; R.C.M. performed the experiment and analyzed the data. G.P. and A.S. performed infection experiments of wheat plants with *Fusarium* and blast fungal pathogens. R.C.M. carried out large-scale agroinfiltrations in *N. benthamiana* and collected the leaf material. A.E.D. performed compound purifications, NMR analyses and structural assignments. G.P., A.R., and C.O. analyzed genomic data and performed bioinformatic analyses. G.P., B.W., P.N., and A.O. supervised research. G.P., R.C.M., and A.O. wrote the manuscript, with contributions from all other co-authors.

## ACKNOWLEDGEMENTS

G.P. lab is supported by an Israel Science Foundation grant (315/24). A.O. acknowledges the support of the BBSRC Institute Strategic Programme Grant ‘Harnessing biosynthesis for sustainable food and health’ (BB/X01097X/1) and the John Innes Foundation. We thank JIC Horticultural Services for assistance with plant cultivation, and Dr. Lionel Hill and Dr. Paul Brett from the JIC Metabolomics platforms for assistance with metabolic analysis. We thank Prof. Diane Saunders for providing the BSMV vectors, Dr. Alba Pacheco-Moreno for providing a benomyl standard and *Fusarium* and *Rhizoctonia* strains, Prof. Michael H. Court for naming UGT genes and Prof. David R. Nelson for naming cytochrome P450 genes.

## Notes

### Competing Interest Statement

The authors have declared no competing interest.

