## Supplementary Information for "Diverse Diterpenoid Phytoalexins Shape Wheat Chemical Defenses"

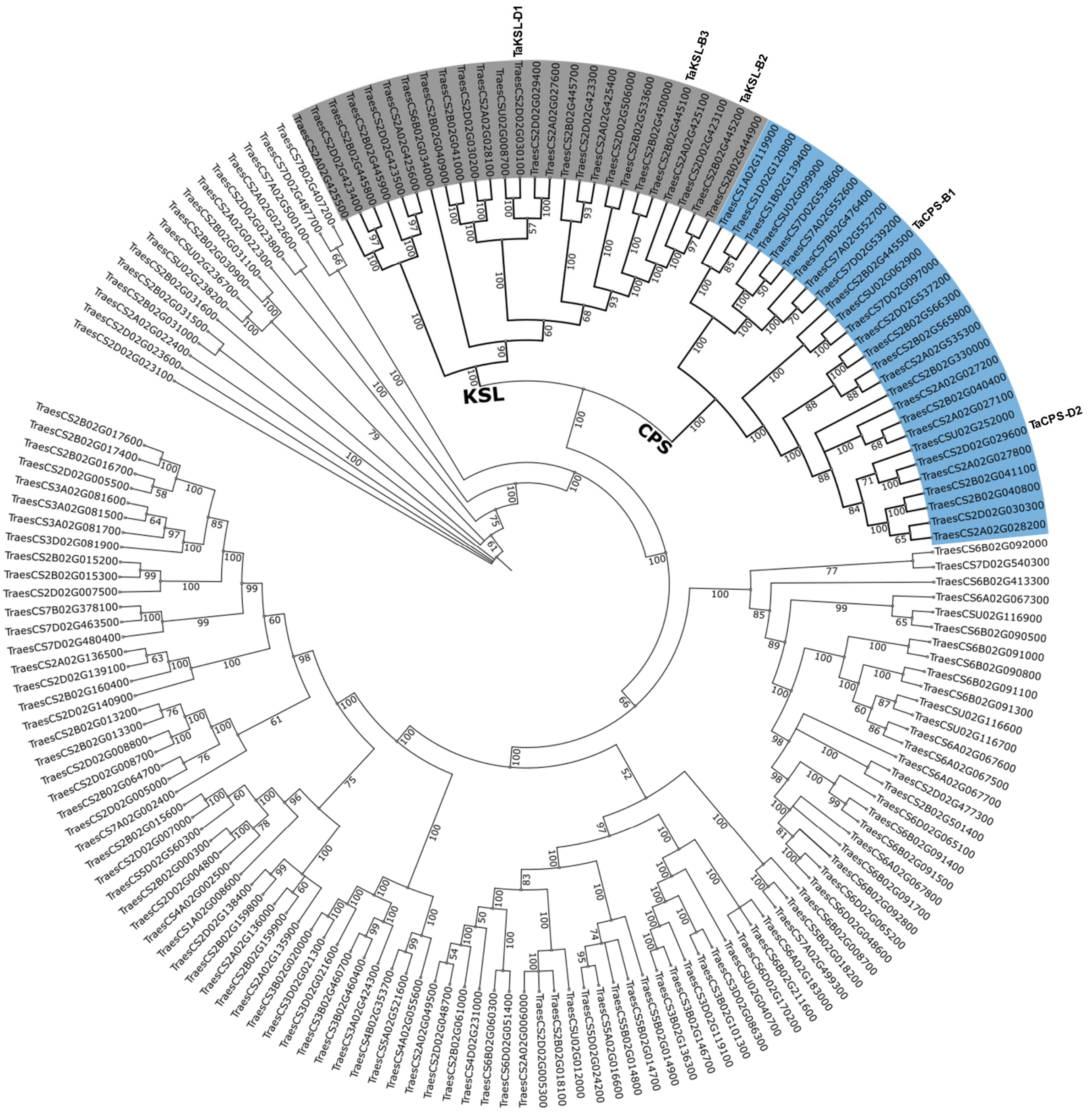


**Supplementary Figure 1. Maximum likelihood tree of wheat terpene synthases.** Wheat proteins containing the ‘terpene synthase, N-terminal’ domain (pfam domain pf01397) were retrieved on Interpro from the UniProtKB database[1]. 213 matching IWGSC v1.1 protein accessions were subsequently obtained from Ensembl Plants (https://plants.ensembl.org). Three sequences with <100 amino-acid-length were omitted along with 42 redundant protein sequences (keeping a single, longest predicted protein per gene ID). The remaining 168 protein sequences were aligned in MEGA11[2] using the MUSCLE algorithm, and a maximum likelihood tree was constructed using default parameters and 500 bootstraps. Decorative adjustments for visualization were done using iTOL v7[3]. CPS branch is highlighted in blue, KSL branch is highlighted in gray.

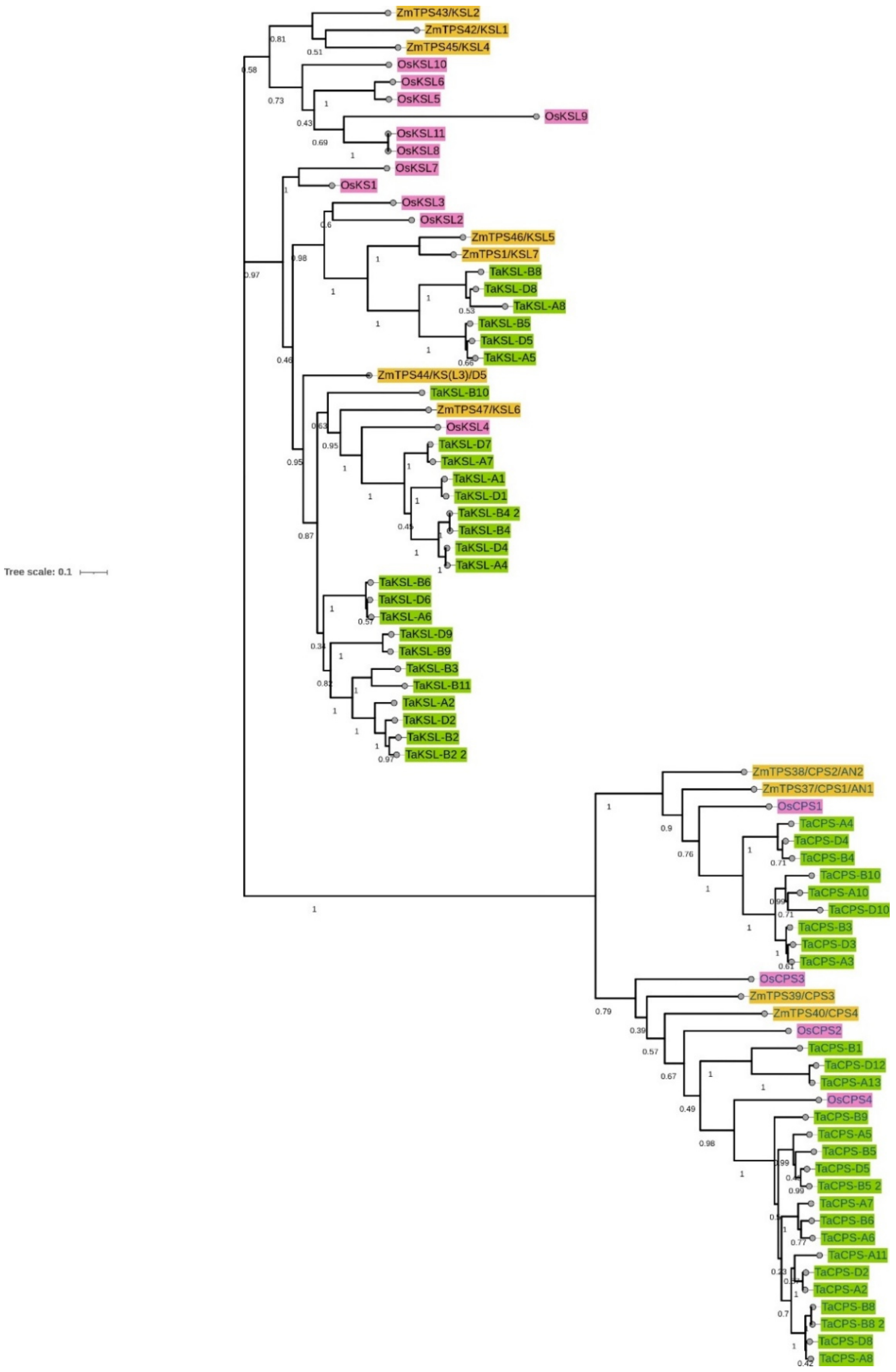


**Supplementary Figure 2. Maximum likelihood tree of wheat diterpene synthases.** 53 diTPS sequences identified in wheat (green) were used to construct a maximum likelihood tree together with known diTPS sequences from maize[4] (orange), and rice[5] (pink), all retrieved from Ensembl Plants (https://plants.ensembl.org). The sequences were aligned in MEGA11[2] using the MUSCLE algorithm, and a maximum likelihood tree was constructed using default parameters and 100 bootstraps.

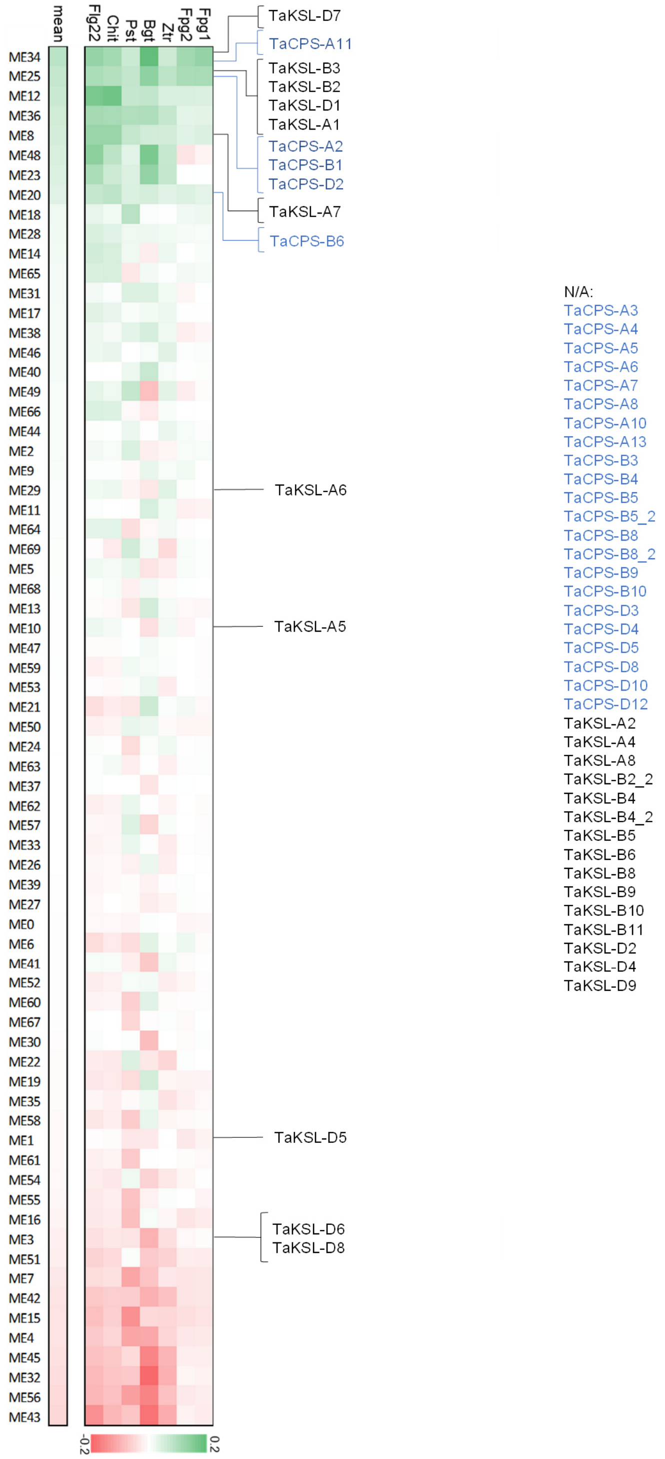


**Supplementary Figure 3.** Assignment of identified wheat *CPS* and *KSL* genes to gene expression modules of a biotic stress related WGCNA[6,7]. 69 gene expression modules are ordered from the most highly upregulated (top) to the most highly downregulated (bottom), following pathogen infection or treatment with chitin or flagellin. All identified *CPS* and *KSL* genes were assigned to the different modules. N/A denotes unassigned (non-expressed) genes. Fpg1, *Fusarium pseudograminearum*; Fpg2, *Fusarium pseudograminearum*; Ztr, *Zymoseptoria tritici*; Bgt, *Blumeria graminis f. sp. tritici*; Pst, *Puccinia striiformis f. sp. Tritici*; Chit, chitin; Flg22, flagellin peptide.

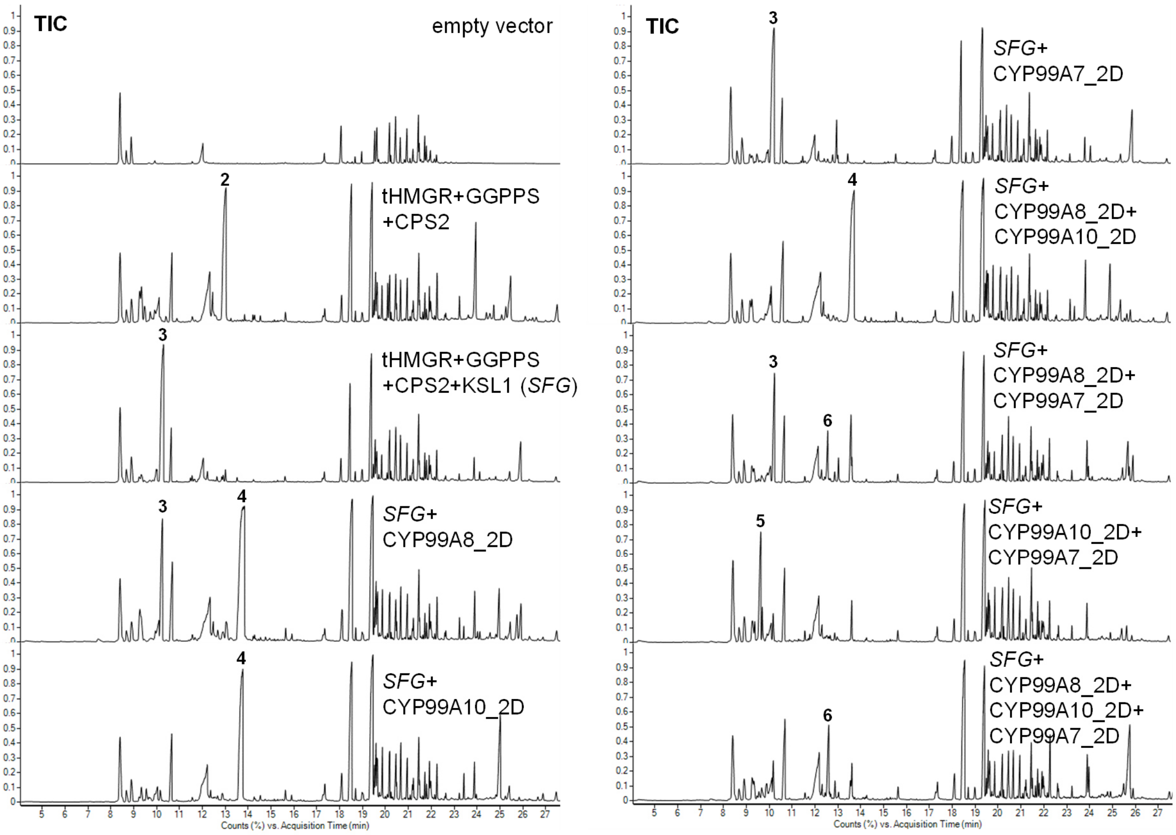


**Supplementary Figure 4. GC-MS total ion chromatograms (TIC) of combinatorial transient expression experiments of BGC1(2D) genes in *Nicotiana benthamiana*.** SFG, scaffold forming genes (tHMGR+TcGGPPS+TaCPS2+TaKSL1). (+)-copalol (**2**), isopimara-7,15-diene (**3**), 7,15-isopimaradien-18-ol (**4**), 18-norisopimara-4(19),7,15-triene (**5**), 3-hydroxy-18-norisopimara-4(19),7-15-triene (**6**).

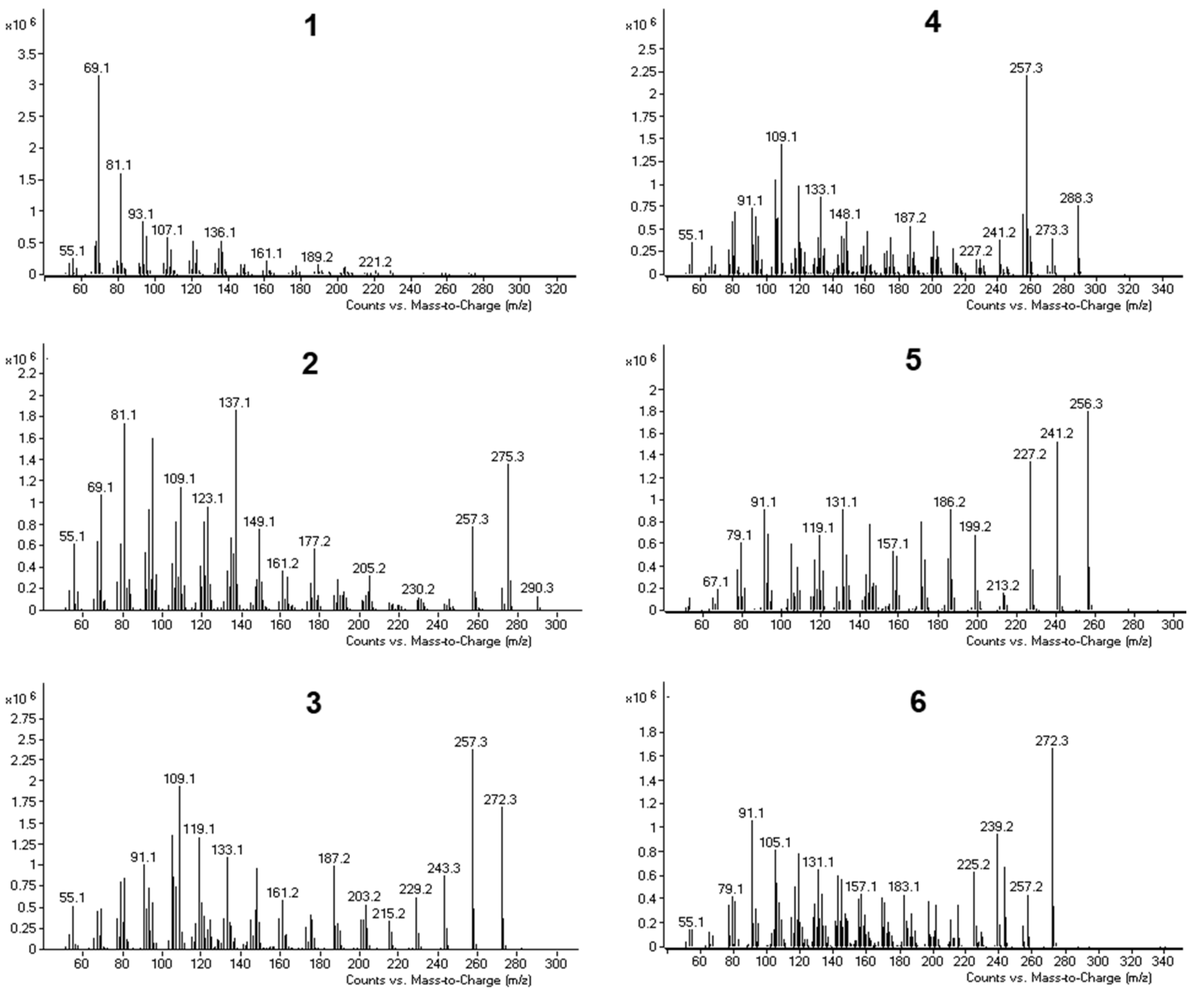


**Supplementary Figure 5. GC-MS mass spectra of BGC1(2D)-encoded pathway products and intermediates.** GGPP **(1)** (+)-copalol (**2**), isopimara-7,15-diene (**3**), 7,15-isopimaradien-18-ol (**4**), 18-norisopimara-4(19),7,15-triene (**5**), 3-hydroxy-18-norisopimara-4(19),7-15-triene (**6**).

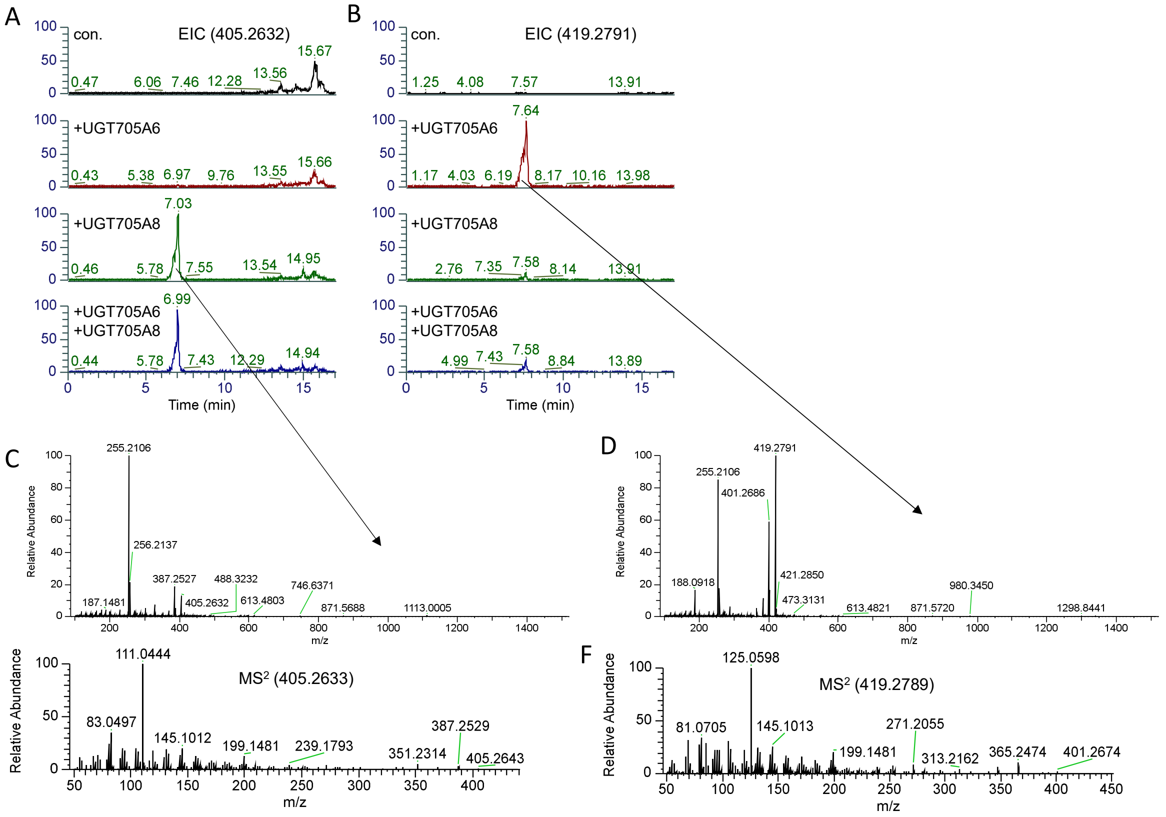


**Supplementary Figure 6. Transient expression of UGT705A6 and UGT705A8 in *Nicotiana benthamiana*.** **A, B**, LC-MS extracted ion chromatograms (EIC) of extracts from *N. benthamiana* leaves infiltrated with agrobacteria carrying expression vectors for UGT705A6 and UGT705A8, combined with other biosynthetic genes from BGC1(2D). **C**, **E**, MS and MS$^2$ spectra of aspisoside B (**8**). **D**, **F**, MS and MS$^2$ spectra of aspisoside A (**7**). Con., tHMGR+GGPPS+TaCPS_D2+TaKSL_D1+CYP99A7_2D+CYP99A8_2D+CYP99A10_2D.

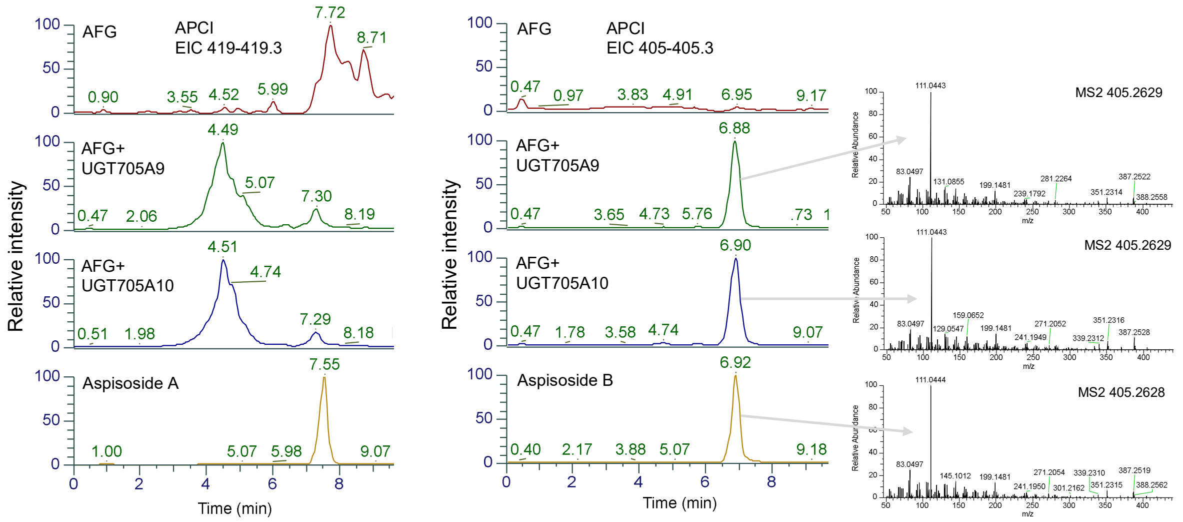


**Supplementary Figure 7. UGT705A9 and UGT705A10 exhibit arabinosyltransferase activity.** Transient expression of UGT705A9 or UGT705A10 in *N. benthamiana*, together with biosynthetic genes for aspisosol (Aspisosol Forming Genes; AFG), results in formation of aspisoside B, but not aspisoside A.

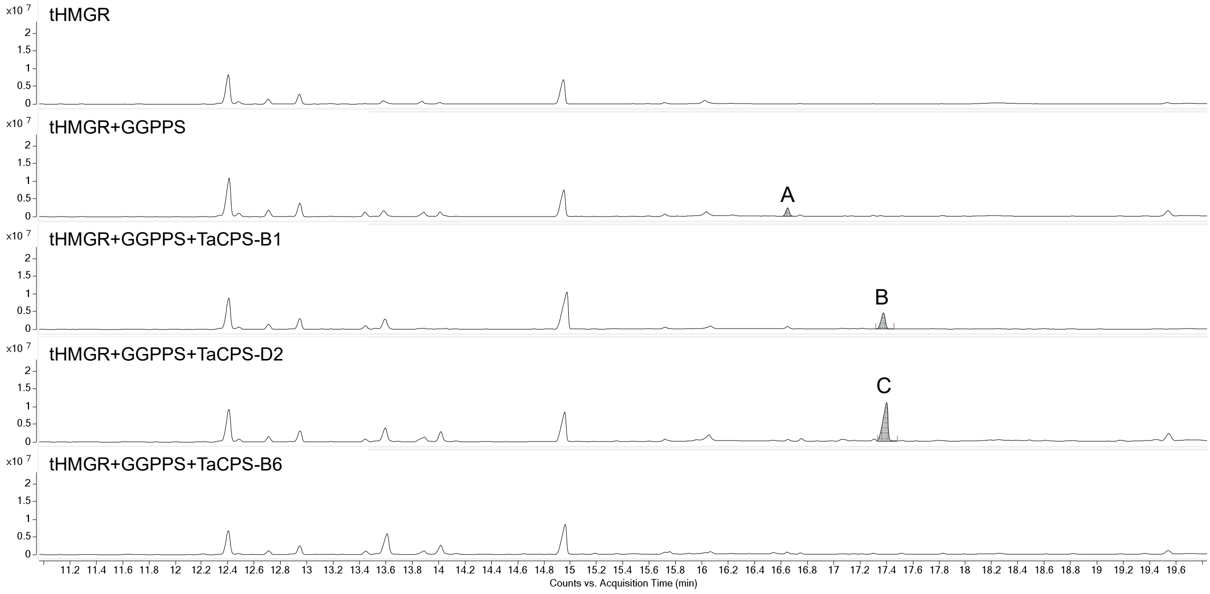


**Supplementary Figure 8. GC-MS chromatograms of leaf extracts following TaCPS transient expression in *N. benthamiana*.** A- geranylgeraniol; B- *ent*-copalol; C- (+)-copalol.

TGATATTTAG**GGTTGGTGCTCACTAGCAAATCACCATCATATCCGGAGGTAGAAGTGGGT**
**GAATTGGATGTTGATTACAATAGACGACACATG**GTGAGCATACATTACATATGAATTAAC
TCATTGAGCAGATTAATTAGCTACTACACACAAGCTTACAAACATATACTACTATACTGA
ACACTAATCAAAACAAAATATCAATATTCCTAAAG**GATGAGCCCAGCGAGATGCCACAGA**
**TGATTGATGCCATAAGAACGATGCTGGAGTTATTGGGTGATGGTGAAACATCGACGAATA**
**TTTCTGCCTACGACACGGCATTGGTCGCCCTTGTCAAGAACCTTGAGGGAGGTGATGGAC**
**CGCAATTCCCCTCATGCATTGACTGGATTGTTCAAAATCAACTACCGGATGGTTCATGGG**
**GTGACCCCGCTTTCTTTCTGGTCCAAGATCGGATGATAAGCACCCTGGCATGTGTGGTGG**
**CAGTGAAGTCTTGGAAAATTGACAACGCCAACTTGTGTGATAGAG**GTATATCAATAGCTG
GATTCTTTGTTTTGACTAAGAATTGACTTGTTTATTCCCACAAAAAGATCATTTGTCTAA
TGTGTAGAGGAACATTTTTGTTGTAG**GTGTGTCATTTATCCAAGAAAATATGAGCAGGTT**
**GGTAGAAGAGGAACAGGACTGGATGCCATCCGGCTTCGAGATTAACTTCCCAGCACTCCT**
**AGACAGGGCCAAGGACCTGTATTTGGACATCCCTTATAATCACCCTGTTTTGGAAGAGAT**
**ATATGCCAAGAGAAATCTAAAGCTCTCGAA**GTATATGCATACCTAAATCTAACTTGGATT
TGGACATTGAGTTGAATAGTTTATGTATTATTAGGCAATAATTAATCTGAGTCTCGAACT
ATAG**GATCCCTCTGGAACTACTACACGCCACACCAACCACCCTACTTTTCAGCTTGGAGG**
**GAATGGCAAACTTGCAGTTGGACTGGGAAAAGCTCCTCAAGTTGCGGTGTCCGGACGGCT**
**CCTTCCACTCATCGCCTGCTGCCACAGCTGCTGCCCTCTGTCACACCGGTGACAAGGAAT**
**GCCTTGCGTTTCTGGAGAGGCTCGTCAAGAAGTTCGACGGGGGAG**GTAATTCAAAATCAC
TCTACTTGTTGTACAACCGGCACAAATTCACATTTTTCAAGGGTAGGAGTCCCTTGCAAT
GCCGAATATGTATACTAAGAATTTCCCTTGTTTTCAG**TGCCATGTACCCACGCGATGGAT**
**ATTTTTGAGCACTTATGGGTGGTCGATCGGCTGATGCGTCTGGGGATATCAAGGCACTTC**
**ACATGTGAAATTGAGCAGTGCATGGATTATATTTACAG**GTAGCATAAACTATTTCCGTTG
TATTGCAAATACATGATATTTGGAAATAATTCCACTCAAGAGTTCCTAACTTAACTGCAG
TCAGGCATATACACGACTATTTTATTTGGTAAAGCCATCAAAGTATCAGCAGATTTATCA
TAAAATACATGCTCATATATAAATTTTGTTTAGCAACATATTGCGTATCAAAATAGCAGT
TAGAGGCATGCTGGAGACTTCCAAATCATAAATCAGGTTTTACTTTTAACCAAGGGAGTA
CATGTTTGGAAACTCATTGCCTGAACATAAGTTAGACTTTTACATGGTCTCCAACATTCA
TGATCGCTATTACTCATACCTATATGTTAGCATAAAACTAACCAAACTCATACTTTATAT
GAATAAATGTTAATAATTTCAGTGATACTACCTTCACCCAATACATCTAAATACTTGTTA
TATATATATATATATTTTGAAACGGAGTAATAATAGTGTTTTTCTTCAAG**ACCGATAGAG**
**TACTATTTTGAGCGGAGCTGTGTGTATATATACACTTTGAACAGGCGCTGGACTCAAAAG**
**GGACTGGCTCATAACGCGCACTGCCCAACCACGGATATTGATGACACGGCCATGGGTTTC**
**CGTCTCCTCCGTCAGCATGGCTATGACGTCACTCCAT**GTATGTAACTAATATTTTGTCAG
TCAAATTTATGAATAAAACTAGATTCAAAATTCGAGAAGTGCTTCTTTGTTTGTTTTGAG
GTAGTAGCGCCTTGCCATGCATAACTACTCCGTCATGGTAGATCAAGTTTGCCTAATGTG
CATCCACAATGCAG**CCGTGTTTAAGCACTTTGAGAAGGACGGCAAGTTTGTATGCTTCCC**
**CATGGAGACGAACCACTCATCTGTCACCCCAATGCACAACACTTATCGTGCATCTCAATT**
**TATGTTCGCCGGTGACGACAACGTCCTAGCACGGGCTGGGCACTATTGCCATTCATTCCT**
**CCAAGAGAGACAAGCCTCCAACAAACTGTACGACAAGTGGATCATCGCCAAGGACCTGCC**
**GGGTGAG**GTATGTCCCAACACATTCAAGTTAATTATATATGAGGAATTGCAGTCCAGCTA

**Supplementary Figure 9. Partial gene sequence of TaCPS-B6.** Automatic gene annotation according to the gene entry for TraesCS2B02G040400 on Ensemble Plants (https://plants.ensembl.org/). Predicted exons are highlighted. Green highlight- Start codon. In red font- sequence annotated as intronic but found in the cDNA-amplified coding sequence.

```
OsCPS1          KCFAYIDRIIKKFDGGVPNVYPVDLFEHIWVVDRLERLGISRYFQREIEQNMDYVN----     400
OsCPS2          ELLEYLETAINNFDGGAPCTYPVDNFDRLWSVDRLRRLGISRYFTSEIEEYLEYAY----     356
TaCPS-B1        KCLKYIDEIVKKFDGGAPCVYPVDLYERLWAVDRLTRLGISRHFTSEIKECLDFTY----     350
OsCPS4          KCFEYLDGIVKKFNGGVPCIYPLDVYERLWAVDRLTRLGISRHFTSEIEDCLDYIF----     347
TaCPS-B6        ECLAFLERLVKKFDGGVPCTHAMDIFEHLWVVDRLMRLGISRHFTCEIEQCMDYIYRPIE     283
HvCPS2          ECQAFLDRLIKKFDGGVPCSHSMDTFEQVWVVDRLMHLGISRHFTSEIDQFLEFIY----     342
TaCPS-D2        ECHAFLDRLIQKFEGGVPCSHSMDTFEQLWVVDRLMRLGISRHFTSEIQQCLEFIY----     379
                :   :::  :::*:**.*  : :* ::::* **** :*****:*  **.: :::

OsCPS1          --------------RHWT---EDGICWARNSNVKEVDDTAMAFRLLRLHGYNVSPSVFKN     443
OsCPS2          --------------RHL---SPDGMSYGGLCPVKDIDDTAMAFRLLRLHGYNVSSSVFNH     399
TaCPS-B1        --------------RHWTQVEDDGLSHAGSCSAADIDDTAMGFRLLRLNGYHVNPCALKK     396
OsCPS4          --------------RNWT---PDGLAHTKNCPVKDIDDTAMGFRLLRLYGYQVDPCVLKK     390
TaCPS-B6        YYFERSCVYIYTLNRRWT---QKGLAHNAHCPTTDIDDTAMGFRLLRQHGYDVTPSVFKH     340
HvCPS2          --------------RRWT---NKGLAHNVHCPIADIDDTAMGFRLLRQHGYEVNPSVFKQ     385
TaCPS-D2        --------------RRWT---QKGLAHNMHCPIPDIDDTAMGFRLLRQHGYDVTPSVFKH     422
                              *.      .*:.    .   ::*****.*****  **.*  ..:::
```

**Supplementary Figure 10. Multiple sequence alignment of CPS protein sequences from wheat, rice, and barley.** Protein sequence alignment shows an 18 amino acid-long sequence found in TaCPS-B6 that is absent from related CPS proteins. The alignment was carried out with Clustal Omega (https://www.ebi.ac.uk/jdispatcher/msa/clustalo) using default parameters.

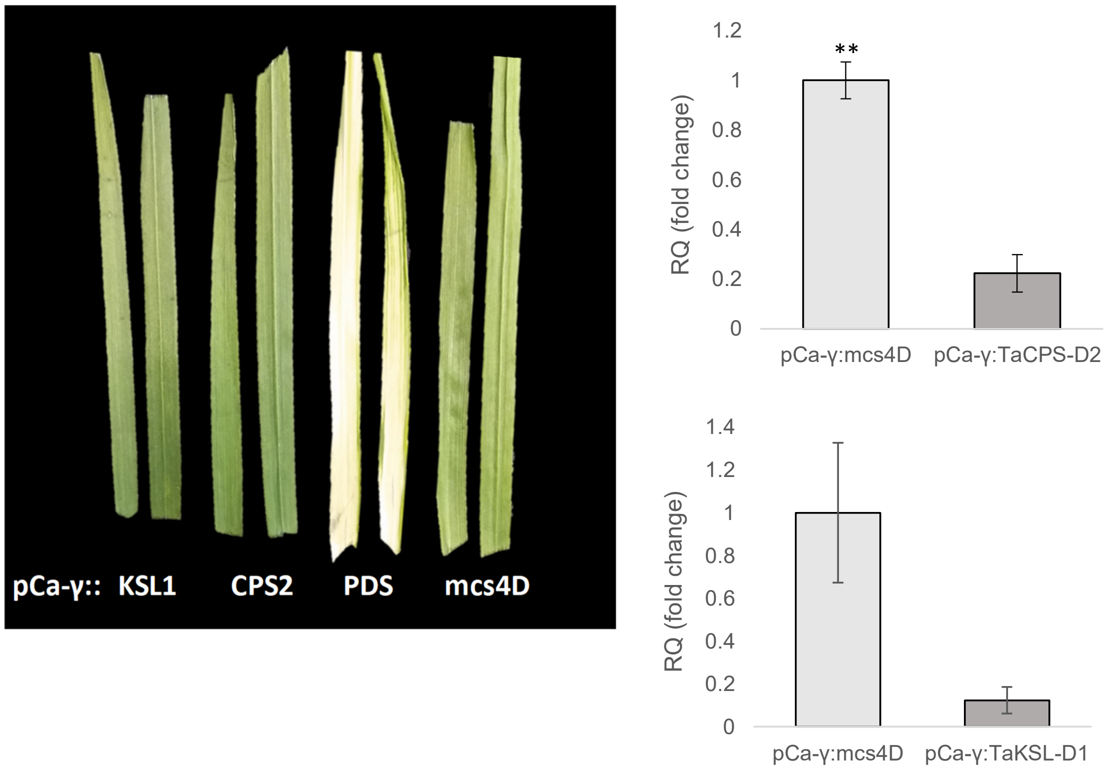


**Supplementary Figure 11. Virus Induced Gene Silencing of *TaCPS-D2* and *TaKSL-D1* in wheat. A,** bleaching phenotype was observed in wheat plants in which phytoene desaturase was silenced by expression of pCa-γ::PDS. **B**, qRT-PCR analysis shows respective reduction of transcript abundance of *TaCPS-D2* and *TaKSL-D1* in plants in which pCa-γ::TaCPS-D2 or pCa-γ::TaKSL-D1 were expressed, compared to control, pCa-γ::mcs4D. Asterisks denote t-test statistical significance of differential expression. **, p-val<0.01.

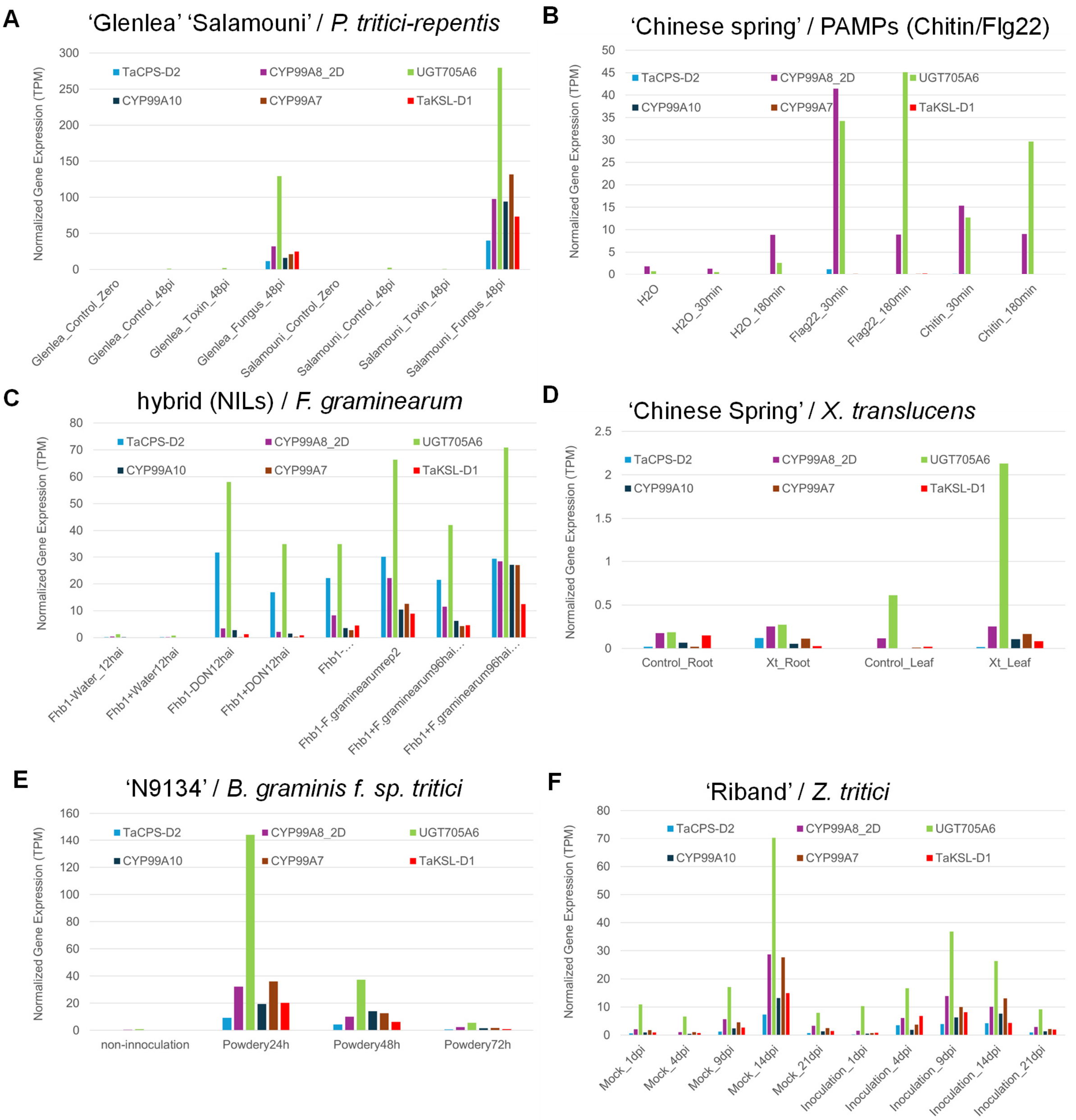


**Supplementary Figure 12. Gene expression data of BGC1(2D) genes in six biotic stress experiments**. Normalized values (transcripts per million) are shown, derived from RNA-seq data obtained from Wheatomics 1.0[8] (http://wheatomics.sdau.edu.cn/). **A**, infection of wheat varieties 'Glenlea' and 'Salamouni' with fungal pathogen *Pyrenophora tritici-repentis* or infiltration with the toxin Ptr ToxA (https://www.ncbi.nlm.nih.gov/bioproject/PRJNA327829). **B**, elicitation of cv. 'Chinese Spring' plants with PAMPs chitin or Flg22[7]. **C**, infection with *Fusarium graminearum* of wheat near isogenic lines that are resistant (Fhb1+) or susceptible (Fhb1-) to Fusarium head blight[9]. **D**, infection of cv. 'Chinese Spring' plants with bacterial pathogen *Xanthomonas translucens*[10]. **E**, infection of wheat line N9134 with powdery mildew (*Blumeria graminis* f. sp. *tritici*; *Bgt*)[11]. **F**, infection of cv. 'Riband' plants with *Zymoseptoria tritici*[12].

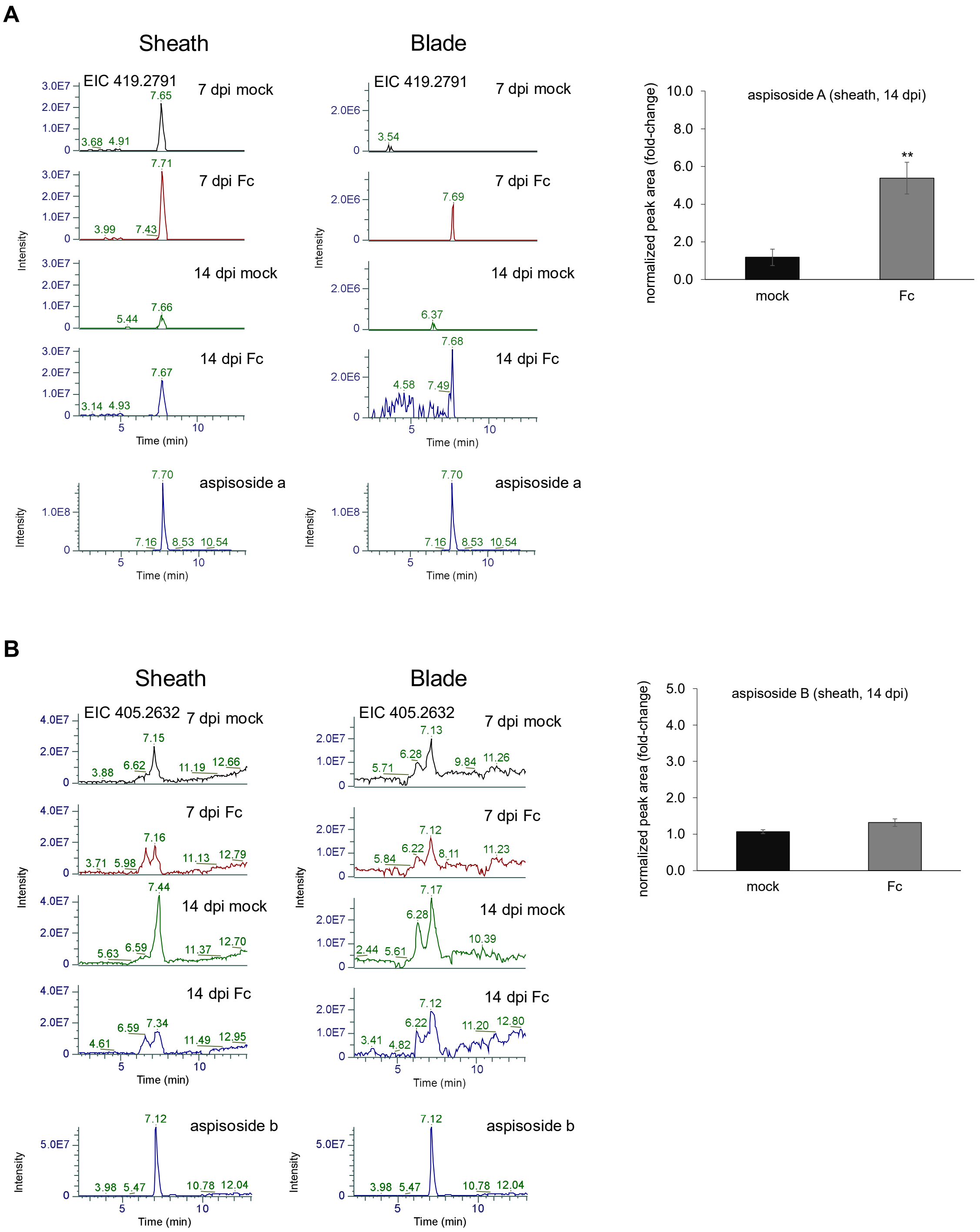


**Supplementary Figure 13. Extracted ion chromatograms of aspisoside A and aspisoside B in wheat.** Aspisoside A (**A**) and aspisoside B (**B**) were detected in extracts of sheath and blade tissues from wheat plants infected with *Fusarium culmorum*, 7 and 14 days post infection. Relative quantification (fold-change) values indicate means of three biological replicates ± SEM. Asterisks denote the statistical significance of a two-tailed *t*-test. $^{**}P < 0.01$.

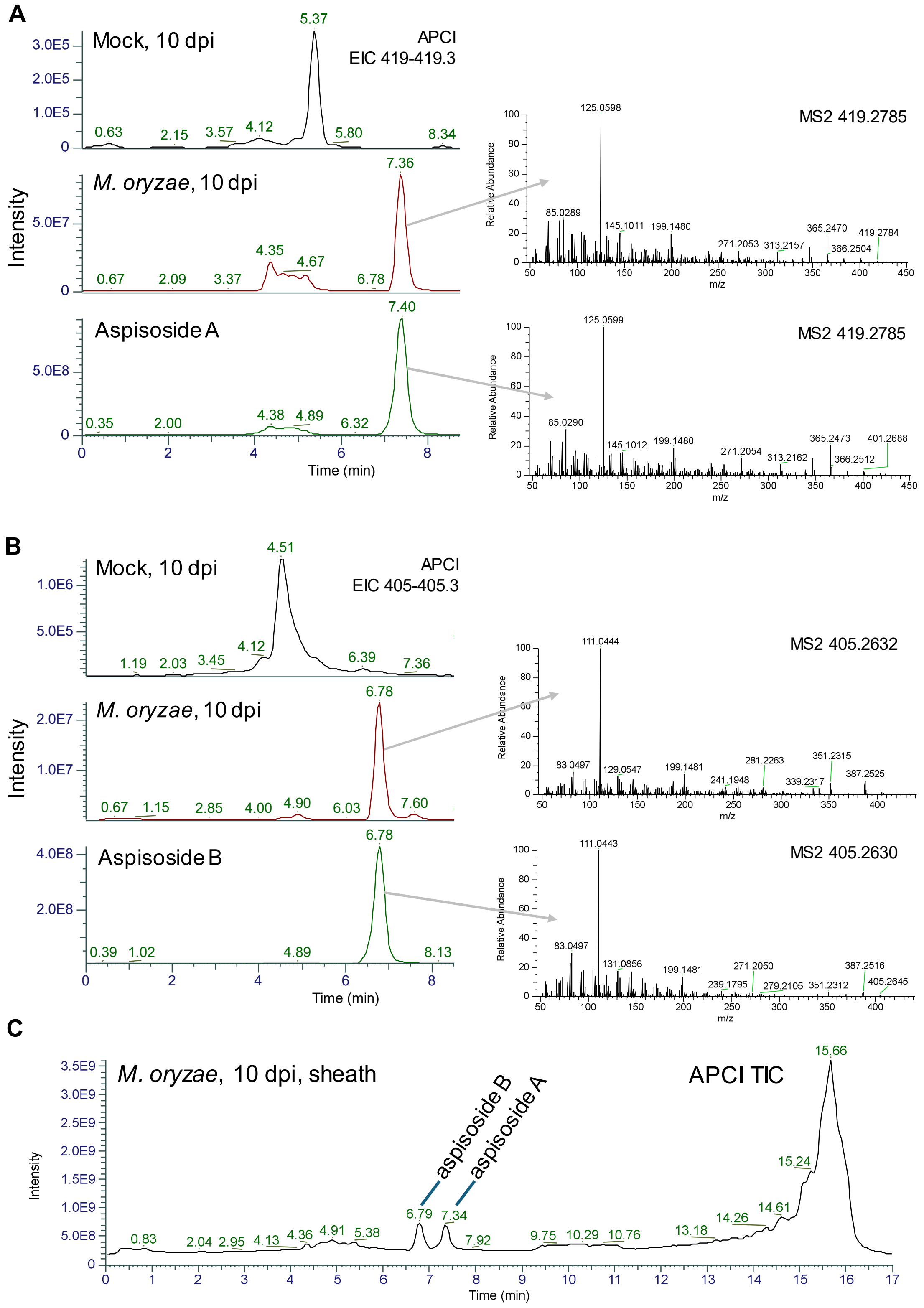


**Supplementary Figure 14. Extracted ion chromatograms and MS$^2$ spectra of aspisoside A and aspisoside B in wheat.** Aspisoside A (**A**) and aspisoside B (**B**) were detected in extracts of sheath tissue from wheat plants infected with *Magnaporthe oryzae*, 10 days post infection (dpi), exhibiting the same retention time and MS$^2$ as the compounds purified from infiltrated *N. benthamiana* leaves. **C**, APCI total ion chromatogram of sheath extract from *M. oryzae*-infected plants, 10 dpi.

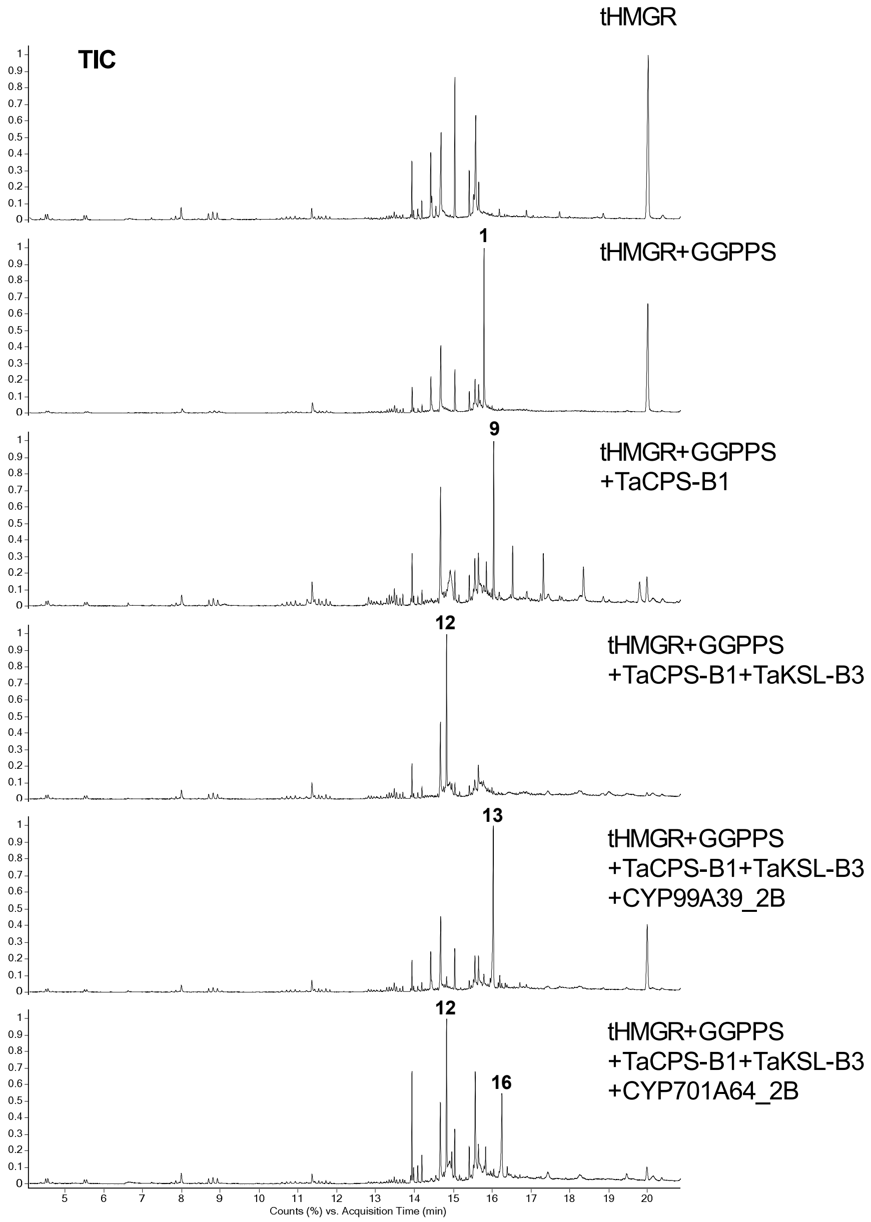


**Supplementary Figure 15. GC-MS total ion chromatograms (TIC) of combinatorial transient expression experiments of scutenol B biosynthetic genes in *N. benthamiana*.** GGPP (**1**), *ent*-copalol (**9**), (Z)-biformene (**12**), (Z)-biformene-3α-ol (scutenol B) (**13**), (Z)-biformene-19-ol (**16**).

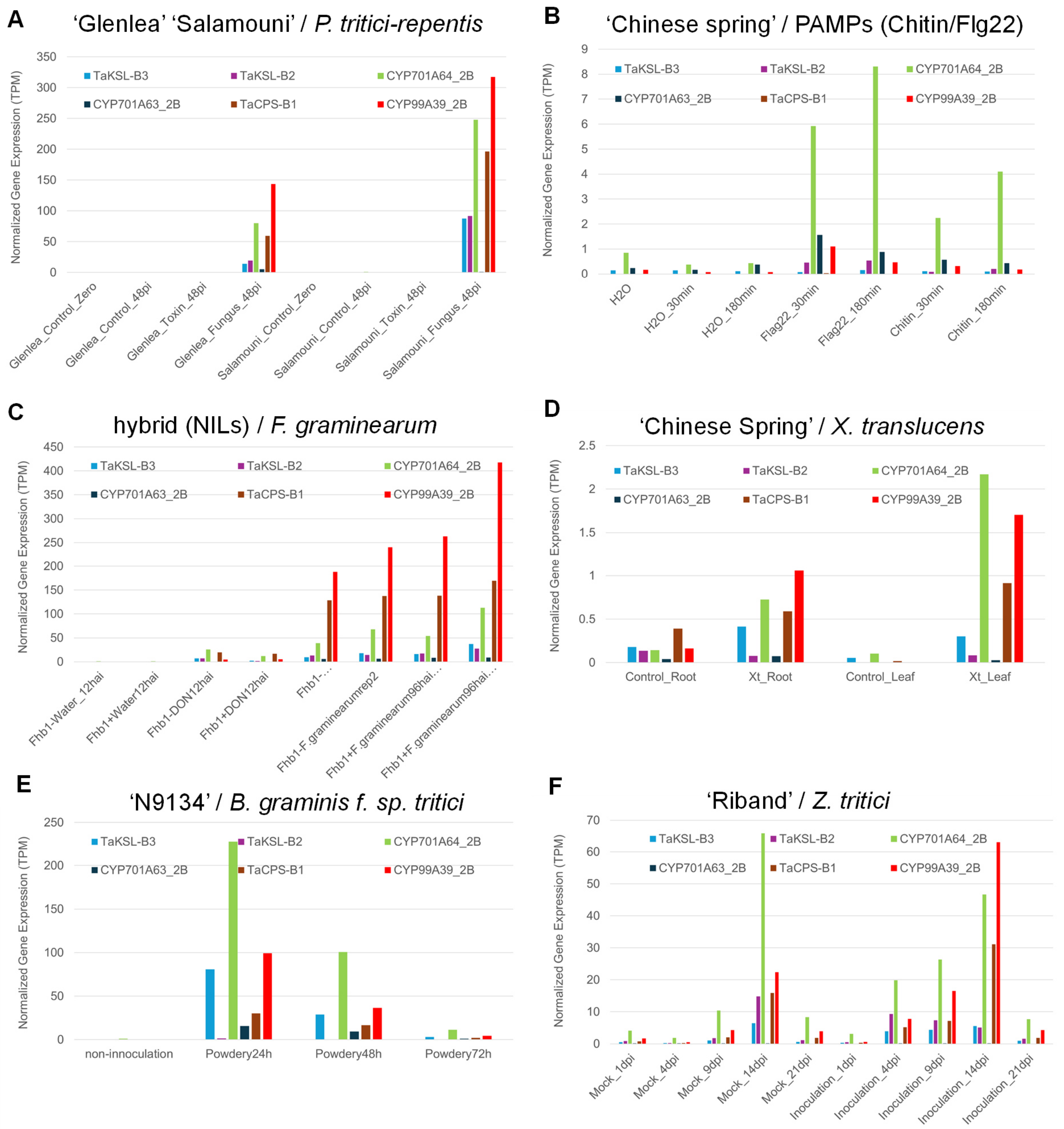


**Supplementary Figure 16. Gene expression data of BGC2(2B) genes in six biotic stress experiments**. Normalized values (transcripts per million) are shown, derived from RNA-seq data obtained from Wheatomics 1.0[8] (http://wheatomics.sdau.edu.cn/). **A**, infection of wheat varieties 'Glenlea' and 'Salamouni' with fungal pathogen *Pyrenophora tritici-repentis* or infiltration with the toxin Ptr ToxA (https://www.ncbi.nlm.nih.gov/bioproject/PRJNA327829). **B**, elicitation of cv. 'Chinese Spring' plants with PAMPs chitin or Flg22[7]. **C**, infection with *Fusarium graminearum* of wheat near isogenic lines that are resistant (Fhb1+) or susceptible (Fhb1-) to Fusarium head blight[9]. **D**, infection of cv. 'Chinese Spring' plants with bacterial pathogen *Xanthomonas translucens*[10]. **E**, infection of wheat line N9134 with powdery mildew (*Blumeria graminis* f. sp. *tritici*; *Bgt*)[11]. **F**, infection of cv. 'Riband' plants with *Zymoseptoria tritici*[12].

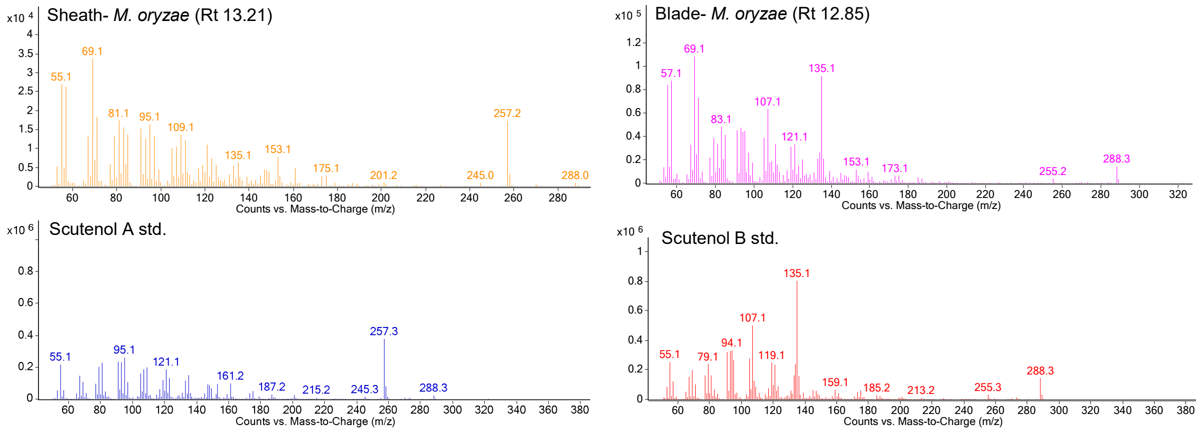


**Supplementary Figure 17. Comparison of GCMS mass spectra of scutenol A and scutenol B detected in wheat to compounds purified from *N. benthamiana*.**

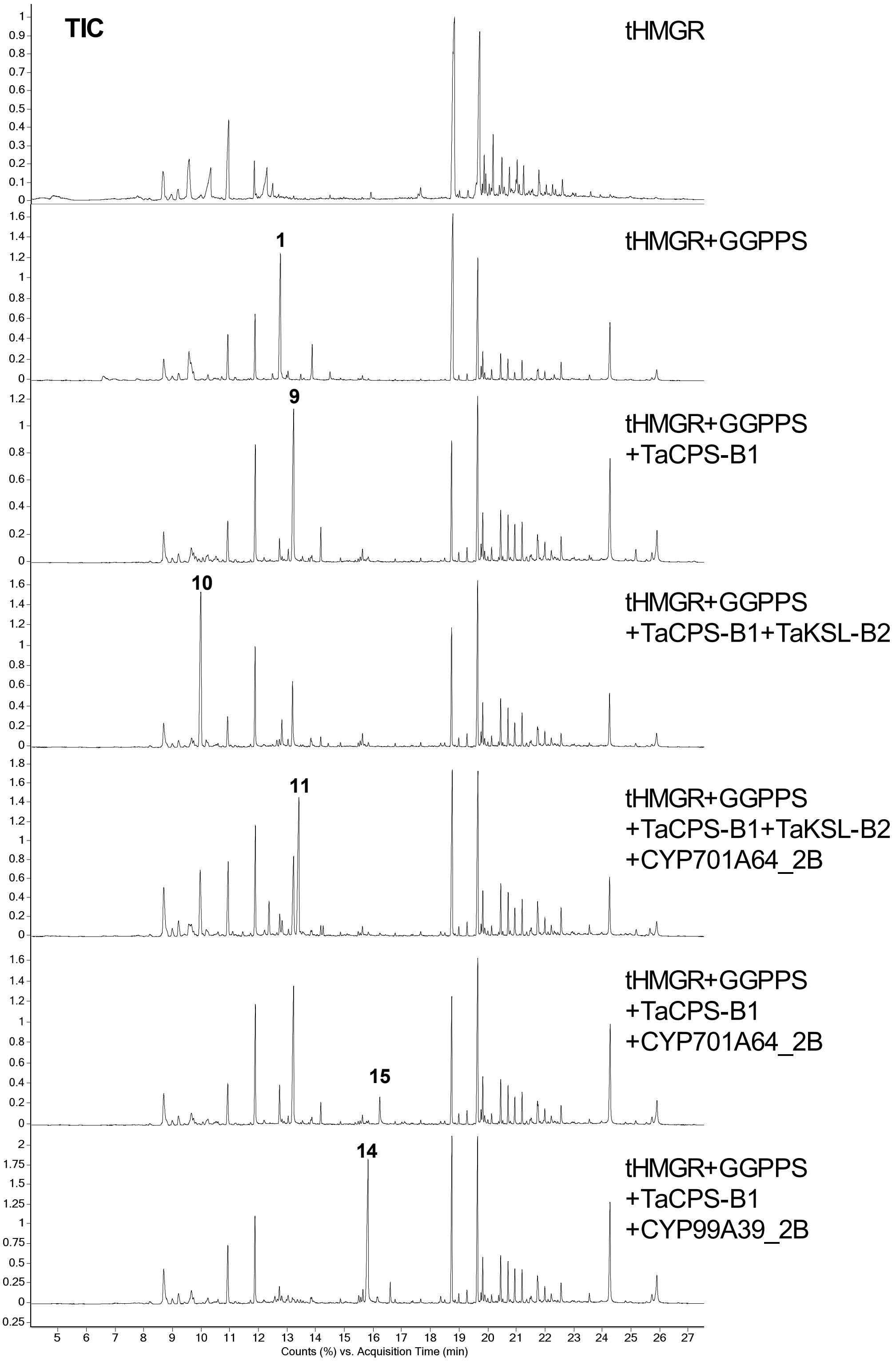


**Supplementary Figure 18. GC-MS total ion chromatograms (TIC) of combinatorial transient expression experiments of scutenol A biosynthetic genes in *N. benthamiana*.** GGPP (**1**), *ent*-copalol (**9**), *ent*-pimara-8(14),15-diene (**10**), *ent*-pimara-8(14),15-diene-19-ol (scutenol A) (**11**), *ent*-15,19-dihydroxy-labda-8(17),13(E)-diene (**14**). *ent*-3α,15-dihydroxy-labda-8(17),13(E)-diene (**15**).

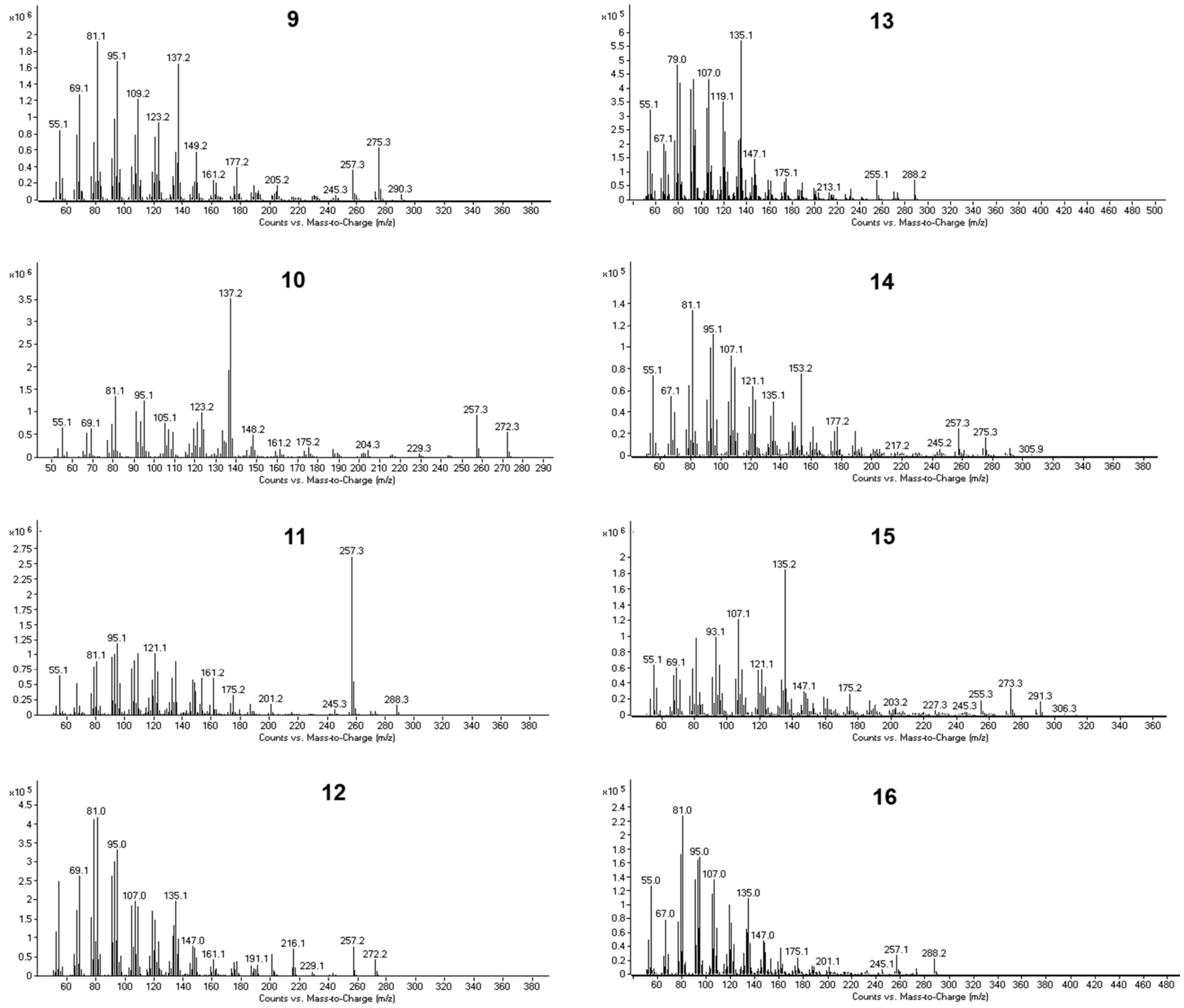


**Supplementary Figure 19. GC-MS mass spectra of BGC2(2B)-encoded pathway products and intermediates.** GGPP (**1**), *ent*-copalol (**9**), *ent*-pimara-8(14),15-diene (**10**), *ent*-pimara-8(14),15-diene-19-ol (scutenol A) (**11**), (Z)-biformene (**12**), (Z)-biformene-3α-ol (scutenol B) (**13**), *ent*-15,19-dihydroxy-labda-8(17),13(E)-diene (**14**), *ent*-3α,15-dihydroxy-labda-8(17),13(E)-diene (**15**), (Z)-biformene-19-ol (**16**).

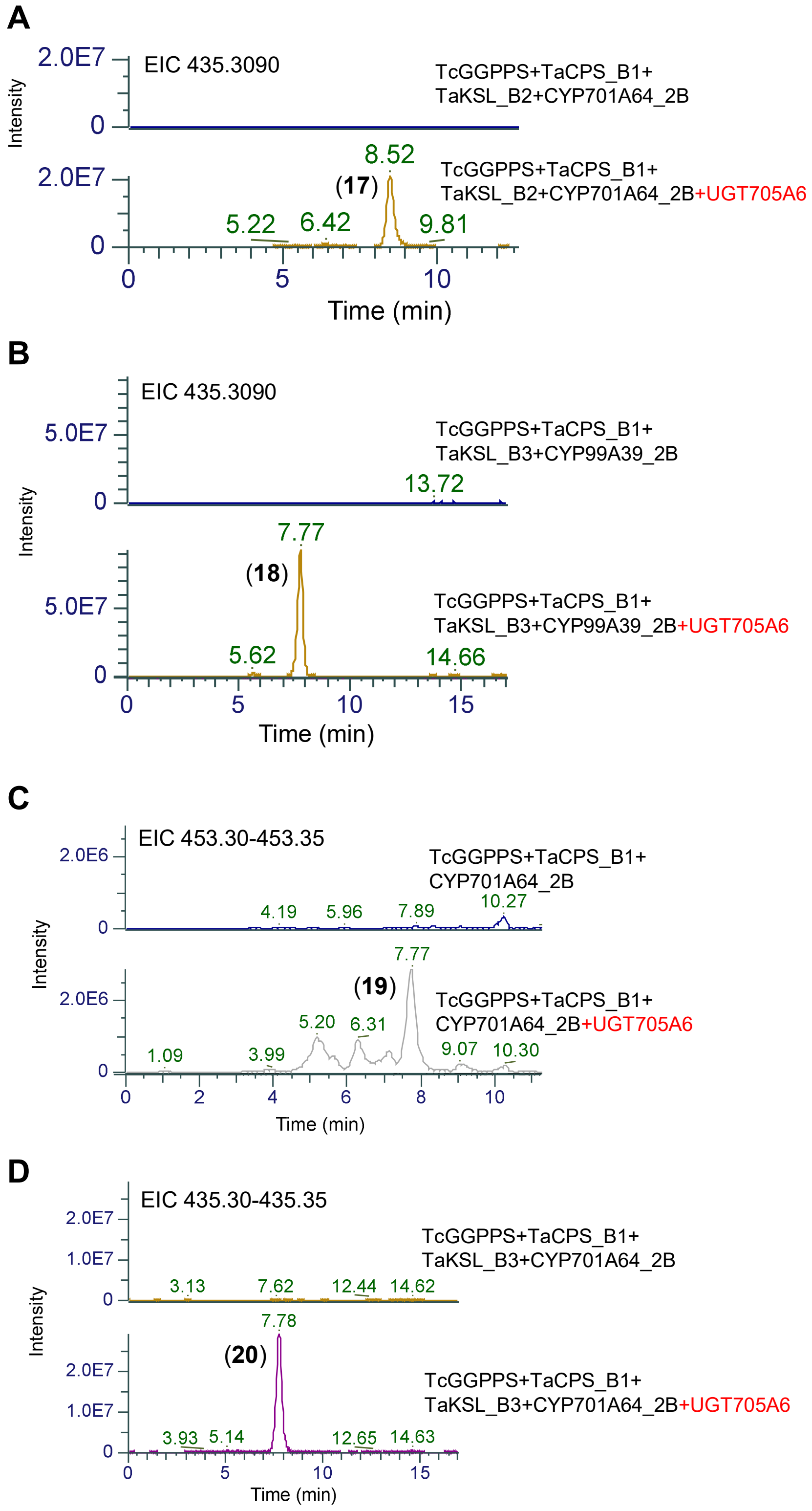


**Supplementary Figure 20. UGT705A6 glycosylates BGC2(2B)-encoded products.** Transient co-expression of UGT705A6 in *N. benthamiana* with BGC2(2B) genes results in formation of molecules with masses corresponding to an addition of a deoxyhexose (+146 Da). *ent*-pimara-8(14),15-diene-19-deoxyhexoside (**17**), (Z)-biformene-3-deoxyhexoside (**18**), *ent*-15-hydroxy-labda-8(17),13(E)-diene-19-deoxyhexoside (**19**), (Z)-biformene-19-deoxyhexoside (**20**).

**Table S1- Co-expression analysis of BGC1(2A) and BGC1(2D) genes**

**BGC1(2A) (TaKSL-A1 bait)**

| Assigned gene name | IWGSC refseq ID | Annotation | PCC |
|---|---|---|---|
| TaKSL-A7 | TraesCS2A02G027600 | Kaurene synthase | 0.16 |
| CYP99A9_2A | TraesCS2A02G027700 | Cytochrome P450 | 0.36 |
| TaCPS-A11 | TraesCSU02G252000 | Copalyl diphosphate synthase | 0.07 |
| **TaKSL-A1** | **TraesCSU02G008700** | **Kaurene synthase** | **1** |
| CYP99A7_2A | TraesCSU02G008800 | Cytochrome P450 | 0.91 |
| CYP99A10_2A | TraesCSU02G008900 | Cytochrome P450 | 0.90 |
| UGT705A7 | TraesCSU02G009000 | Glycosyltransferase | 0.82 |
| CYP99A8_2A | TraesCSU02G009100 | Cytochrome P450 | 0.85 |
| TaCPS-A2 | TraesCS2A02G027800 | Copalyl diphosphate synthase | 0.83 |
| - | TraesCS2A02G027900 | RING/FYVE/PHD zinc finger superfamily protein | -0.08 |
| UGT705A8 | TraesCS2A02G028000 | Glycosyltransferase | 0.78 |
| TaKSL-A4 | TraesCS2A02G028100 | Kaurene synthase | -0.05 |
| TaCPS-A8 | TraesCS2A02G028200 | Copalyl diphosphate synthase | 0 |

**BGC1(2D) (TaKSL-D1 bait)**

| Assigned gene name | IWGSC refseq ID | Annotation | PCC |
|---|---|---|---|
| CYP71F22_2D | TraesCS2D02G029300 | Cytochrome P450 | 0.30 |
| TaKSL-D7 | TraesCS2D02G029400 | Kaurene synthase | -0.01 |
| CYP99A9_2D | TraesCS2D02G029500 | Cytochrome P450 | 0.40 |
| TaCPS-D2 | TraesCS2D02G029600 | Copalyl diphosphate synthase | 0.73 |
| CYP99A8_2D | TraesCS2D02G029700 | Cytochrome P450 | 0.79 |
| UGT705A6 | TraesCS2D02G029800 | Glycosyltransferase | 0.77 |
| CYP99A10_2D | TraesCS2D02G029900 | Cytochrome P450 | 0.82 |
| CYP99A7_2D | TraesCS2D02G030000 | Cytochrome P450 | 0.87 |
| **TaKSL-D1** | **TraesCS2D02G030100** | **Kaurene synthase** | **1** |
| TaKSL-D4 | TraesCS2D02G030200 | Kaurene synthase | -0.11 |
| TaCPS-D8 | TraesCS2D02G030300 | Copalyl diphosphate synthase | 0 |

PCC: Pearson Correlation Coefficient

### Table S2- Oligos used in this study

| | |
|---|---|
| **Primers for CDS cloning into pDONR207 (5'-3')** | |
| TcGGPPS-F | GGGGACAAGTTTGTACAAAAAAGCAGGCTATGGCTTCCTATCAAGAATGC |
| TcGGPPS-R | GGGGACCACTTTGTACAAGAAAGCTGGGTTCAGTTTTGCCTGAATGCAATG |
| TaCPS-D2-F | GGGGACAAGTTTGTACAAAAAAGCAGGCTATGATTAGCAAATCCCCATCCTATC |
| TaCPS-D2-R | GGGGACCACTTTGTACAAGAAAGCTGGGTCTAAATGACATCCTGAAATATG |
| TaKSL-D1-F | GGGGACAAGTTTGTACAAAAAAGCAGGCTATGGCAATGTTGGCAGAAGAAACC |
| TaKSL-D1-R | GGGGACCACTTTGTACAAGAAAGCTGGGTTTATTGTTCTGACTGAGCATCCAAAG |
| CYP99A8_2D-F | GGGGACAAGTTTGTACAAAAAAGCAGGCTATGGAGTTAACAGGAGCCACATTAT |
| CYP99A8_2D-R | GGGGACCACTTTGTACAAGAAAGCTGGGTTCAACTGCATGCCTTATACGGTG |
| CYP99A10_2D-F | GGGGACAAGTTTGTACAAAAAAGCAGGCTATGGAGCTAAGCGCAGCCACCCTACT |
| CYP99A10_2D-R | GGGGACCACTTTGTACAAGAAAGCTGGGTTTAAAAACTGCATGCCTTGCACGGTG |
| CYP99A7_2D-F | GGGGACAAGTTTGTACAAAAAAGCAGGCTATGGAGATGGAGCTAAGCCTAGGCG |
| CYP99A7_2D-R | GGGGACCACTTTGTACAAGAAAGCTGGGTTCAGCTTTGCAGTGGAACCTCCTTATAC |
| UGT705A6-F | GGGGACAAGTTTGTACAAAAAAGCAGGCTATGGCTTCAGCCACTAGCAGCGGCA |
| UGT705A6-R | GGGGACCACTTTGTACAAGAAAGCTGGGTCTATCCTCTTGCTGCGATGAGATCATTG |
| UGT705A8-F | GGGGACAAGTTTGTACAAAAAAGCAGGCTATGGCTTCTACCACTAGCAGCGGC |
| UGT705A8-R | GGGGACCACTTTGTACAAGAAAGCTGGGTTTAGCCTCTTGCTGCCATGAGATC |
| UGT705A9-F | GGGGACAAGTTTGTACAAAAAAGCAGGCT ATGGCTTCTTCTACCACTAGCAA |
| UGT705A9-R | GGGGACCACTTTGTACAAGAAAGCTGGGTCTAGCTCCGCCCCACCTTTGC |
| UGT705A10-F | GGGGACAAGTTTGTACAAAAAAGCAGGCTATGGCTTCTTCTATCACTAGCAG |
| UGT705A10-R | GGGGACCACTTTGTACAAGAAAGCTGGGTCTAGTTCCGCCCCTCCTTTGCTGC |
| TaCPS-B1-F | GGGGACAAGTTTGTACAAAAAAGCAGGCTATGGGCGTGGTGAATGTGCAAACC |
| TaCPS-B1-R | GGGGACCACTTTGTACAAGAAAGCTGGGTCTAAATGACATCTTCGAATAAGACCTTG |
| TaKSL-B2-F | GGGGACAAGTTTGTACAAAAAAGCAGGCTATGGCCGAAGAAAATGCATGTCTCC |
| TaKSL-B2-R | GGGGACCACTTTGTACAAGAAAGCTGGGTTTAATTTACTGACTGCATGGGCAAAG |
| CYP701A64_2B-F | GGGGACAAGTTTGTACAAAAAAGCAGGCTATGGAGTCGCTGCTGGCGGCGCTCGC |
| CYP701A64_2B-R | GGGGACCACTTTGTACAAGAAAGCTGGGTTCACTTGCTCCCTCTCGGAGAGAGATGC |
| CYP99A39_2B-F | GGGGACAAGTTTGTACAAAAAAGCAGGCTATGGAGCTAAGCGCAGCCACCCTC |
| CYP99A39_2B-R | GGGGACCACTTTGTACAAGAAAGCTGGGTTCAAAGTTCCATGGGAACATTGTACG |
| **Primers for Goldenbraid cloning** | |
| TaCPS-B6-F | GCGCCGTCTCGCTCGAATGGATGAGCCCAGCGAGAT |
| TaCPS-B6-R | GCGCCGTCTCGCTCAAAGCCTAAATGACATCCTGGAATATGAC |
| **Primers for VIGS (5'-3')** | |
| TaCPS-D2-F | AAGGAAGTTTAAGTATATTTATAGCAGCAAACTTCTG |
| TaCPS-D2-R | AACCACCACCACCGTGAAGAACCCTCTTTCTTTCTTTTTTTGAC |
| TaKSL-D1-F | AAGGAAGTTTAACACTCAAAACATCACCTCCAAGTATC |
| TaKSL-D1-R | AACCACCACCACCGTCCACTCAAACTCCAACATACCAGTC |
| **Primers for qRT-PCR (5'-3')** | |
| TUBB-F | CAAGGAGGTGGACGAGCAGATG |
| TUBB-R | GACTTGACGTTGTTGGGGATCCA |
| TaCPS-D2-F | GATGGACCGCAATTTCCCTC |
| TaCPS-D2-R | CAAGACTTCACTGCCACGAC |
| TaKSL-D1-F | AGGATGAACTCTGCCACCTC |
| TaKSL-D1-R | TCATGAGTGGATAGGGTGGC |
| CYP99A8_2D-F | GGCCCTGGACAAGATGATCT |
| CYP99A8_2D-R | ATGGGGAAGTCAAGCTCTCC |
| CYP99A10_2D-F | GGTTCGAGTACTTGCCGTTC |
| CYP99A10_2D-R | AGTAGTAGAGAAGCCGTGCC |
| CYP99A7_2D-F | TTGAGATCGTCGAGGGTTCC |
| CYP99A7_2D-R | GCCATACTGTCCTCGAACCT |
| UGT705A6-F | TATGGACCTCGCCTCAAAGT |
| UGT705A6-R | TCCTCTTGCTGCGATGAGAT |
| UGT705A8-F | GGATGTGCTAGGAATCGGGG |
| UGT705A8-R | ATGGCCGCATGAGACTTTGA |
