## Supplementary Data 4 for "Diverse Diterpenoid Phytoalexins Shape Wheat Chemical Defenses"

**General Procedures**

Organic solvents used for extraction and flash chromatography were reagent grade and used directly without further distillation. HPLC mobile phases were prepared using HPLC grade solvents. LC-MS spectral data were recorded on Q-exactive orbitrap mass spectrometers using Kinetex-XB-$C_{18}$ (50 × 10 mm i.d.; 2.6 μm; USA), (JIC, UK). 1D and 2D NMR spectra were recorded on Bruker Avance 600 MHz spectrometer equipped with a BBFO Plus Smart probe and a triple resonance TCI cryoprobe, respectively (JIC, UK). The chemical shifts are relative to the residual signal solvent ($CDCl_3$ $\delta_H$ 7.26; $\delta_C$ 77.16). Preparative and Semi-preparative HPLC was performed on an Agilent 1290 Infinity (II) system using Luna C18 columns (250 x 21.2 and 250 × 10 mm i.d.; 5 μm).

**1- Extraction, purification and structural characterization of diterpene glycosides 7 and 8**

*TaCPS_D2, TaKASL_D1, TaCYP99A8_2D, TaCYP99A10_2D, TaCYP99A7_2D* were transiently co-expressed separately with *TaUGT705A6* and *TaUGT705A8* in **100** *Nicotiana benthamiana* plants to afford **136 g** of dry leaves, respectively. Subsequently, it was exhaustively extracted using ethyl acetate via speed extractor. The organic extract was collected and dried under reduced pressure to afford an intense green viscus material. Afterwards, it was introduced to normal phase flash chromatography using a gradient of dichloromethane/methanol [100/0 up 80/20] along 30 minutes, affording 12 subfractions F1-F12, where all have been monitored by LC-MS. Promising fraction F6 was further introduced to normal phase flash chromatography using a gradient of dichloromethane/methanol [100/0 up 90/10] along 30 minutes, affording twelve subfractions, where F11 was further purified using repetitive $C_{18}$ semi-preparative HPLC using a long gradient of water/acetonitrile [15/85 up 0/100], 4 mL/min, acidified with 0.1 % FA along 20 min to finally afford 3-*O*-$\alpha$-L-rhamnopyranosyl-18-Norisopimara-4(19),7,15-triene **(7)** (Rt: 7.64, **3.2** mg, pale yellow material) and 3-O-$\alpha$-L-arabinopyranosyl-18-Norisopimara-4(19),7,15-triene **(8)** (Rt: 7.02, **6.5** mg, white powder). All structures were resolved based on extensive HRMS and 1 & 2D-NMR analysis.

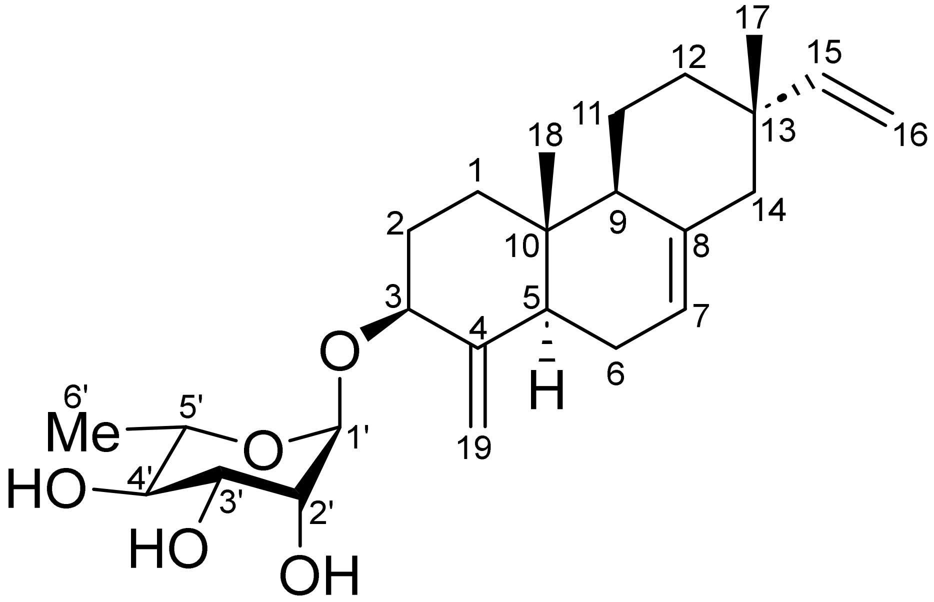

**3-*O*-α-L-rhamnopyranosyl-18-Norisopimara-4(19),7,15-triene (7)**

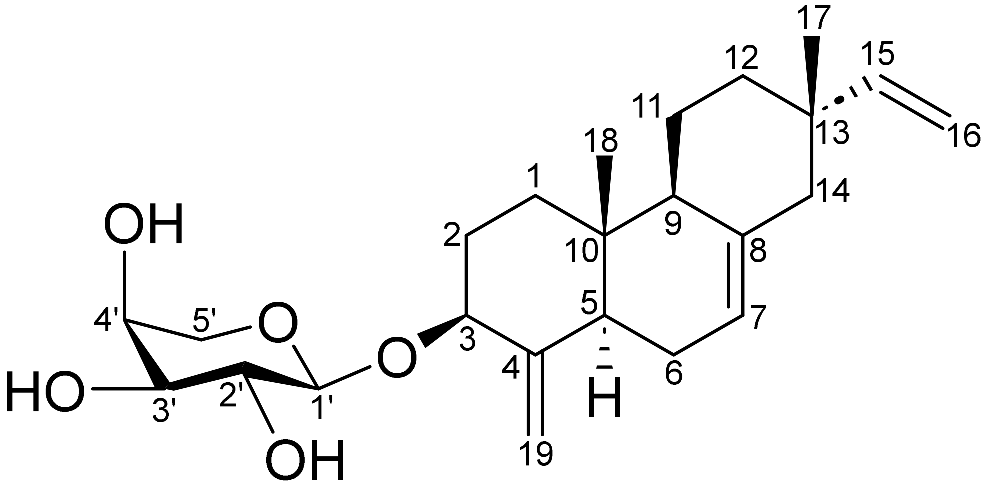

**3-*O*-α-L-arabinopyranosyl-18-Norisopimara-4(19),7,15-triene (8)**

**Table 1**: [1]H, [13]C NMR spectroscopic data for 3-*O*-*α*-L-rhamnopyranosyl-18-Norisopimara-4(19),7,15-triene **(7)** in $CDCl_3$, 600 and 150 MHz.

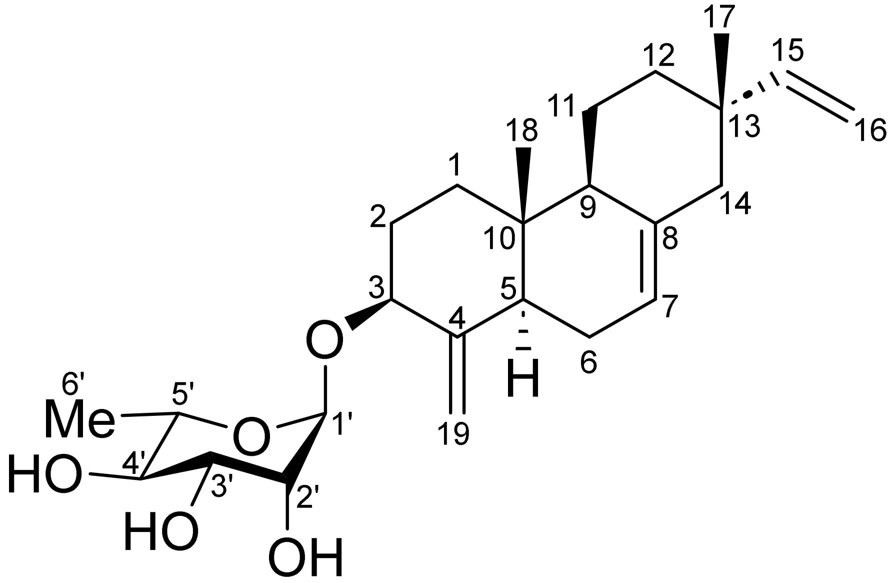

| **Position** | $\delta_C$**, Type** | $\delta_H$**, mult. (*J* in Hz)** |
|---|---|---|
| **1** | 37.4, $CH_2$ | 1.92/1.31, m |
| **2** | 31.1, $CH_2$ | 1.91/1.51, m |
| **3** | 79.1, CH | 4.00, m |
| **4** | 149.1, Cq | - |
| **5** | 45.6, CH | 1.90, m |
| **6** | 25.3, $CH_2$ | 2.07/1.86 |
| **7** | 120.4, CH | 5.37, m |
| **8** | 135.6, Cq | - |
| **9** | 49.7, CH | 1.76, m |
| **10** | 37.5, Cq | - |
| **11** | 21.5, $CH_2$ | 1.65/1.38, m |
| **12** | 36.5, $CH_2$ | 1.50/1.40, m |
| **13** | 37.1, Cq | - |
| **14** | 46.3, $CH_2$ | 2.00/1.94, m |
| **15** | 150.4, CH | 5.82, *dd* (17.5, 10.7) |
| **16** | 109.5, $CH_2$ | 4.94, *dd* (17.5, 1.3) |
| | | 4.88, *dd* (10.7, 1.3) |
| **17** | 21.6, $CH_3$ | 0.88, *s* |
| **18** | 14.1, $CH_3$ | 0.69, s |
| **19** | 104.5, $CH_2$ | 5.03, *d* (1.3)/4.70, *d* (1.1) |
| **Rham-1′** | 99.3, CH | 4.88, *d* (1.2) |
| **Rham-2′** | 71.2, CH | 4.07, d (1.6) |
| **Rham-3′** | 72.0, CH | 3.86, m |
| **Rham-4′** | 74.0, CH | 3.47, m |
| **Rham-5′** | 68.0, CH | 3.83, m |
| **Rham-6′** | 17.7, $CH_3$ | 1.32, *d* (6.3) |

**Scheme 1**: Key $^{1}H$-$^{1}H$ COSY (**bold blue**), $^{1}H$-$^{13}C$ HMBC (H→C, red arrows) observed 3-*O*-*α*-L-rhamnopyranosyl-18-Norisopimara-4(19),7,15-triene **(7)**

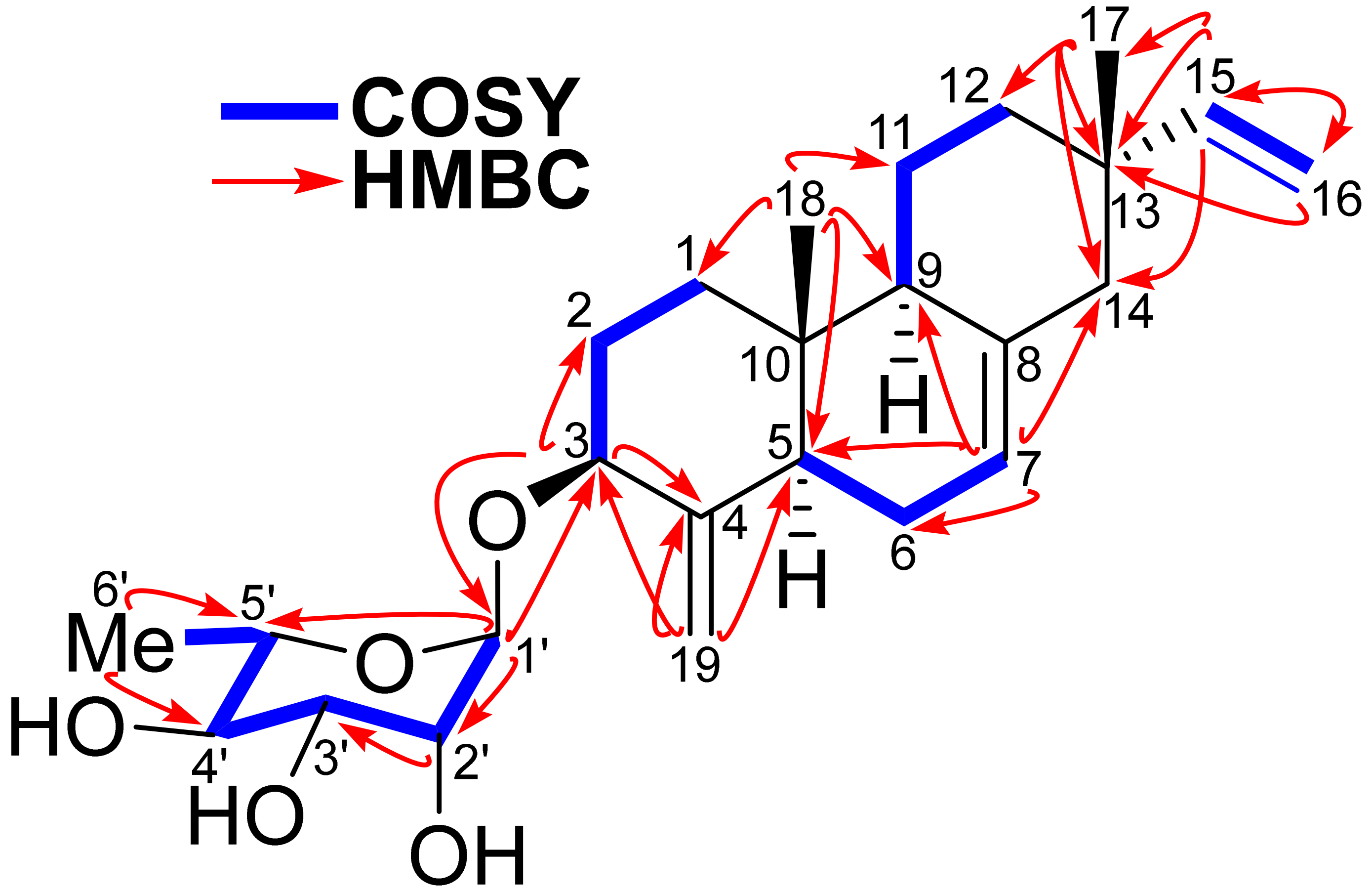

**Scheme 2**: Key $^{1}H$-$^{1}H$ ROESY (H↔H, green) observed for 3-*O*-*α*-L-rhamnopyranosyl-18-Norisopimara-4(19),7,15-triene **(7)**

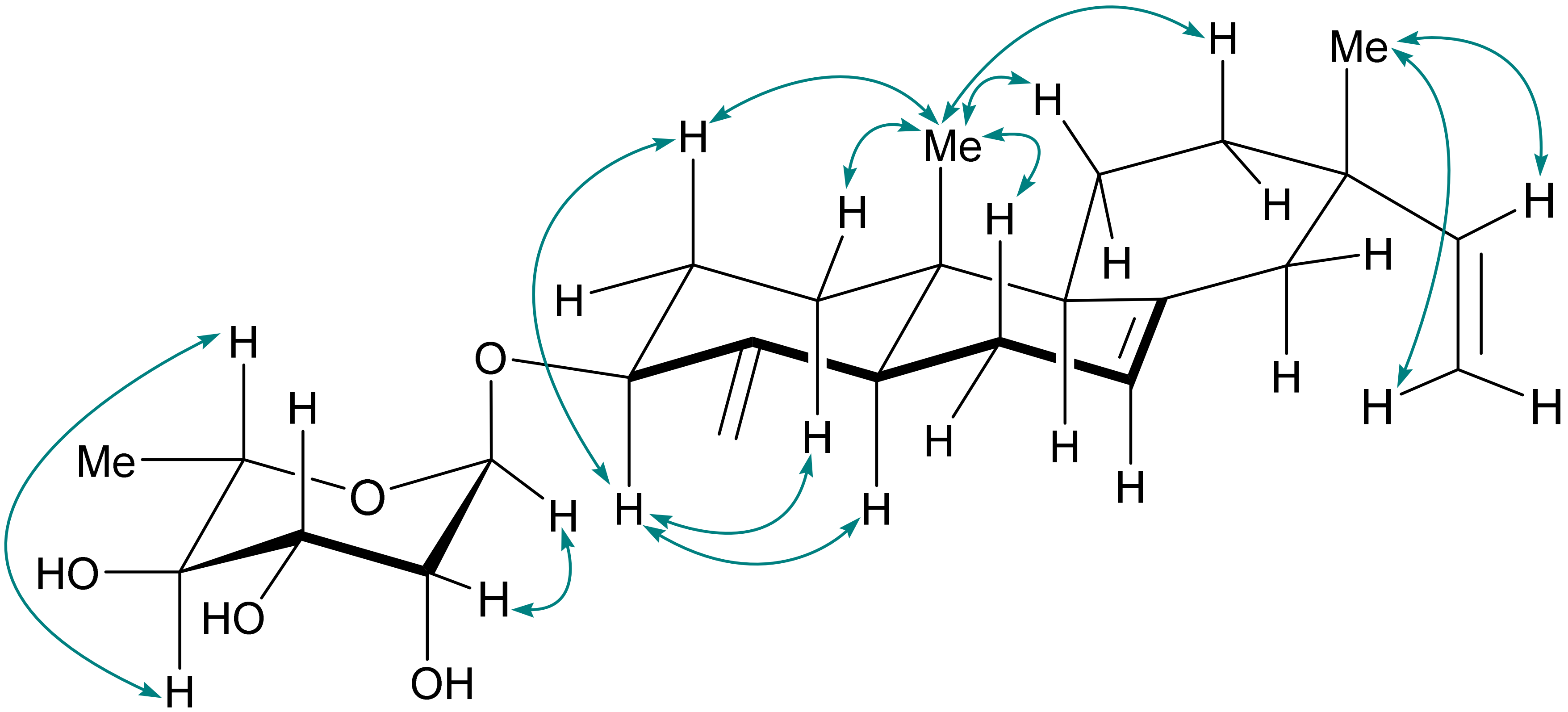

**Figure S1**: [1]H-NMR spectrum of 3-*O*-$\alpha$-L-rhamnopyranosyl-18-Norisopimara-4(19),7,15-triene **(7)** in $CDCl_3$, 600 MHz

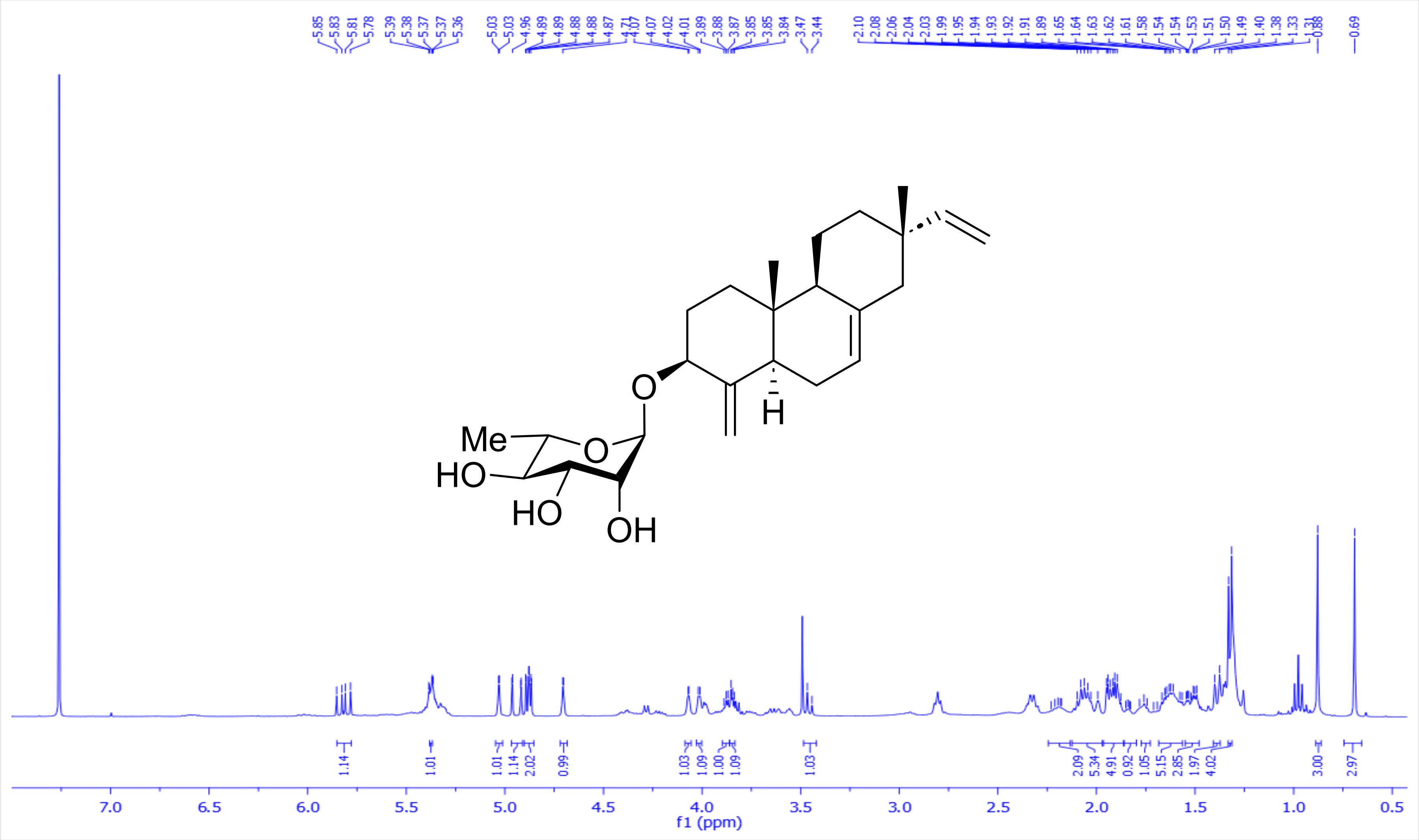

**Figure S2**: [1]H-[1]H COSY spectrum of 3-*O*-$\alpha$-L-rhamnopyranosyl-18-Norisopimara-4(19),7,15-triene **(7)** in $CDCl_3$, 600 MHz

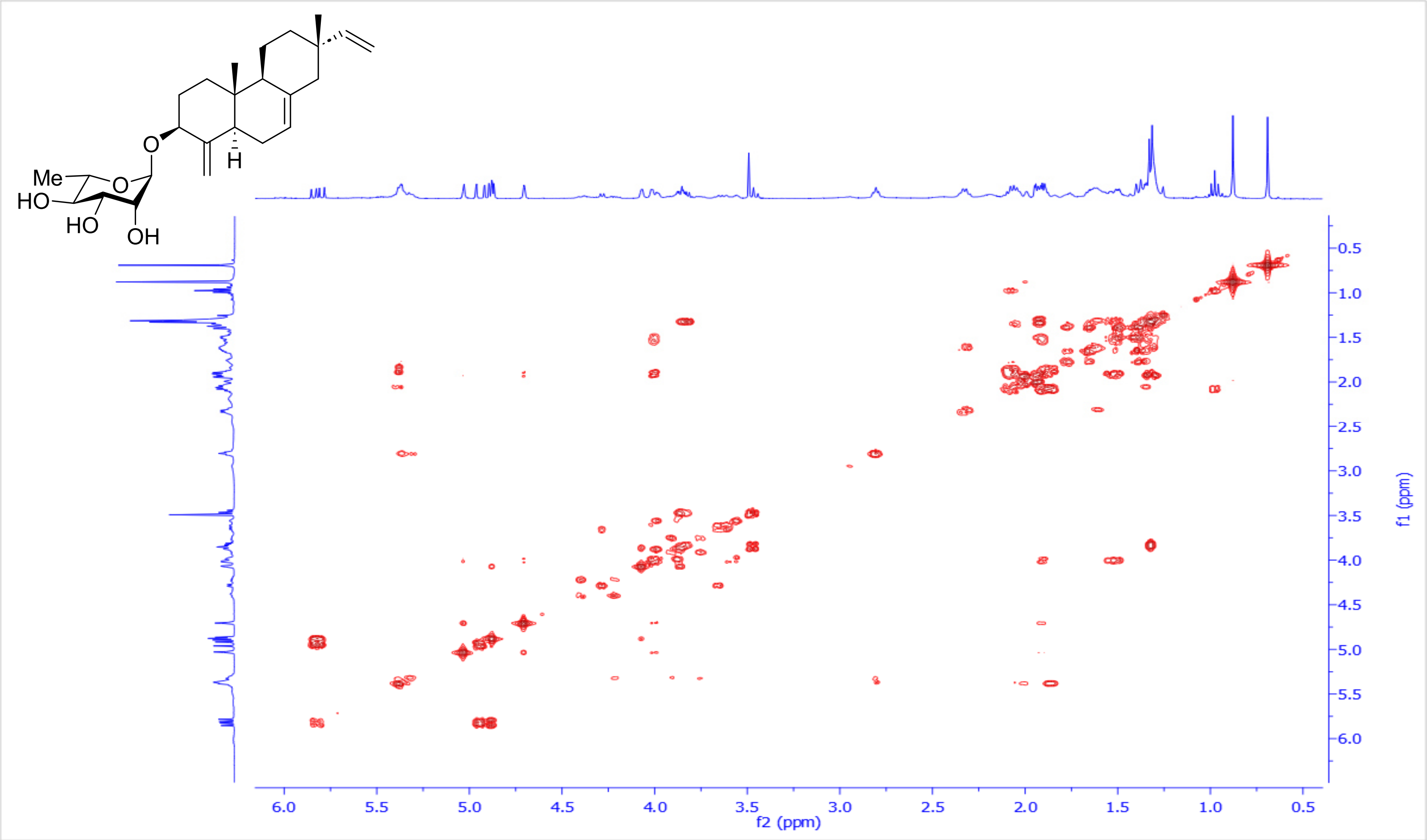

**Figure S3**: $^{1}$H-$^{13}$C HSQC spectrum of 3-*O*-*α*-L-rhamnopyranosyl-18-Norisopimara-4(19),7,15-triene **(7)** in $CDCl_3$, 600 and 150 MHz

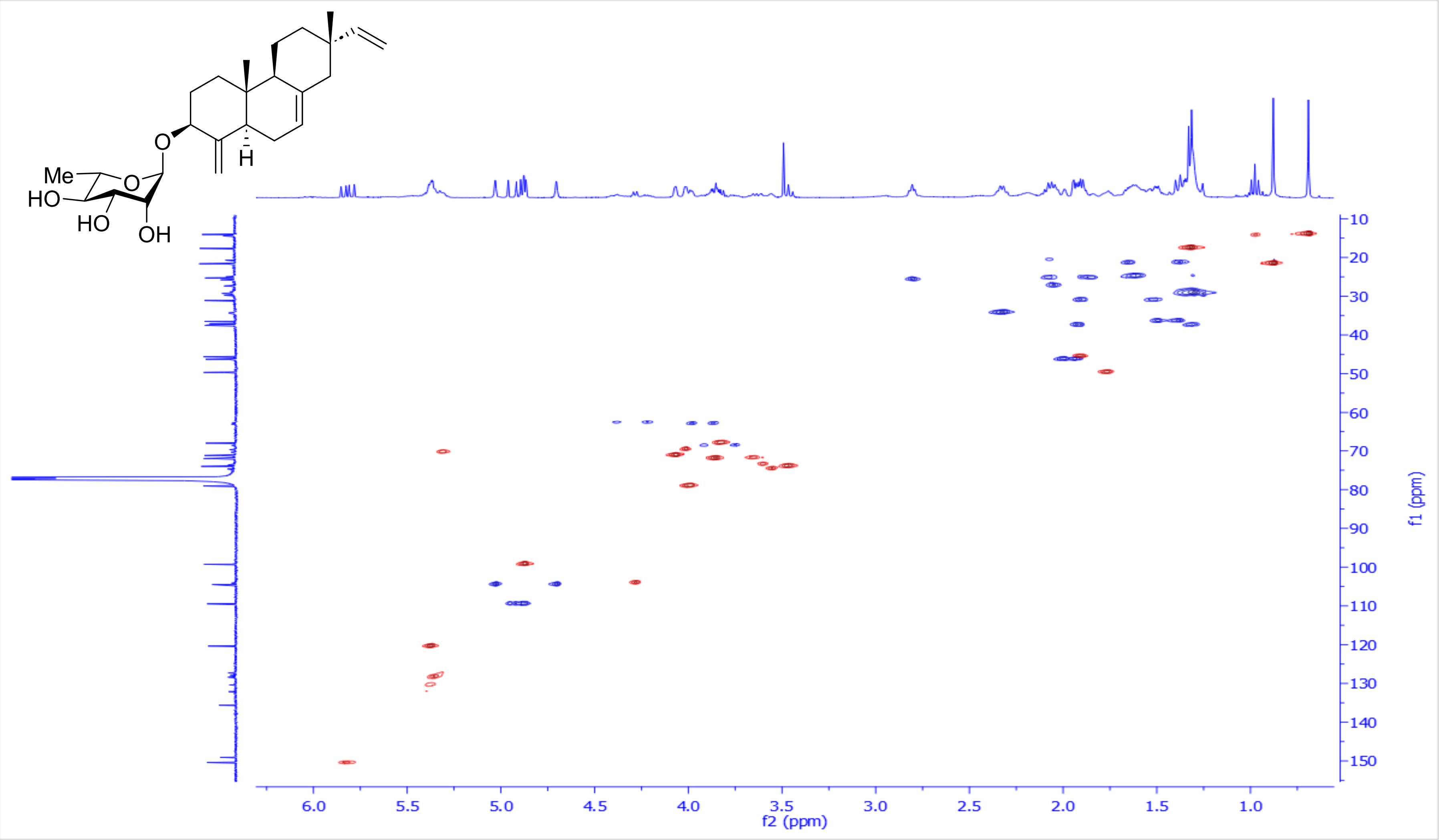

**Figure S4**: $^{1}$H-$^{13}$C HMBC spectrum of 3-*O*-*α*-L-rhamnopyranosyl-18-Norisopimara-4(19),7,15-triene **(7)** in $CDCl_3$, 600 and 150 MHz

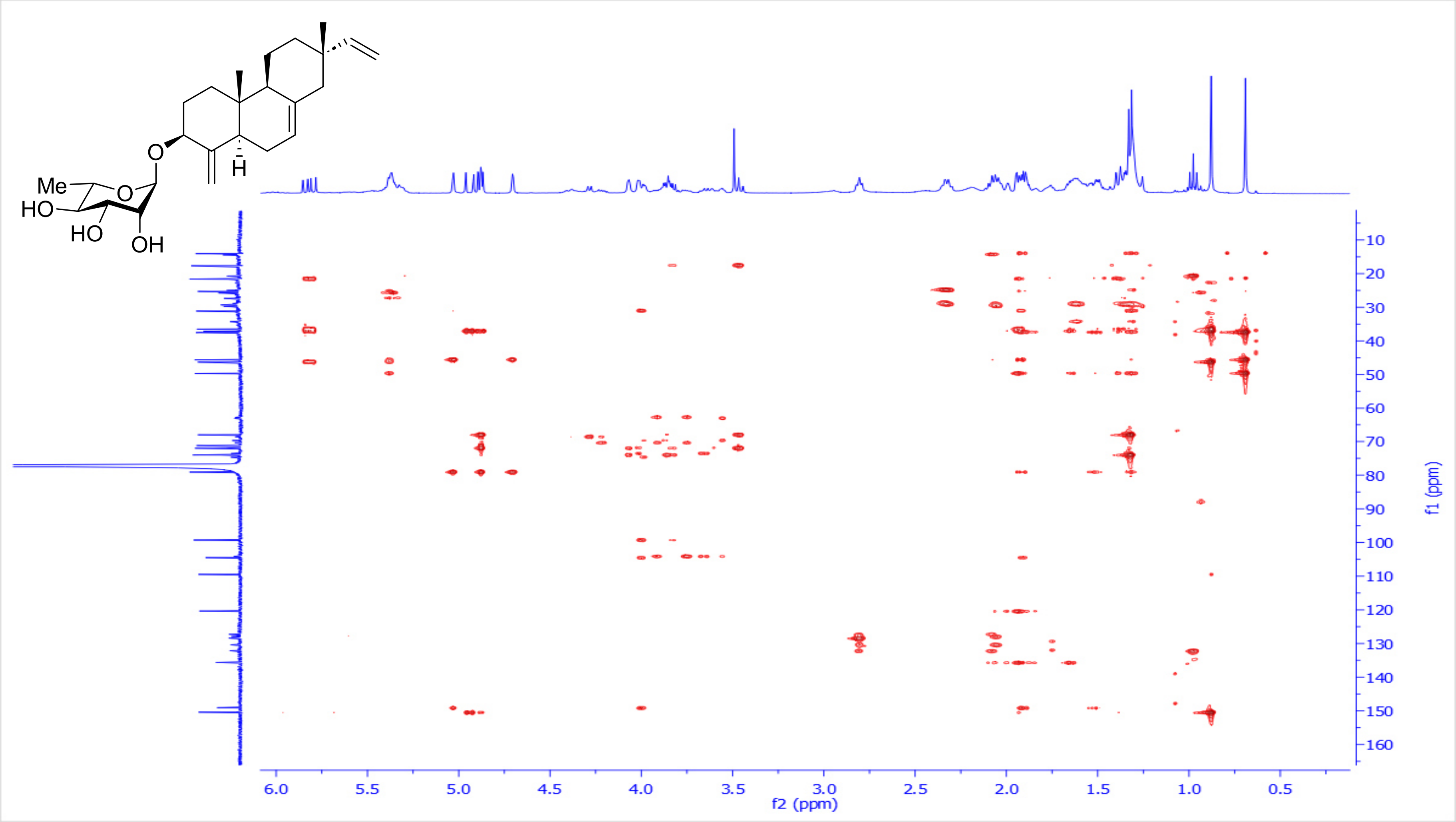

**Figure S5**: $^1$H-$^1$H ROESY spectrum of 3-*O*-*α*-L-rhamnopyranosyl-18-Norisopimara-4(19),7,15-triene **(7)** in $CDCl_3$, 600 MHz

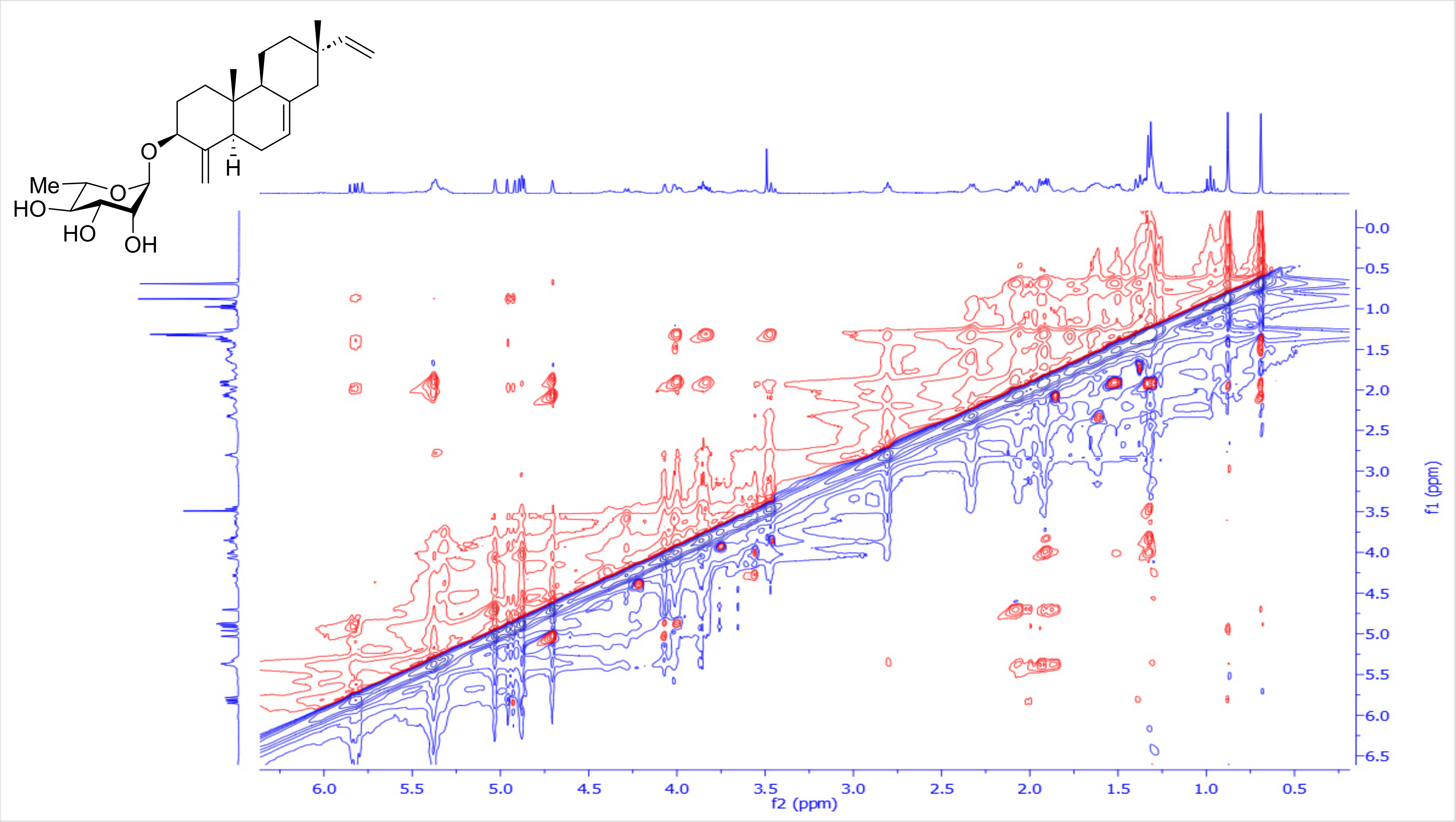

**Figure S6**: $^{13}$C-NMR spectrum of 3-*O*-*α*-L-rhamnopyranosyl-18-Norisopimara-4(19),7,15-triene **(7)** in $CDCl_3$, 150 MHz

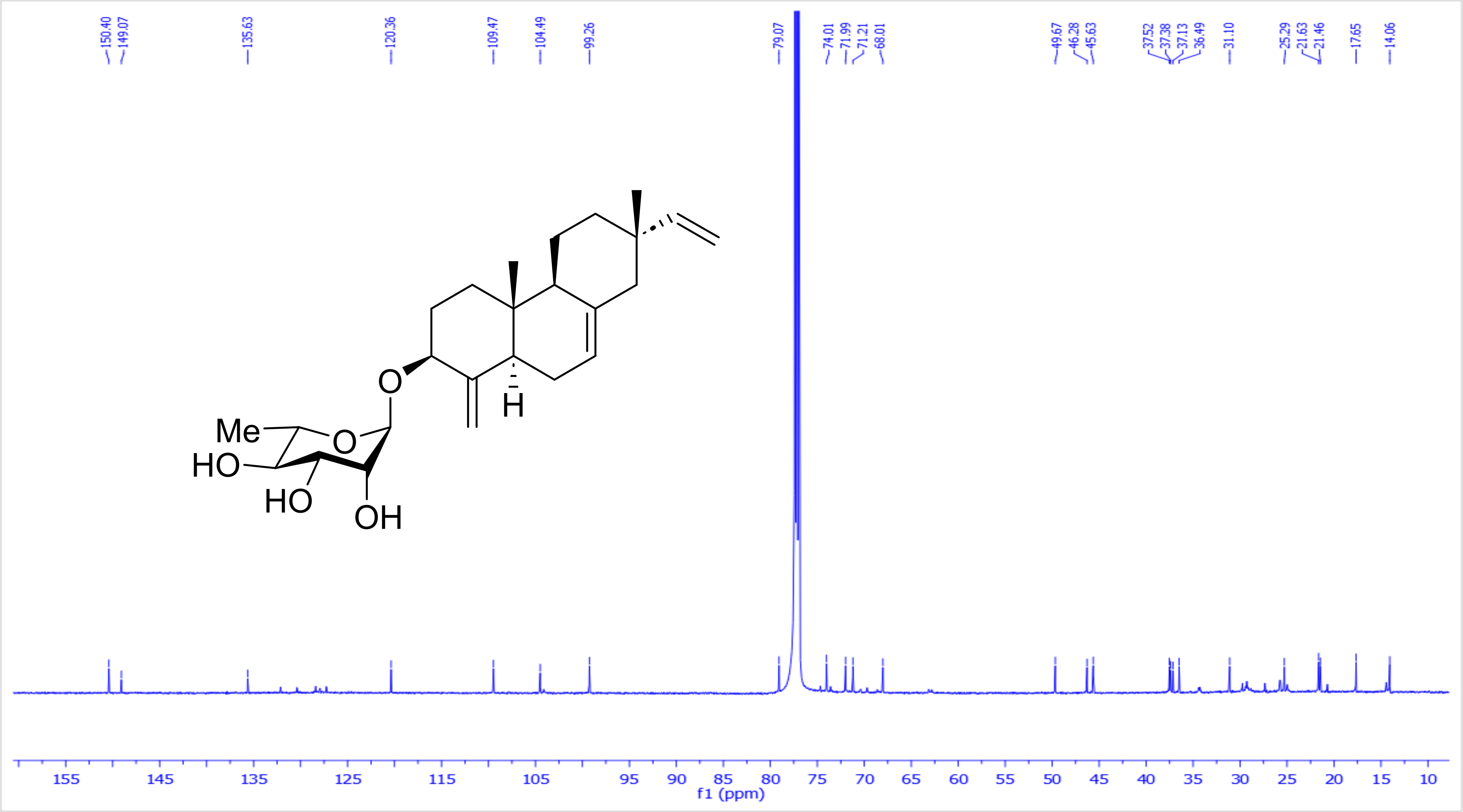

**Figure S7:** HR-MS (+ve) for 3-*O*-$\alpha$-L-rhamnopyranosyl-18-Norisopimara-4(19),7,15-triene **(7)**

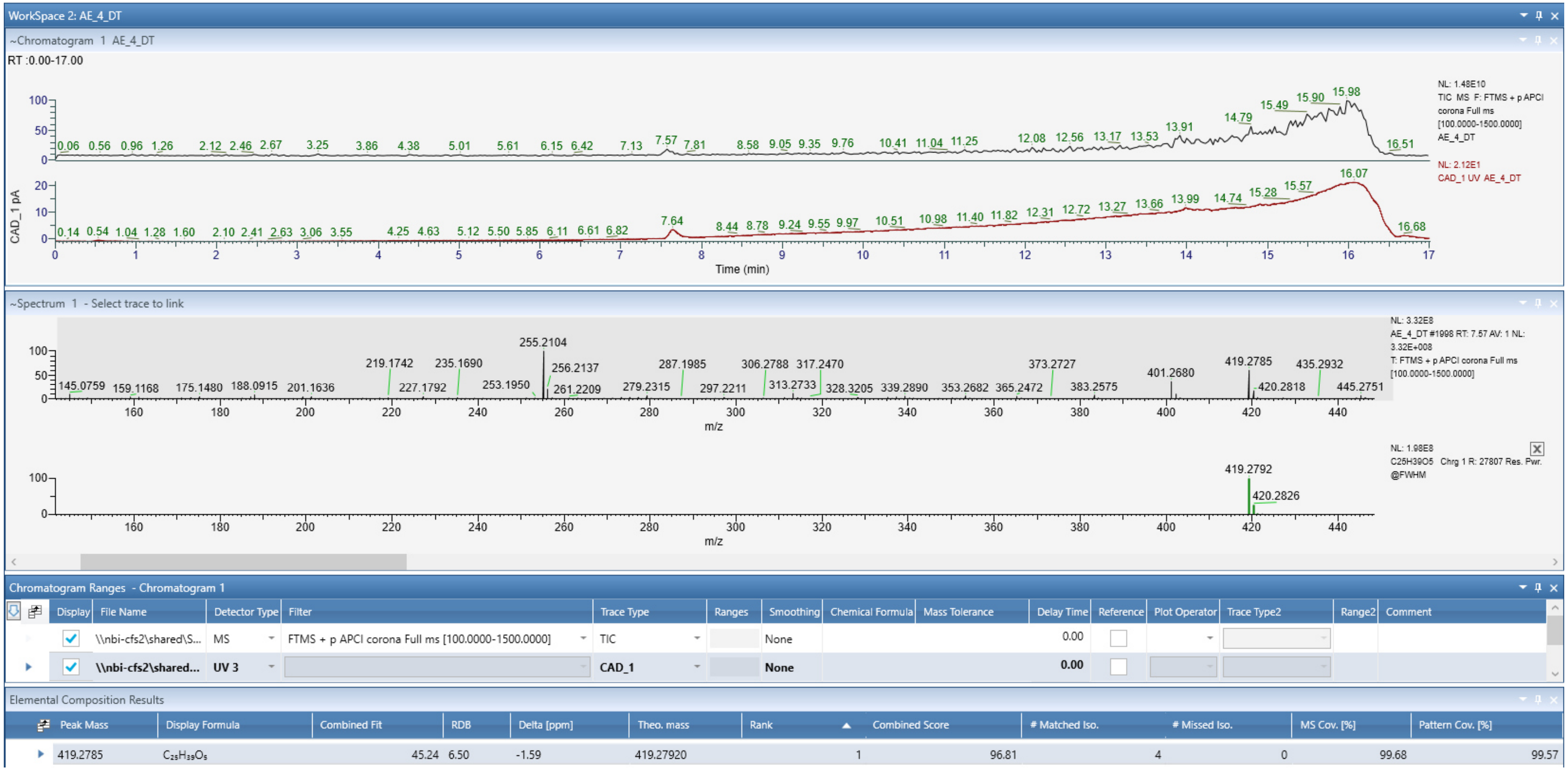

**Table 2**: [1]H, [13]C NMR spectroscopic data for 3-*O*-*α*-L-arabinopyranosyl-18-Norisopimara-4(19),7,15-triene **(8)** in $CDCl_3$, 600 and 150 MHz.

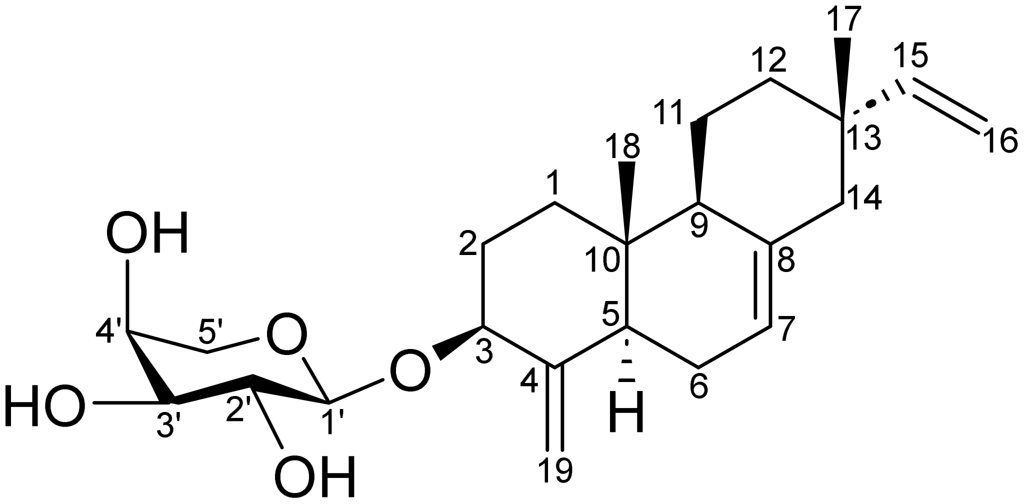

| **Position** | ***δ*C, Type** | ***δ*H, mult. (*J* in Hz)** |
|---|---|---|
| **1** | 37.4, $CH_2$ | 1.93/1.32, m |
| **2** | 31.3, $CH_2$ | 1.96/1.56, m |
| **3** | 79.6, CH | 4.08, m |
| **4** | 148.7, Cq | - |
| **5** | 45.7, CH | 1.90, m |
| **6** | 25.8, $CH_2$ | 2.08/1.85 |
| **7** | 120.3, CH | 5.38, m |
| **8** | 135.8, Cq | - |
| **9** | 49.9, CH | 1.76, m |
| **10** | 37.5, Cq | - |
| **11** | 21.5, $CH_2$ | 1.65/1.39, m |
| **12** | 36.5, $CH_2$ | 1.50/1.40, m |
| **13** | 37.1, Cq | - |
| **14** | 46.2, $CH_2$ | 2.00/1.94, m |
| **15** | 150.4, CH | 5.82, *dd* (17.5, 10.7) |
| **16** | 109.5, $CH_2$ | 4.94, *dd* (17.5, 1.2) |
| | | 4.88, *dd* (10.7, 1.2) |
| **17** | 21.6, $CH_3$ | 0.88, *s* |
| **18** | 14.1, $CH_3$ | 0.69, s |
| **19** | 104.9, $CH_2$ | 5.19/4.74, br s |
| **Ara*p*-1′** | 101.5, CH | 4.55, *d* (5.7) |
| **Ara*p*-2′** | 72.5, CH | 3.59, m |
| **Ara*p*-3′** | 74.4, CH | 3.63, *t* (7.3) |
| **Ara*p*-4′** | 69.9, CH | 3.78, m |
| **Ara*p*-5′** | 64.1, $CH_2$ | 4.08, m/3.38, *dd* (11.9, 7.8) |

**Scheme 3**: Key $^{1}H$-$^{1}H$ COSY (**bold blue**), $^{1}H$-$^{13}C$ HMBC (H→C, red arrows) observed for 3-*O*-*α*-L-arabinopyranosyl-18-Norisopimara-4(19),7,15-triene **(8)**

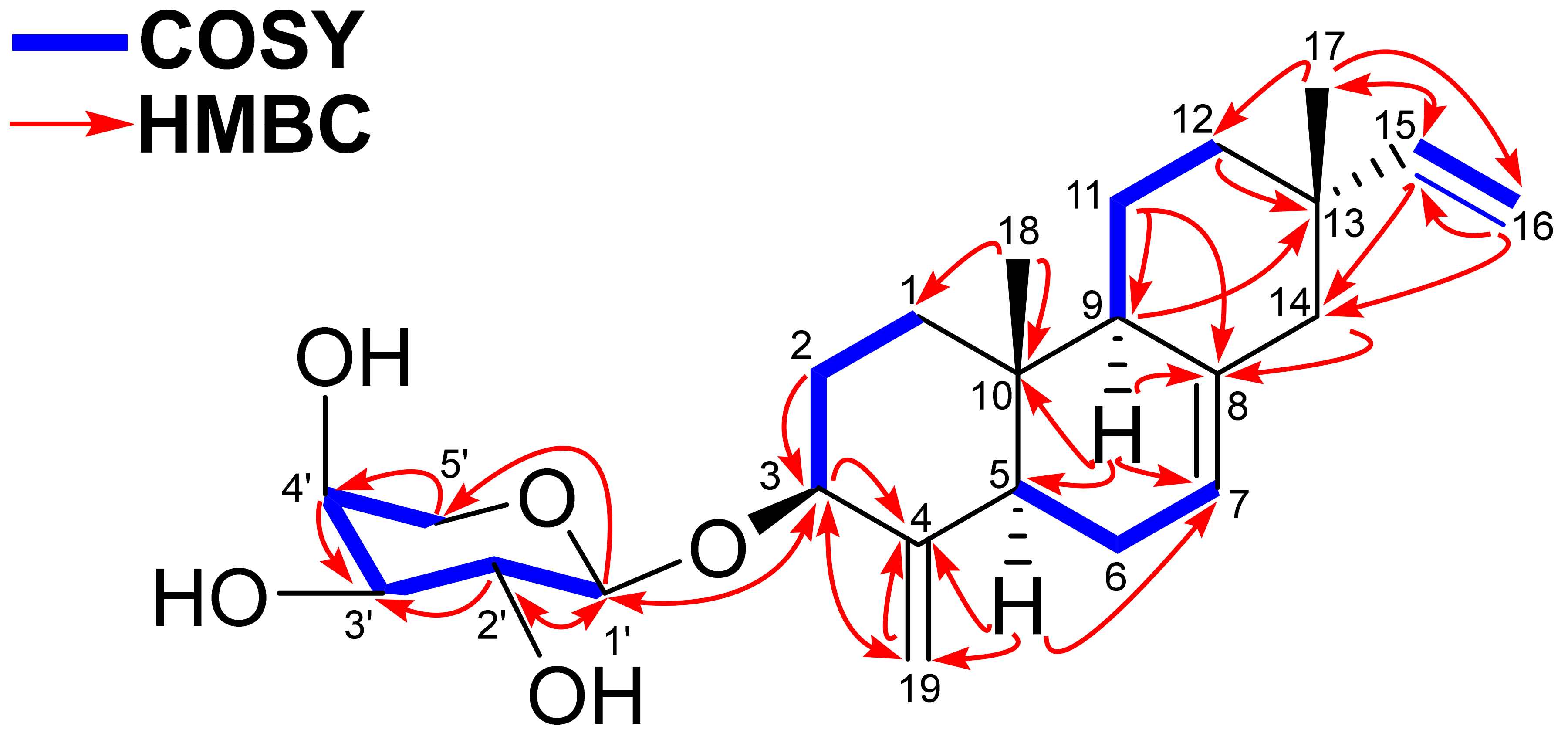

**Scheme 4**: Key ¹H-¹H ROESY (H↔H, green) observed for 3-*O*-*α*-L-arabinopyranosyl-18-Norisopimara-4(19),7,15-triene **(8)**

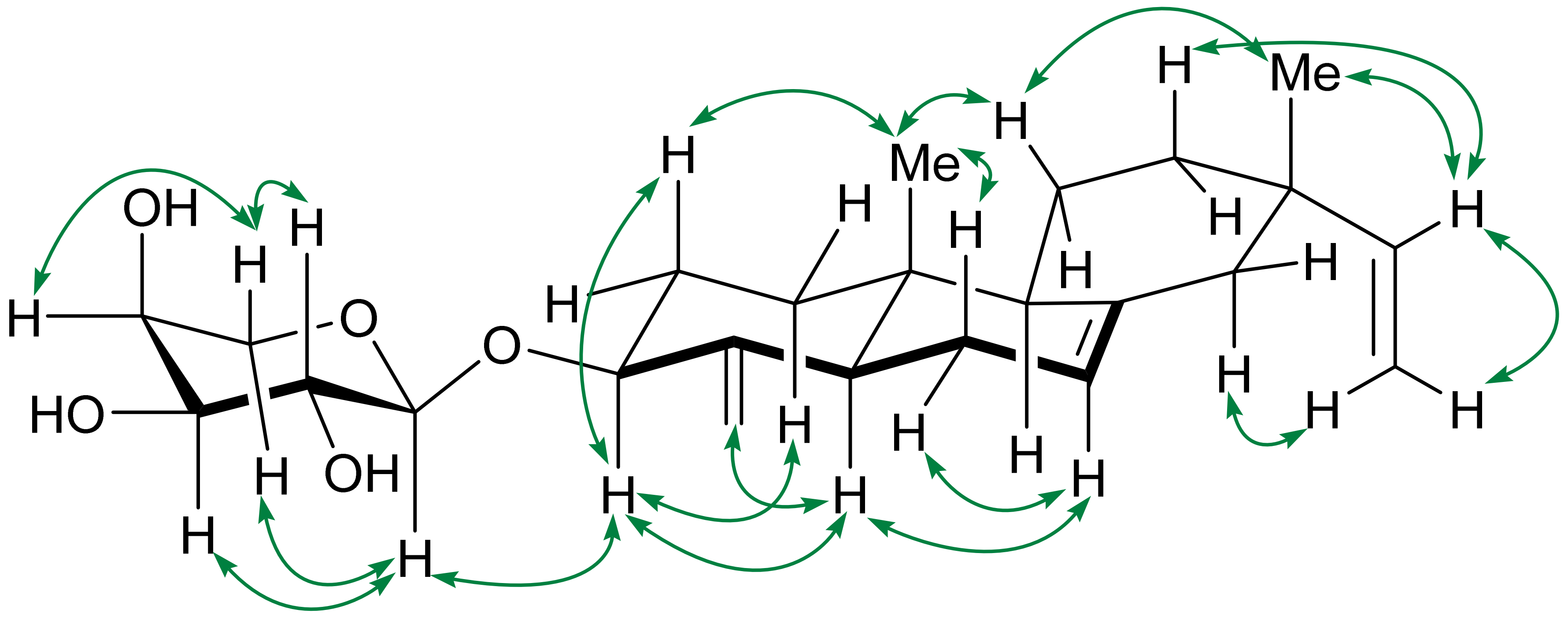

**Figure S8**: [1]H-NMR spectrum of 3-*O*-*α*-L-arabinopyranosyl-18-Norisopimara-4(19),7,15-triene **(8)** in $CDCl_3$, 600 MHz

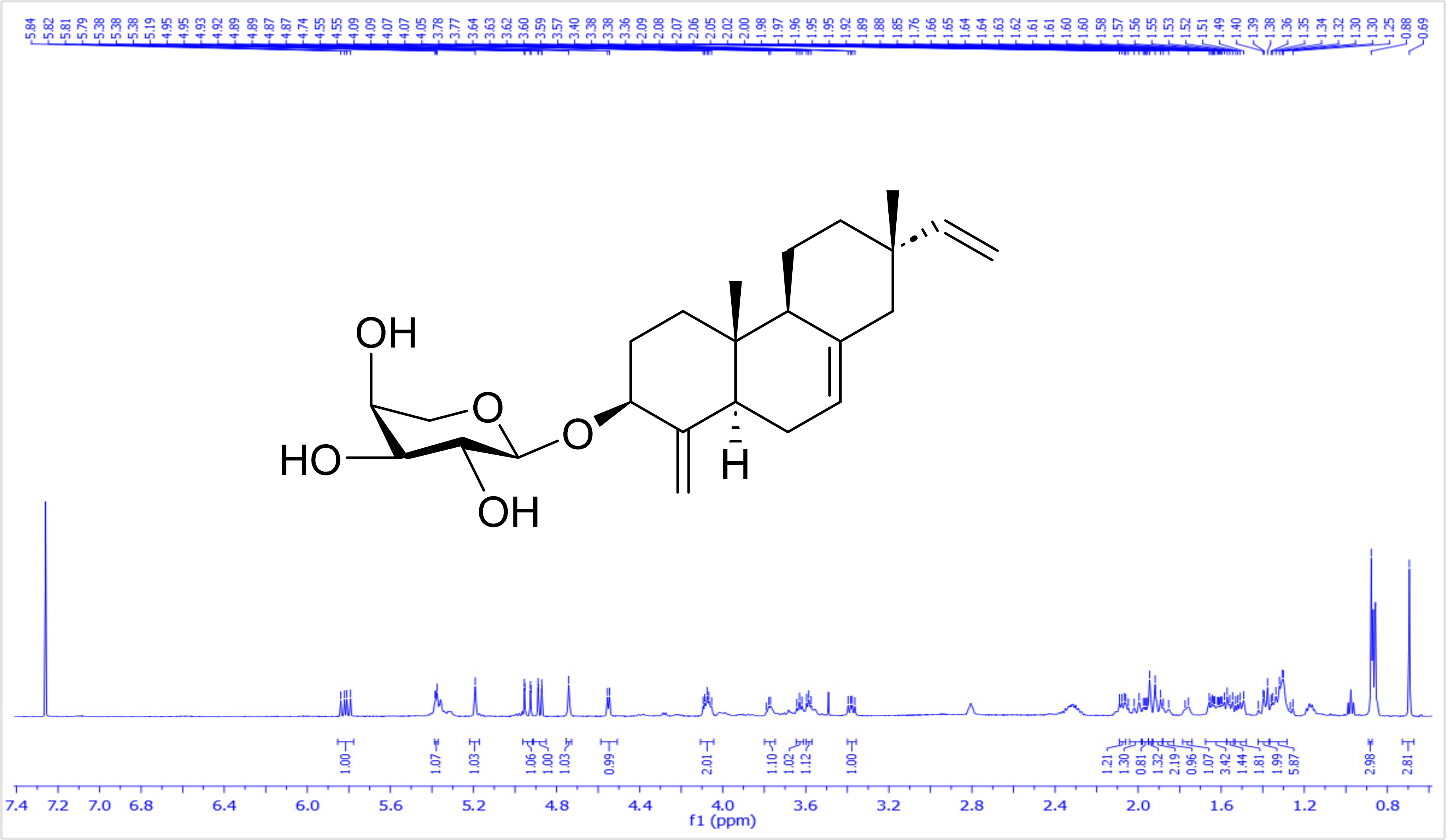

**Figure S9**: [1]H-[1]H COSY spectrum of 3-*O*-*α*-L-arabinopyranosyl-18-Norisopimara-4(19),7,15-triene **(8)** in CDCl₃, 600 MHz

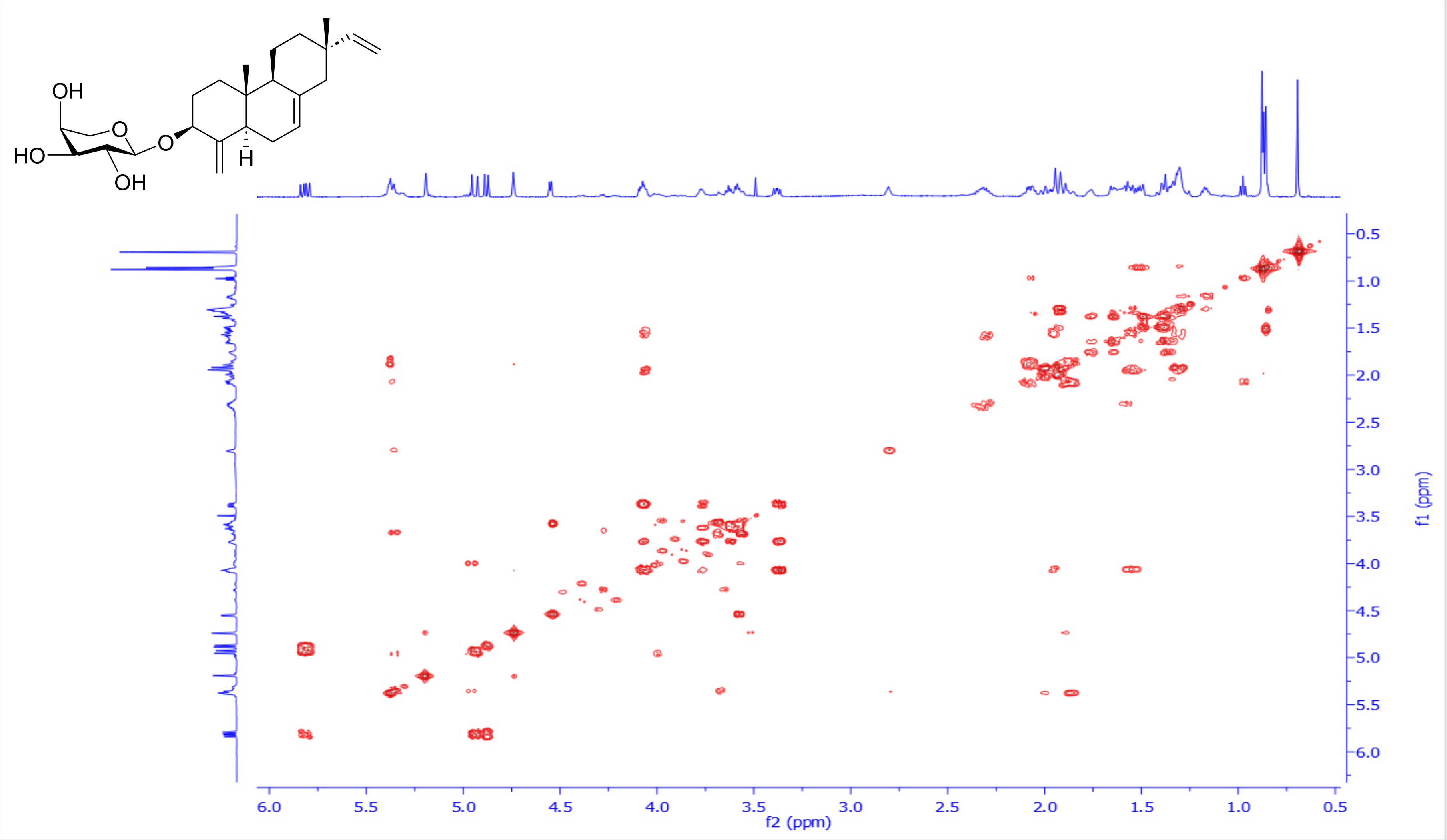

**Figure S10**: $^{1}H$-$^{13}C$ HSQC spectrum of 3-*O*-*α*-L-arabinopyranosyl-18-Norisopimara-4(19),7,15-triene **(8)** in $CDCl_3$, 600 and 150 MHz

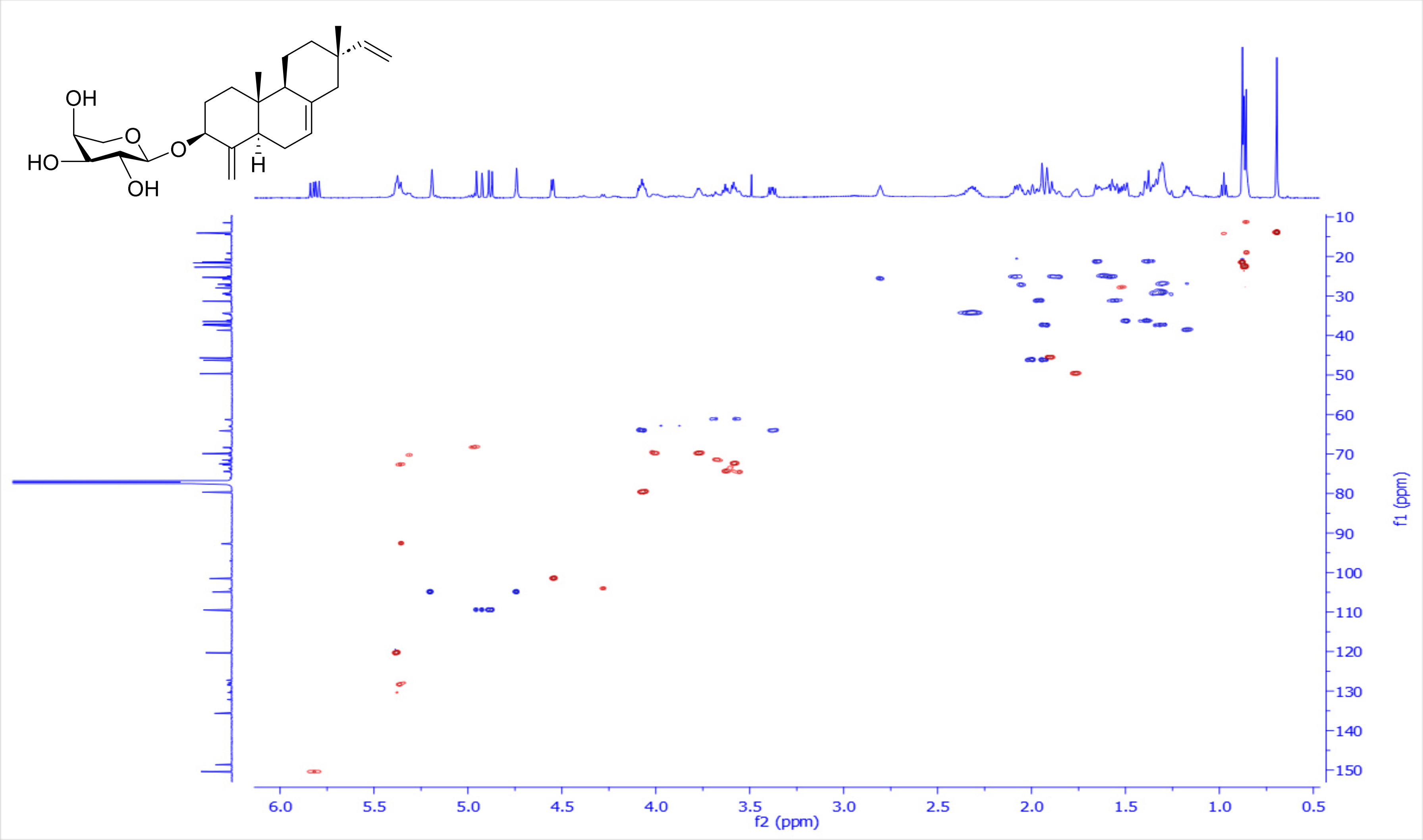

**Figure S11**: $^{1}H$-$^{13}C$ HMBC spectrum of 3-*O*-*α*-L-arabinopyranosyl-18-Norisopimara-4(19),7,15-triene **(8)** in $CDCl_3$, 600 and 150 MHz

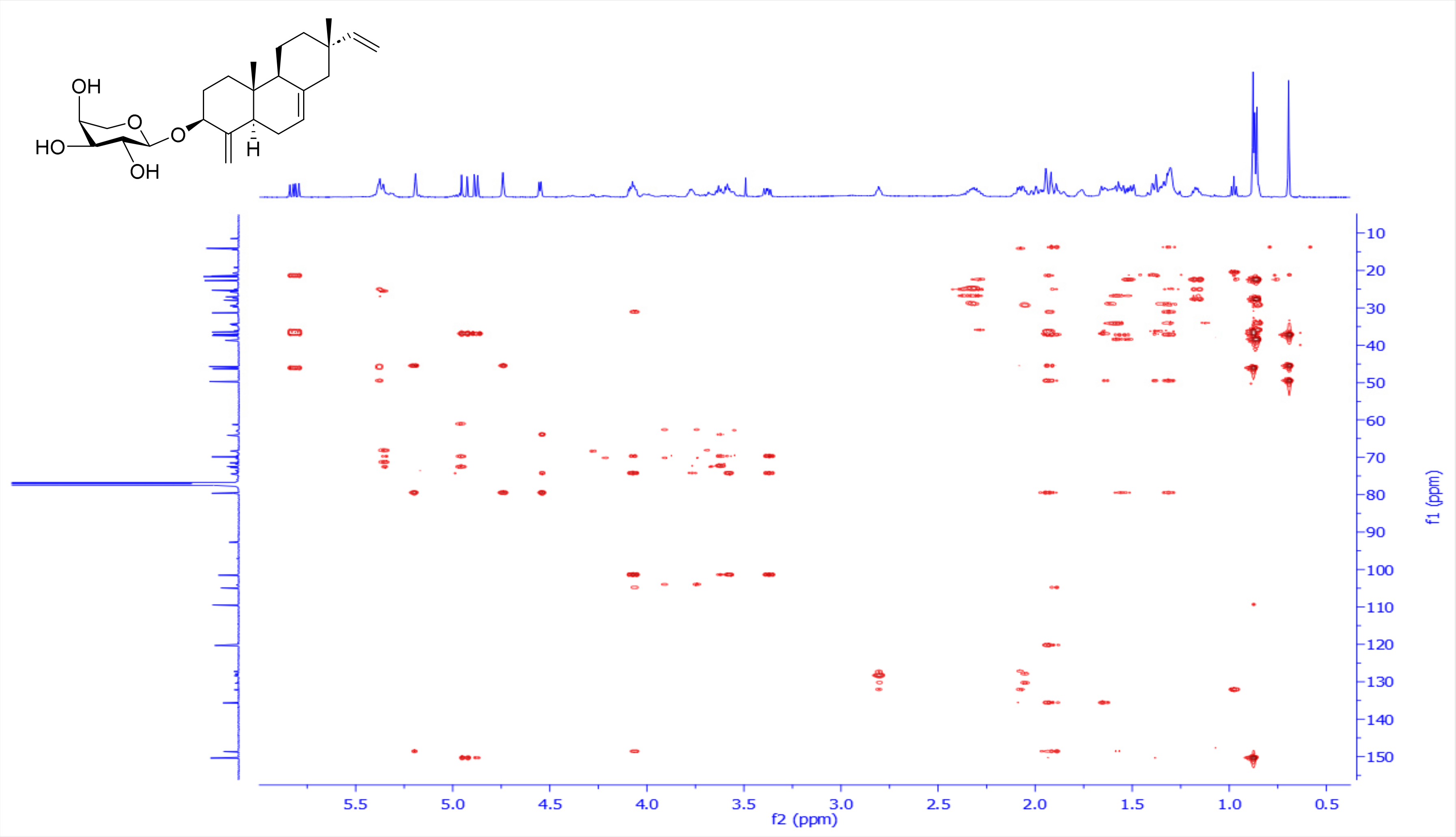

**Figure S12**: $^1$H-$^1$H ROESY spectrum of 3-*O*-*α*-L-arabinopyranosyl-18-Norisopimara-4(19),7,15-triene **(8)** in CDCl$_3$, 600 MHz

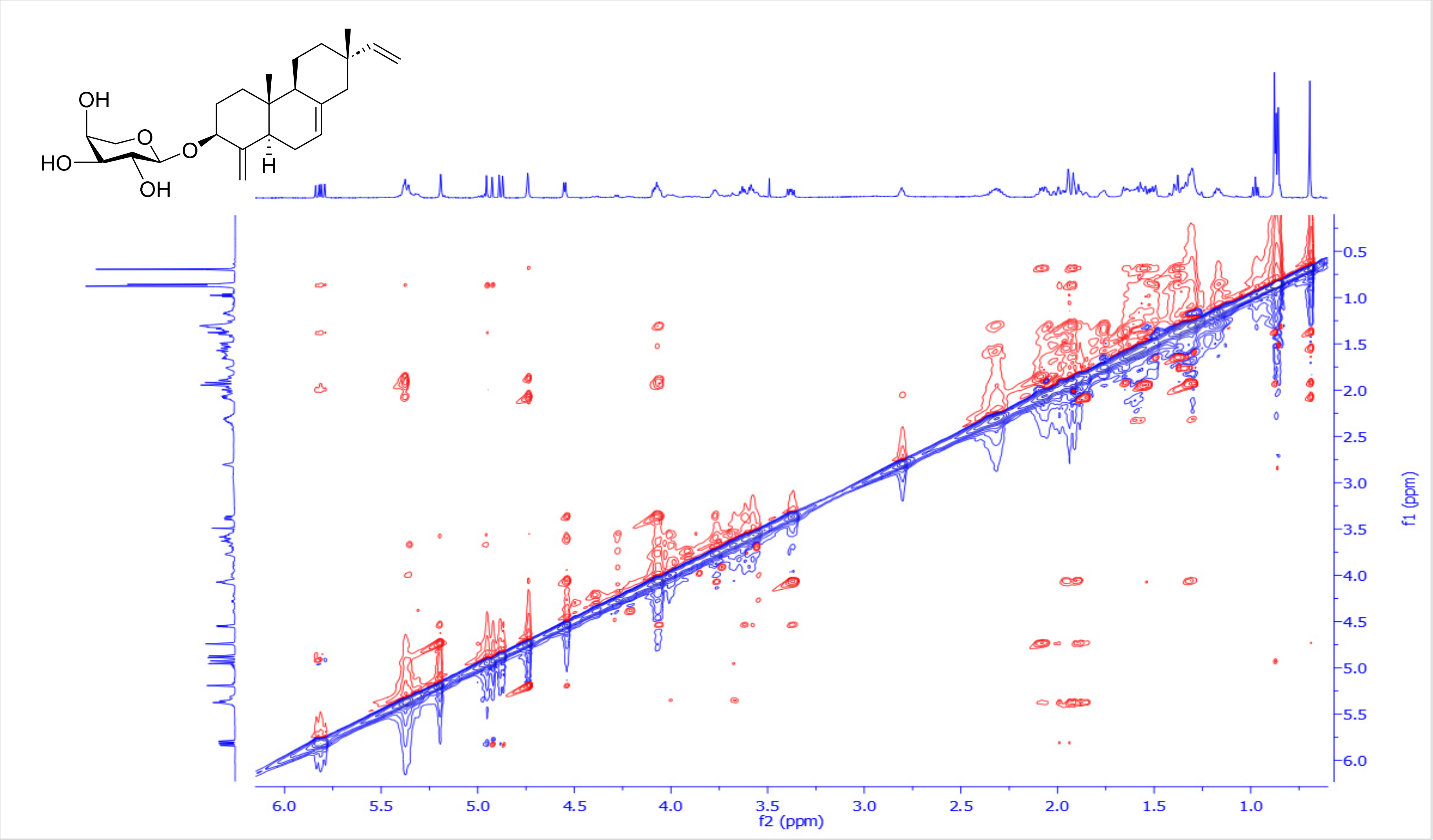

**Figure S13**: $^{13}$C-NMR spectrum of 3-*O*-$\alpha$-L-arabinopyranosyl-18-Norisopimara-4(19),7,15-triene **(8)** in $CDCl_3$, 150 MHz

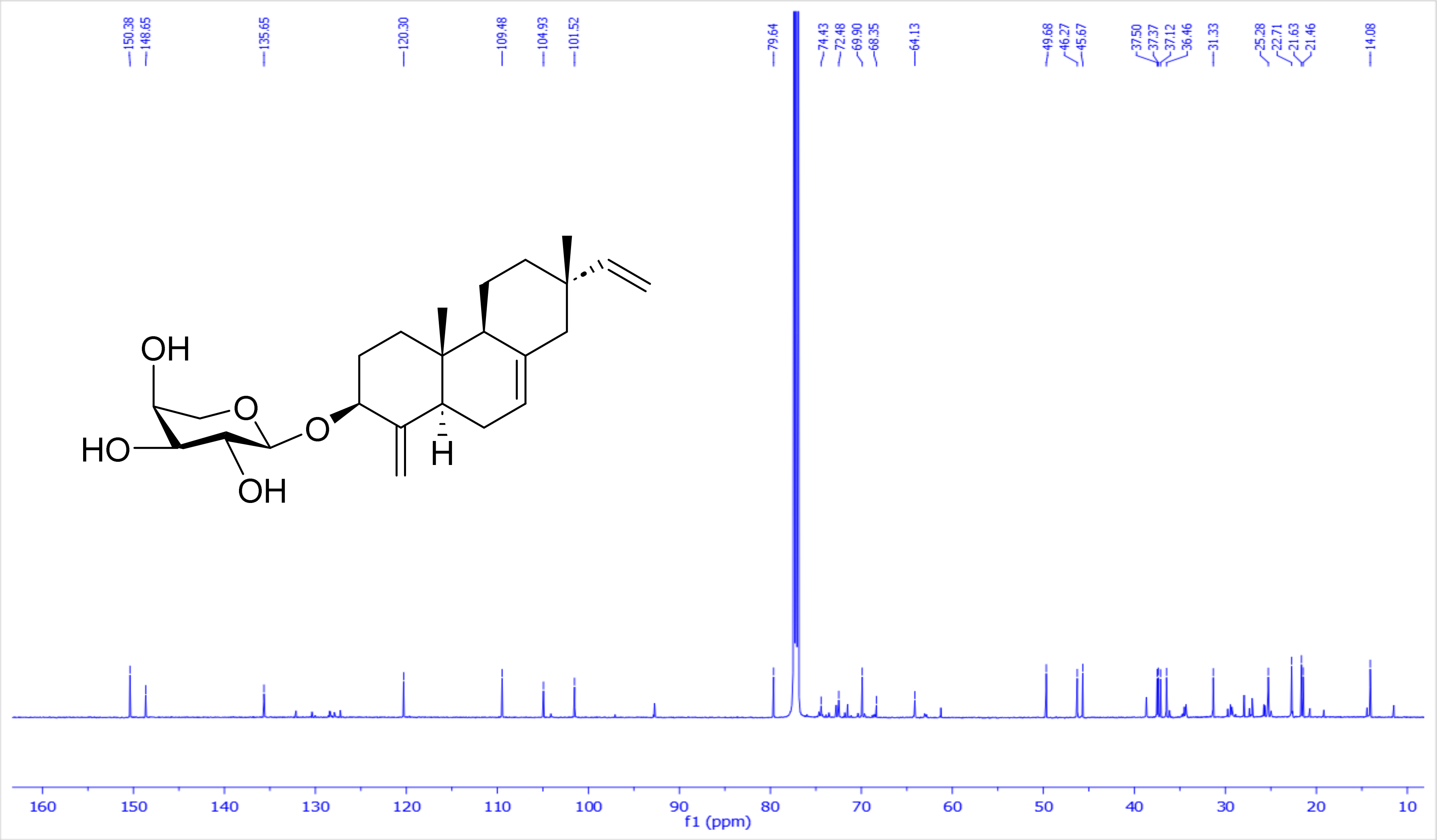

**Figure S14:** HR-MS (+ve) for 3-*O*-$\alpha$-L-arabinopyranosyl-18-Norisopimara-4(19),7,15-triene **(8)**

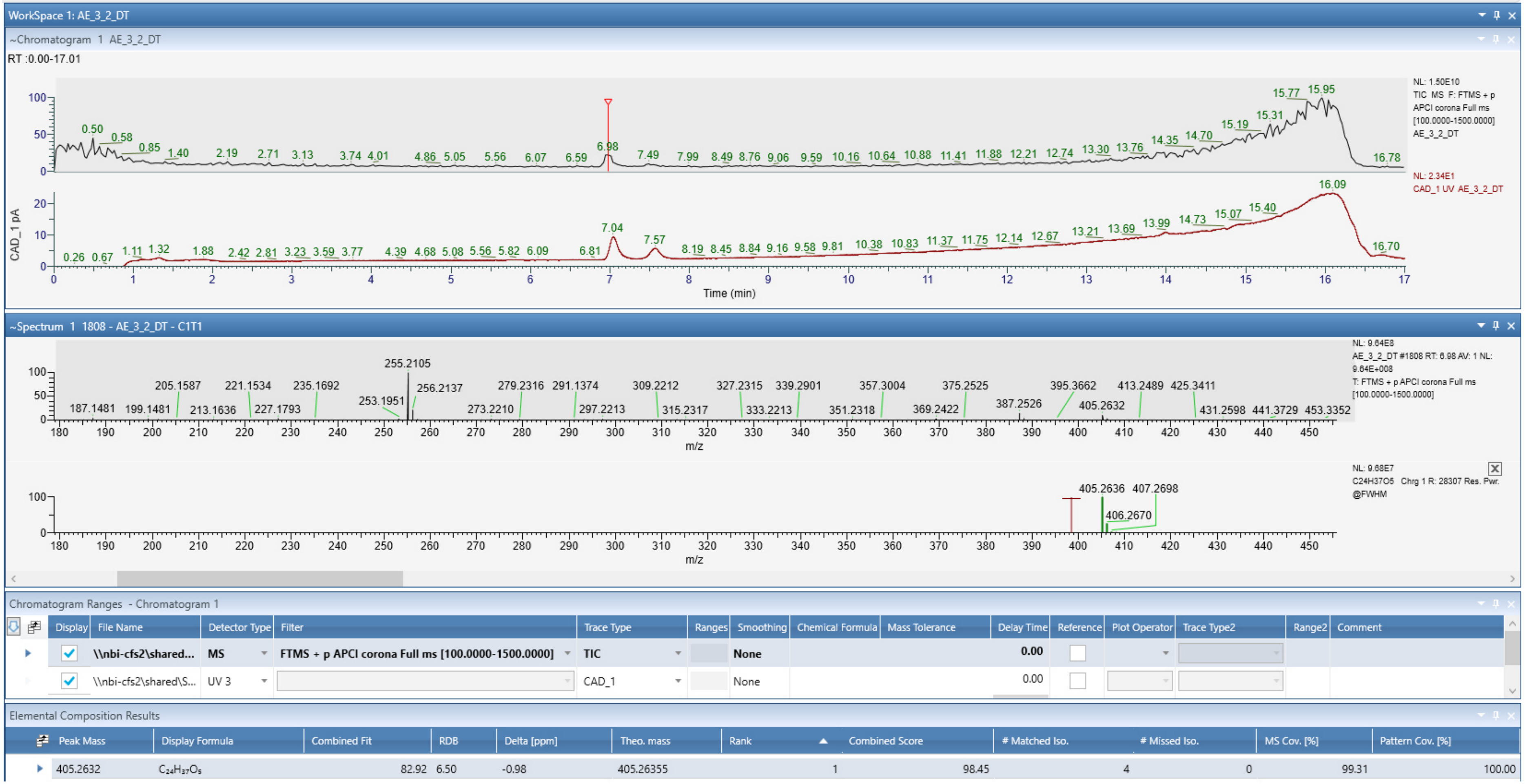

**2- Extraction, purification and structural characterization of *ent*-pimara-8(14), 15-diene-19-ol (11)**

*TaCPS_B1, TaKSL_B2* and *TaCYP701A64_B2* were transiently co-expressed in **75** *Nicotiana benthamiana* plants to afford **60 g** of dry leaves. Subsequently, it was exhaustively extracted using ethyl acetate via speed extractor. The organic extract was collected and dried under reduced pressure to afford an intense green viscus material. Afterwards, it was introduced to normal phase flash chromatography using a linear gradient of hexane/ethyl acetate [100/0 up 45/55] along 30 minutes, affording 12 subfractions F1-F12, where all have been monitored by LC-MS. Promising fraction F3 was further purified using repetitive $C_{18}$ semi-preparative HPLC using a short gradient of water/acetonitrile [15/85 up 0/100], 4 mL/min, acidified with 0.1 % FA along 22 min to finally afford *ent*-pimara-8(14), 15-diene-19-ol (**11**) (**4.3** mg, white powder). Its full structure was resolved based on extensive 1D & 2D-NMR data analysis.

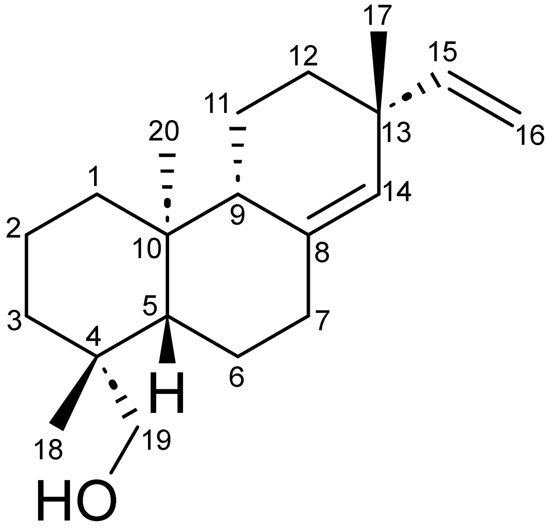

***ent*-pimara-8(14), 15-diene-19-ol (11)**

**Table 3**: [1]H, [13]C NMR spectroscopic data for *ent*-pimara-8(14), 15-diene-19-ol (**11**) in $CDCl_3$, 600 and 150 MHz.

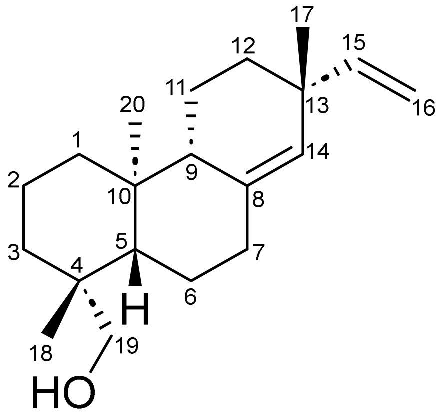

| Position | $\delta_C$, Type | $\delta_H$, mult. ($J$ in Hz) |
|---|---|---|
| **1** | 39.3, $CH_2$ | 1.67/1.04, *m* |
| **2** | 18.70, $CH_2$ | 1.44/1.03, *m* |
| **3** | 35.6, $CH_2$ | 1.84/0.96, *m* |
| **4** | 38.5, Cq* | - |
| **5** | 55.9, CH | 1.21, *m* |
| **6** | 22.6, $CH_2$ | 1.68/1.34, *m* |
| **7** | 36.3, $CH_2$ | 2.32, *ddd* (14.3, 4.5, 2) |
| | | 2.03, *td* (13.7, 5.7) |
| **8** | 138.3, Cq | - |
| **9** | 51.7, CH | 1.69, *m* |
| **10** | 39.1, Cq | - |
| **11** | 19.4, $CH_2$ | 1.51/1.31, *m* |
| **12** | 35.9, $CH_2$ | 1.54/1.21, *m* |
| **13** | 38.7, Cq* | - |
| **14** | 128.3, CH | 5.13, *d* (1.5) |
| **15** | 147.5, CH | 5.71, *dd* (17.5, 10.4) |
| **16** | 113.0, $CH_2$ | 4.95, *dd* (10.4, 1.9) |
| | | 4.90, *dd* (17.3, 1.9) |
| **17** | 29.6, $CH_3$ | 0.99, *s* |
| **18** | 27.2, $CH_3$ | 0.99, s |
| **19** | 65.4, $CH_2$ | 3.82, *d* (10.8) |
| | | 3.44, *d* (10.8) |
| **20** | 16.0, $CH_3$ | 0.70, *s* |

*Chemical shifts may be interchanged

**Scheme 5**: Key ¹H-¹H COSY (**bold blue**), ¹H-¹³C HMBC (H→C, red arrows) observed for *ent*-pimara-8(14), 15-diene-19-ol (**11**)

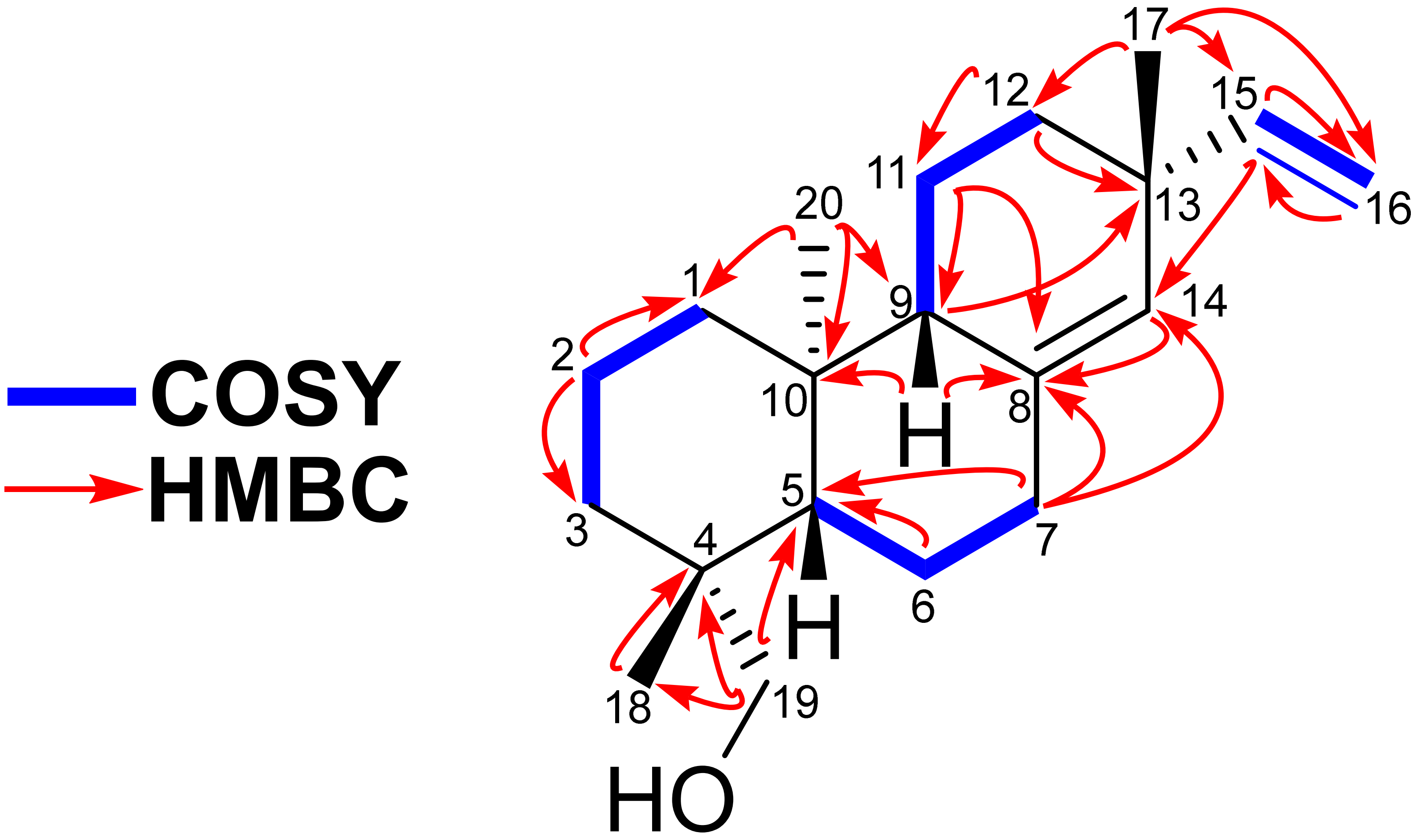

**Scheme 6**: Key $^1H$-$^1H$ ROESY (H↔H, green) observed for *ent*-pimara-8(14), 15-diene-19-ol (**11**)

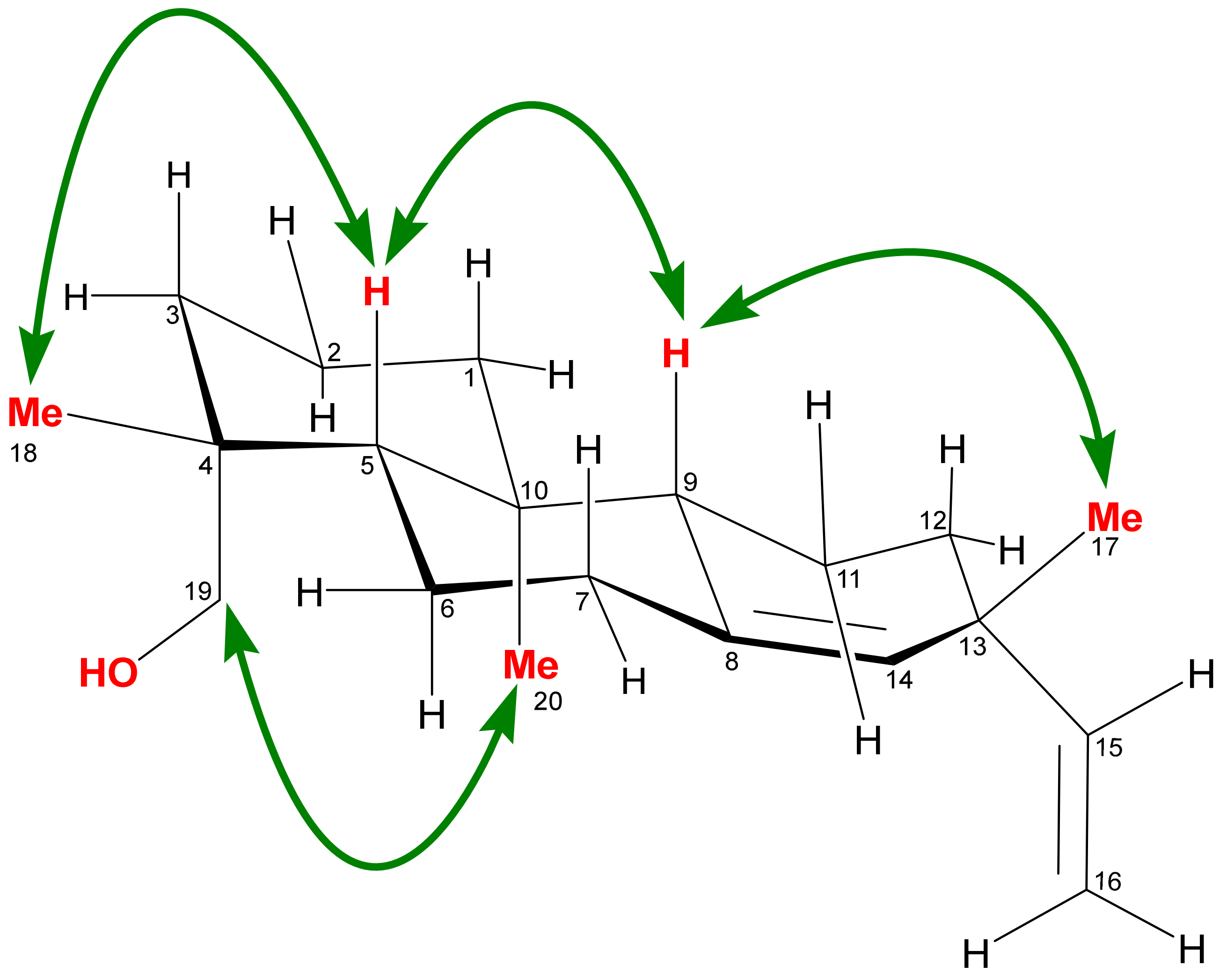

**Figure S15**: [1]H-NMR spectrum of *ent*-pimara-8(14), 15-diene-19-ol (**11**) in $CDCl_3$, 600 MHz

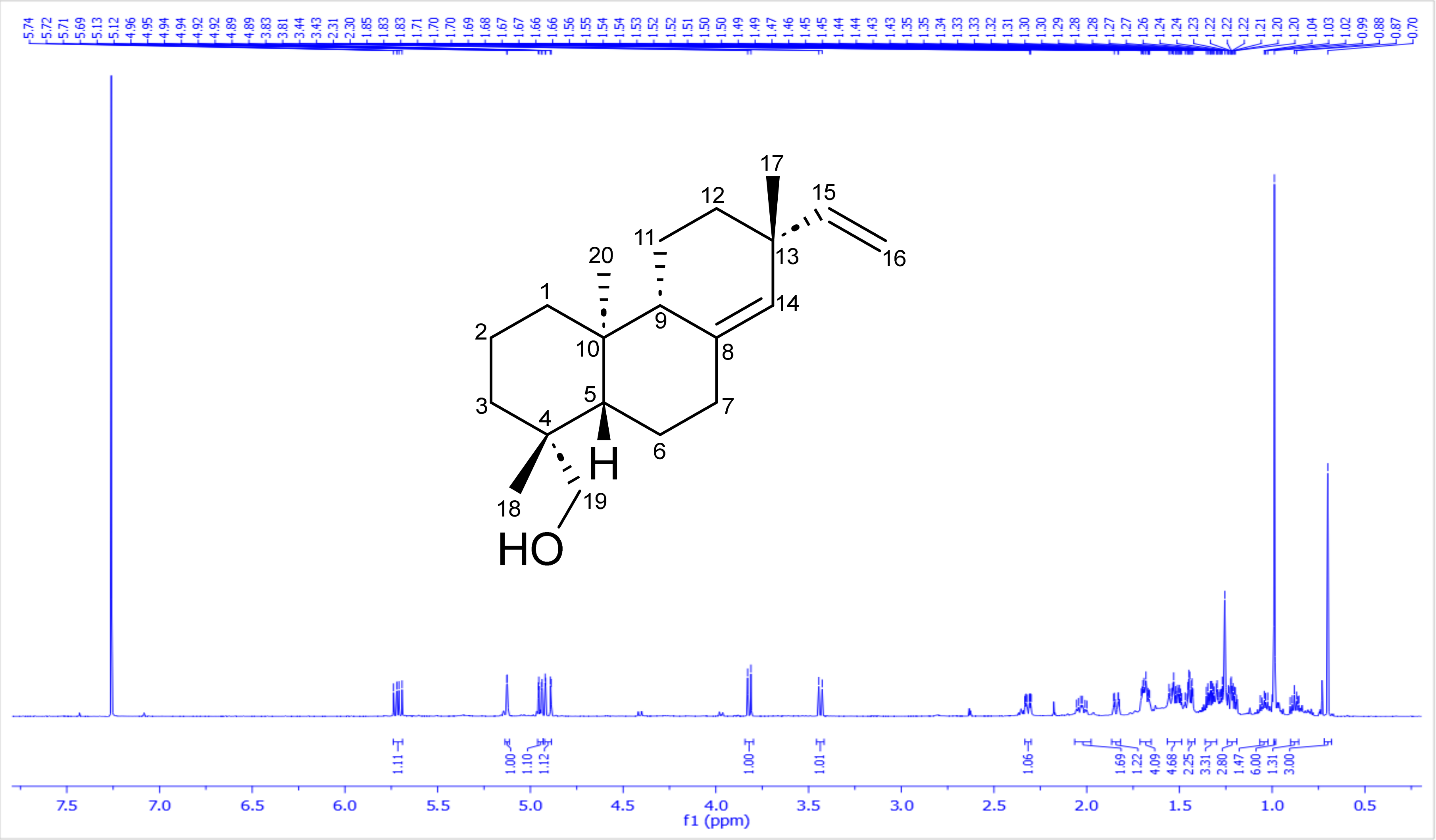

**Figure S16**: $^1H$-$^1H$ COSY spectrum of *ent*-pimara-8(14), 15-diene-19-ol (**11**) in $CDCl_3$, 600 MHz

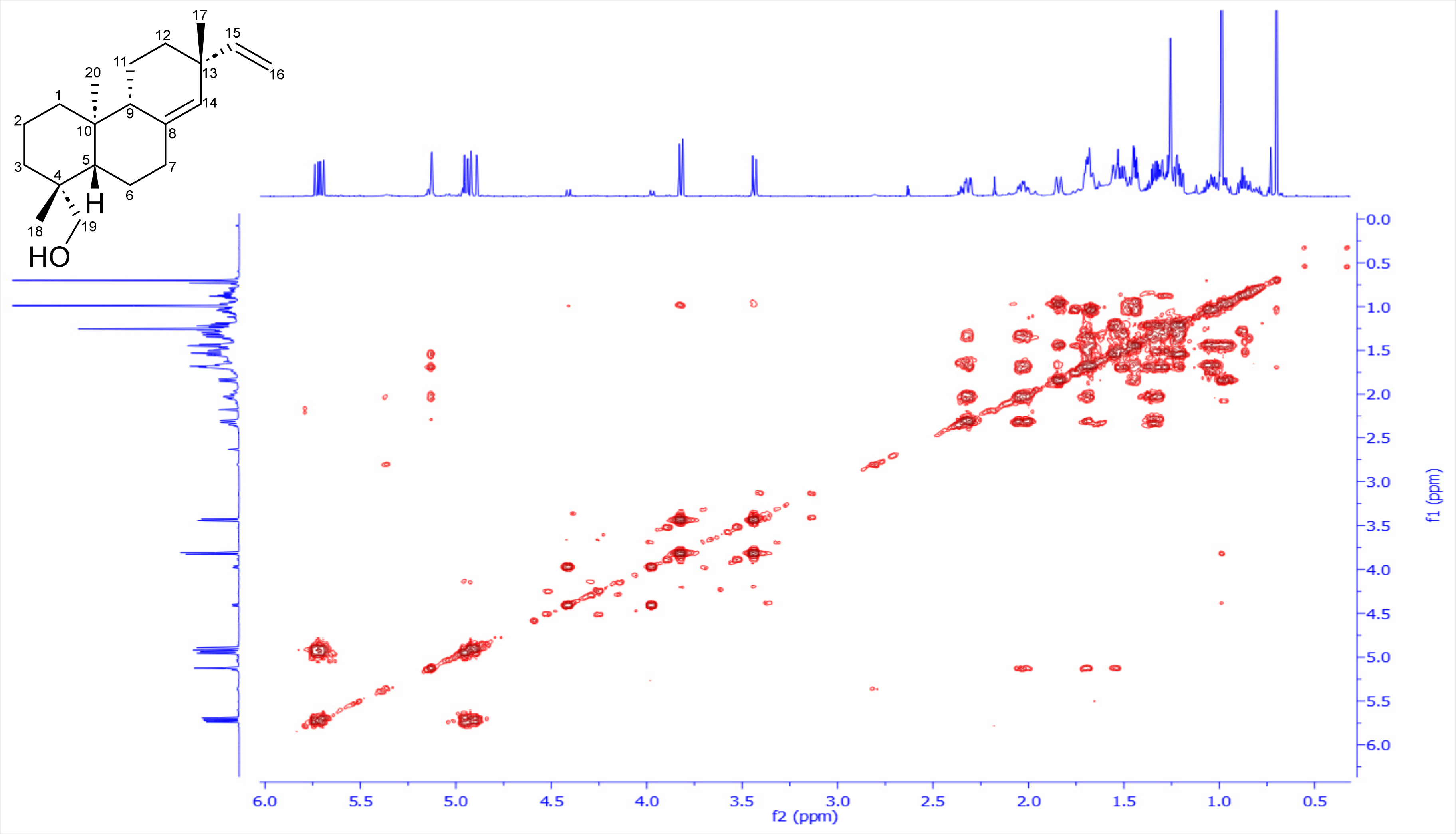

**Figure 17**: [1]H-[13]C HSQC spectrum of *ent*-pimara-8(14), 15-diene-19-ol (**11**) in $CDCl_3$, 600 and 150 MHz

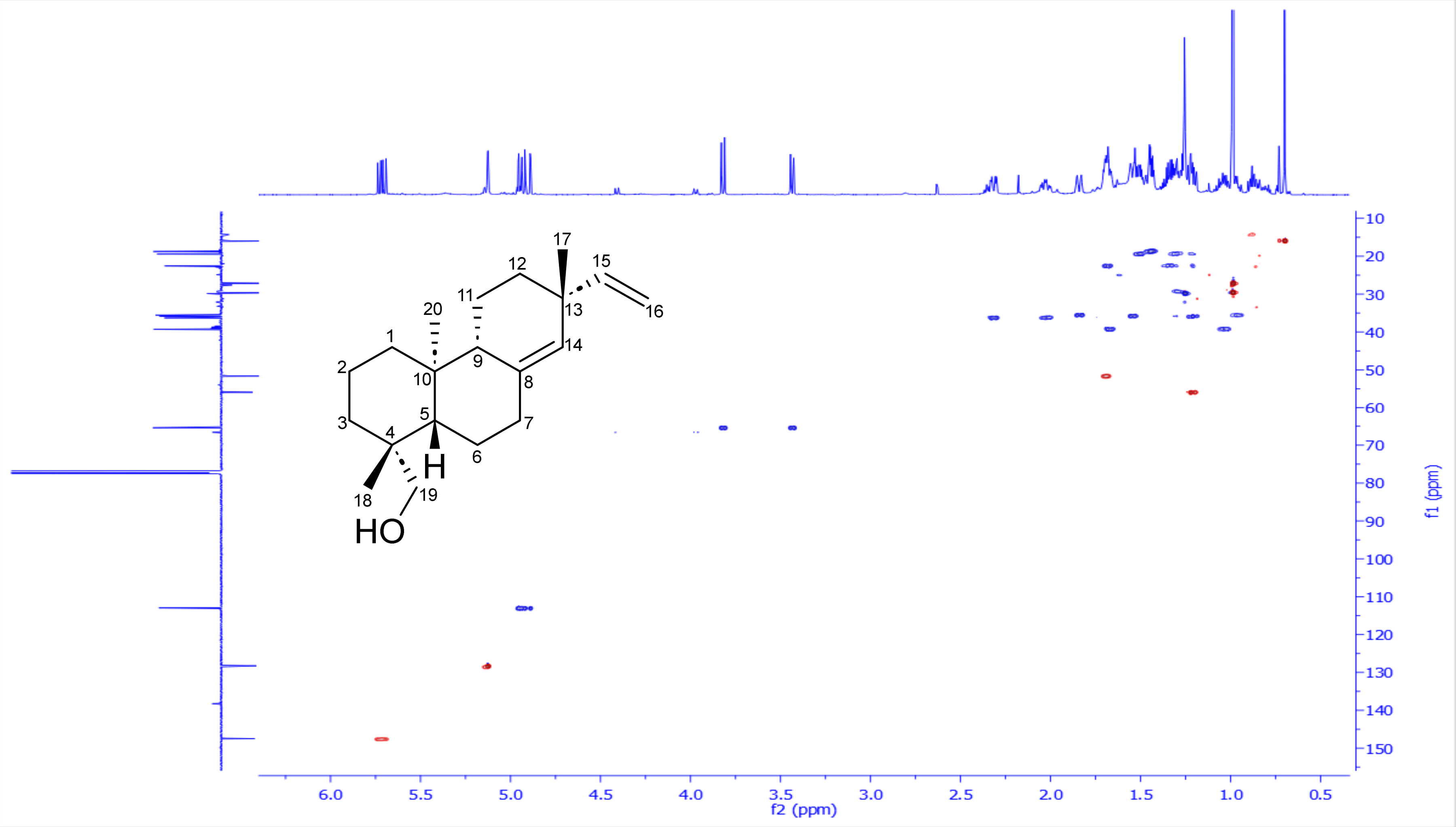

**Figure S18**: $^{1}H$-$^{13}C$ HMBC spectrum of *ent*-pimara-8(14), 15-diene-19-ol (**11**) in $CDCl_3$, 600 and 150 MHz

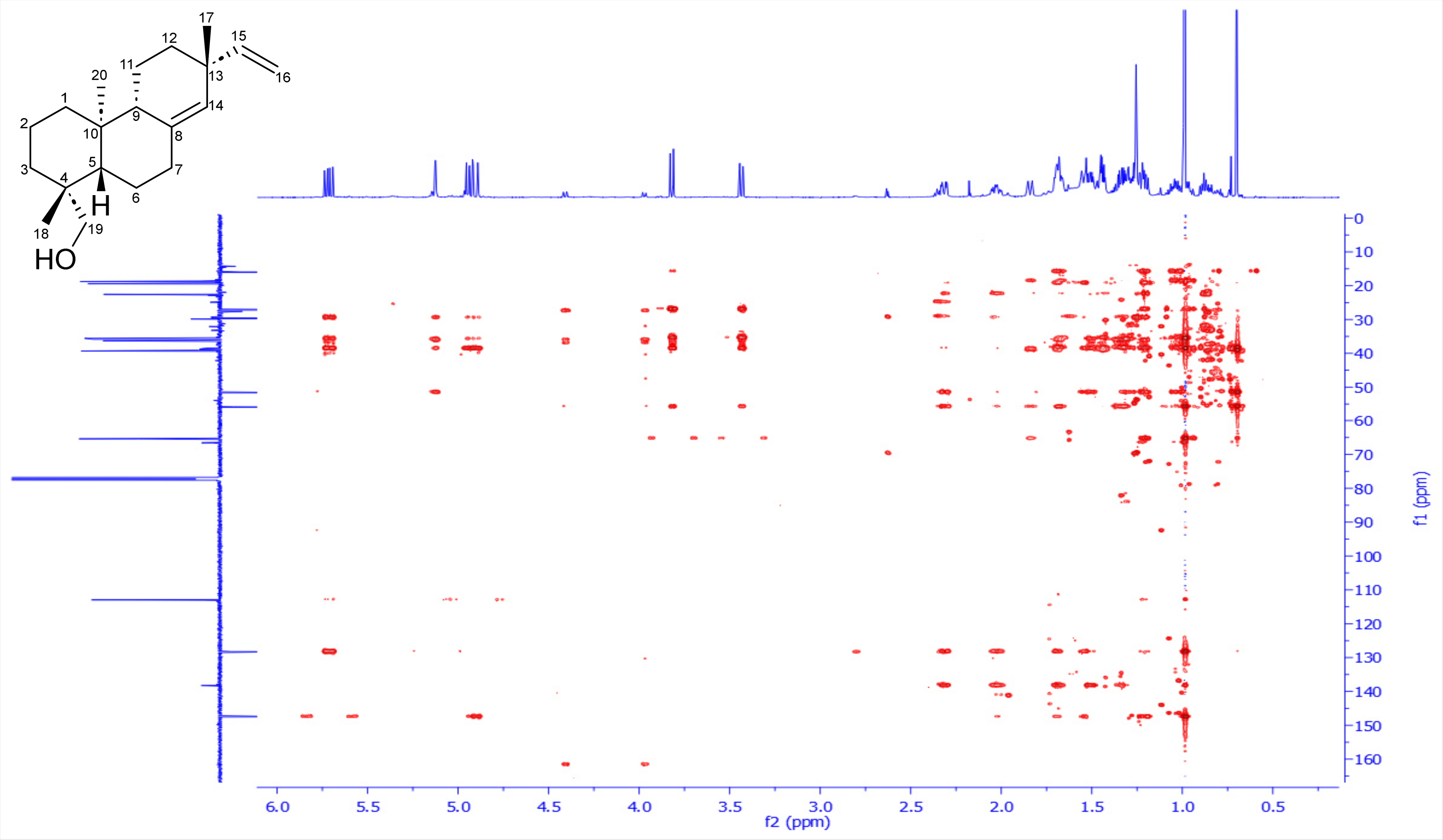

**Figure S19**: $^1$H-$^1$H ROESY spectrum of *ent*-pimara-8(14), 15-diene-19-ol (**11**) in $CDCl_3$, 600 MHz

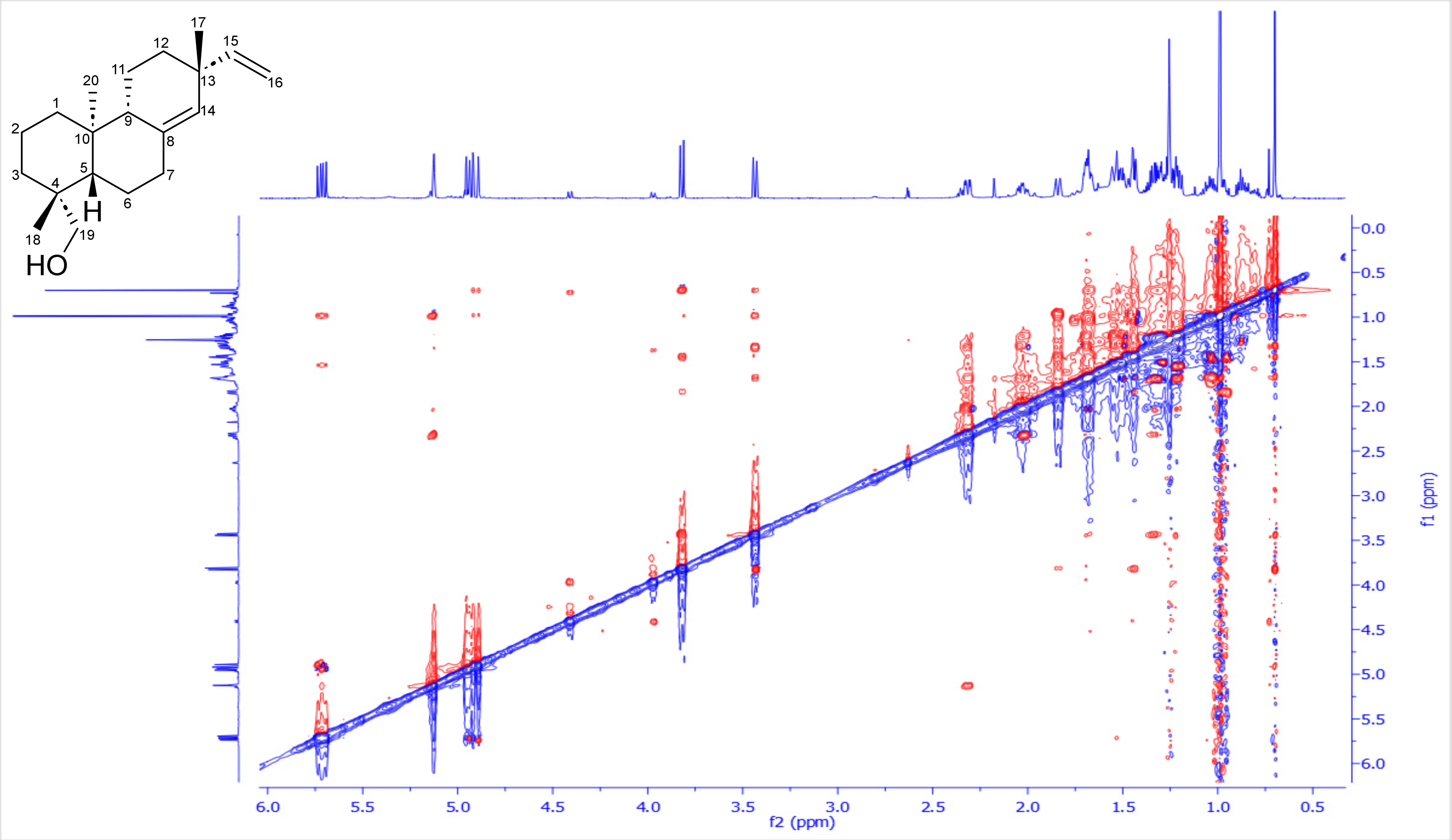

**Figure S20**: DEPTQ-135 spectrum of *ent*-pimara-8(14), 15-diene-19-ol (**11**) in $CDCl_3$, 150 MHz

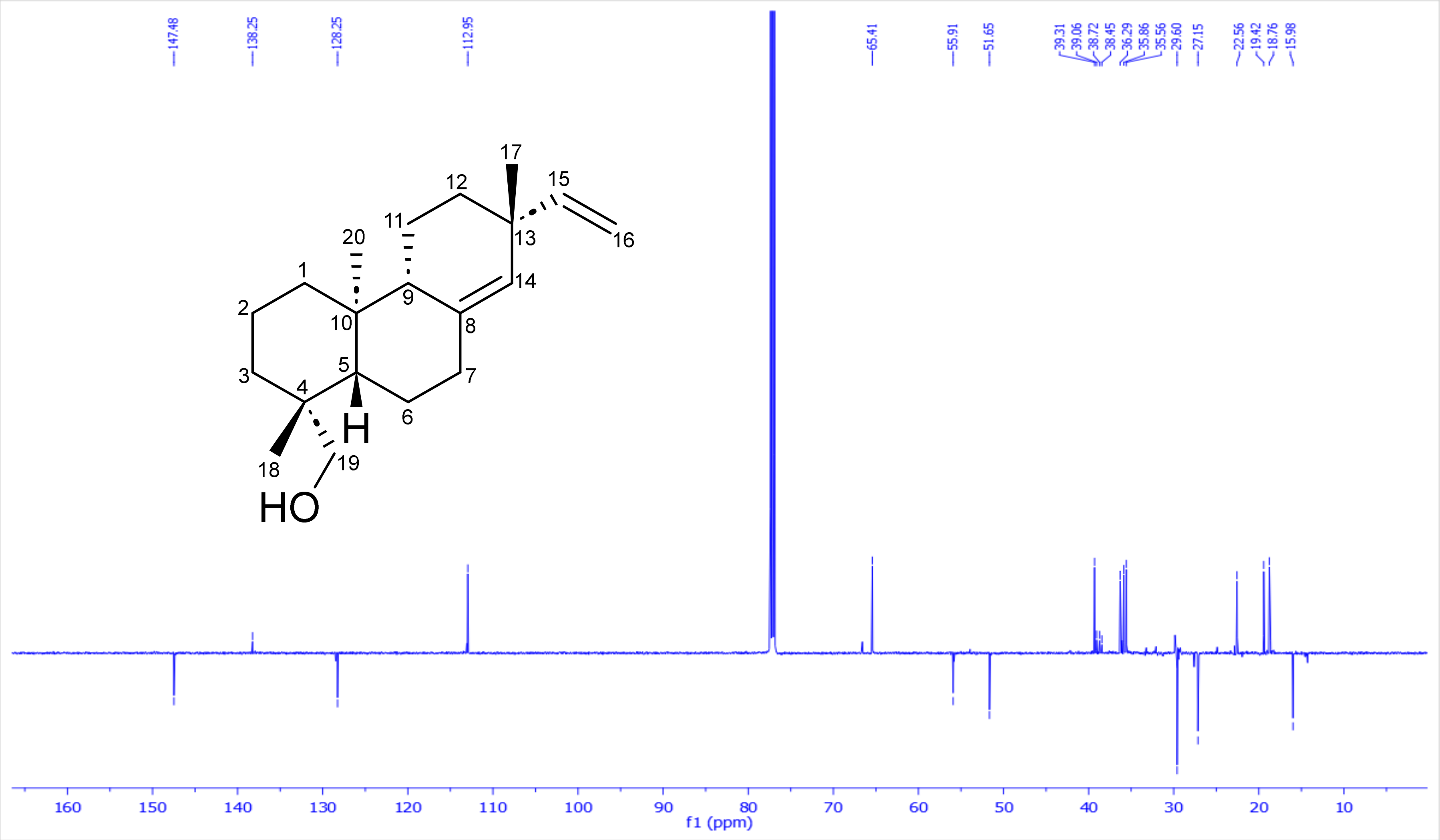

**3. Extraction, purification and structural characterization of *ent*-3-$\alpha$-hydroxylabd-8(17),12Z,14-triene/[(Z)-biformene-3$\alpha$-ol] (13)**

*TaCPS_B1, TaKSL_B3* and *TaCYP99A39_B2* were transiently co-expressed in **75** *Nicotiana benthamiana* plants to afford **80 g** of dry leaves. Subsequently, it was exhaustively extracted using ethyl acetate via speed extractor. The organic extract was collected and dried under reduced pressure to afford an intense green viscus material. Afterwards, it was introduced to three repetitive normal phase flash chromatography using a linear gradient of hexane/ethyl acetate [100/0 up 50/50] along 30 minutes, affording 12 subfractions F1-F12, where all have been monitored by GC-MS. Promising fractions F5-F6 were combined and further purified using repetitive $C_{18}$ semi-preparative HPLC using a short linear gradient of water/acetonitrile [15/85 up 0/100], 4 mL/min, acidified with 0.1 % FA along 27 min to finally afford *ent*-3-$\alpha$--hydroxylabda-8(17),12(Z)-14-triene/[(Z)-biformene-3$\alpha$-ol] (**13**) (**95** mg, pale yellow oily material). Its full structure was resolved based on extensive 1D & 2D-NMR data analysis.

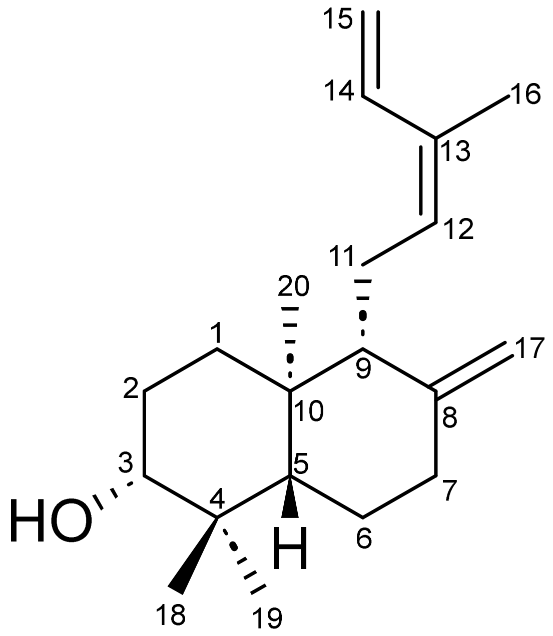

*ent*-3-$\alpha$-hydroxylabd-8(17),12Z,14-triene/[(Z)-biformene-3$\alpha$-ol] (13)

**Table 4**: $^1$H, $^{13}$C NMR spectroscopic data for *ent*-3-$\alpha$-hydroxylabda-8(17),12(Z)-14-triene/[(Z)-biformene-3$\alpha$-ol] (**13**) in $CDCl_3$, 600 and 150 MHz.

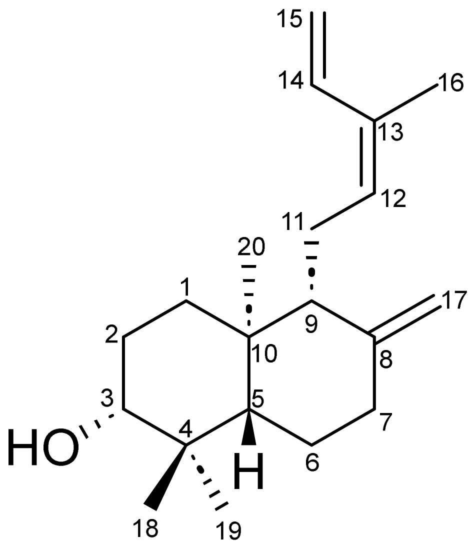

| **Position** | $\delta_C$**, Type** | $\delta_H$**, mult. (*J* in Hz)** |
|---|---|---|
| **1** | 37.3, $CH_2$ | 1.82, *dt* (13.1, 3.6) |
| | | 1.23, *dd* (13.4, 3.8) |
| **2** | 28.0, $CH_2$ | 1.71/1.60, *m* |
| **3** | 78.9, CH | 3.26, *dd* (11.8, 4.4) |
| **4** | 39.3, Cq | - |
| **5** | 54.7, CH | 1.11, *dd* (12.5, 2.7) |
| **6** | 23.9, $CH_2$ | 1.73, *m*/1.40, *dd* (12.9, 4.3) |
| **7** | 38.1, $CH_2$ | 2.41, *dd* (4.2, 2.4) |
| | | 2.00, *td* (13.1, 5.1) |
| **8** | 148.1, Cq | - |
| **9** | 57.2, CH | 1.68, *m* |
| **10** | 39.5, Cq | - |
| **11** | 22.3, $CH_2$ | 2.39, *dd* (4.1, 2.5)/2.18, *m* |
| **12** | 131.6, CH | 5.30, *t* (6.7) |
| **13** | 131.9, Cq | - |
| **14** | 134.0, CH | 6.78, *dd* (17.3, 10.8) |
| **15** | 113.5, $CH_2$ | 5.18, *d* (16.9) |
| | | 5.09, *d* (10.8) |
| **16** | 19.9, $CH_3$ | 1.77, br *d* (1.0) |
| **17** | 108.2, $CH_2$ | 4.84, br *d* (1.3) |
| | | 4.49, br *d* (1.0) |
| **18** | 28.5, $CH_3$ | 1.00, *s* |
| **19** | 15.6, $CH_3$ | 0.78, *s* |
| **20** | 14.6, $CH_3$ | 0.73, *s* |

**Scheme 7**: Key [1]H-[1]H COSY (**bold blue**), [1]H-[13]C HMBC (H→C, red arrows) observed for *ent*-3-*α*-hydroxylabda-8(17),12(Z)-14-triene/[(Z)-biformene-3α-ol] (**13**)

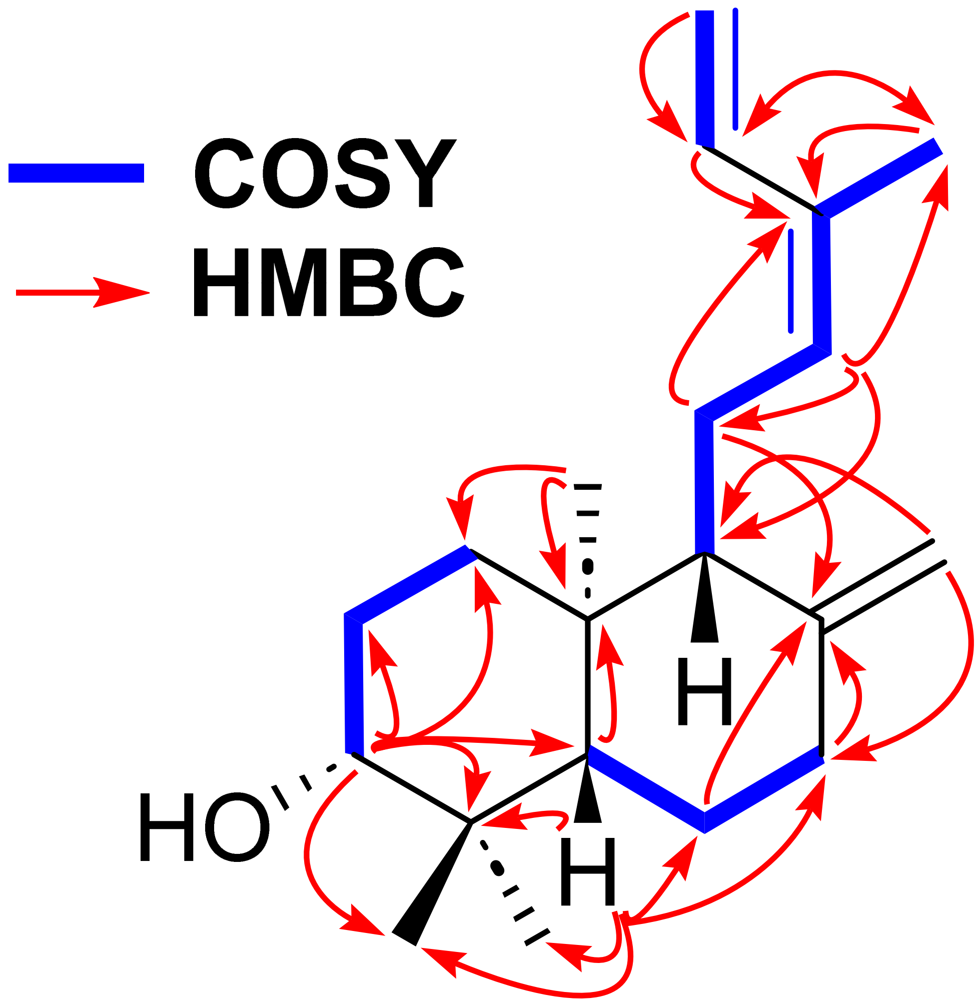

**Scheme 8**: Key ¹H-¹H ROESY (H↔H, green) observed for *ent*-3-α-hydroxylabda-8(17),12(Z)-14-triene/[(Z)-biformene-3α-ol] (**13**)

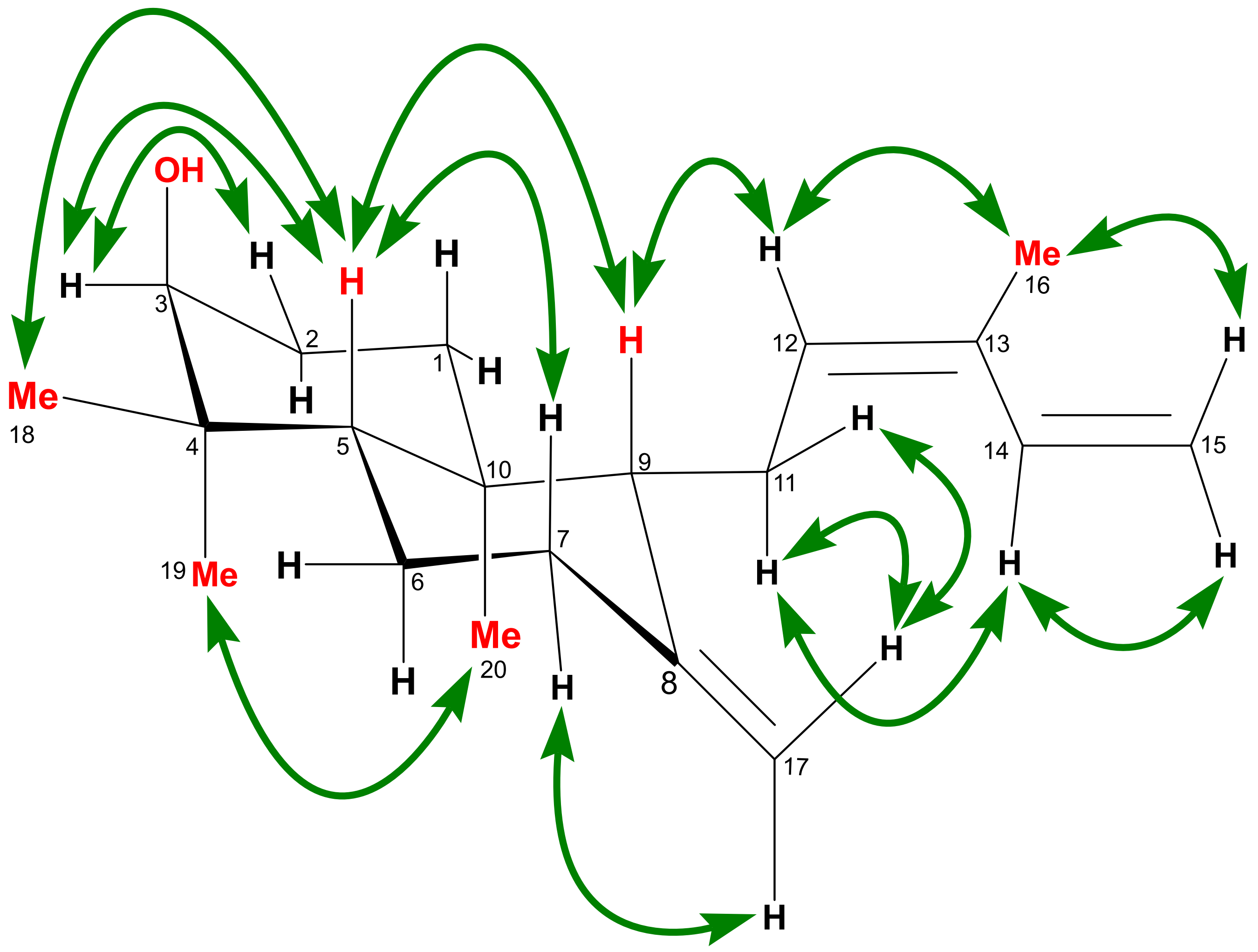

**Figure S21**: [1]H-NMR spectrum of *ent*-3-$\alpha$-hydroxylabda-8(17),12(Z)-14-triene/[(Z)-biformene-3$\alpha$-ol] (**13**) in $CDCl_3$, 600 MHz

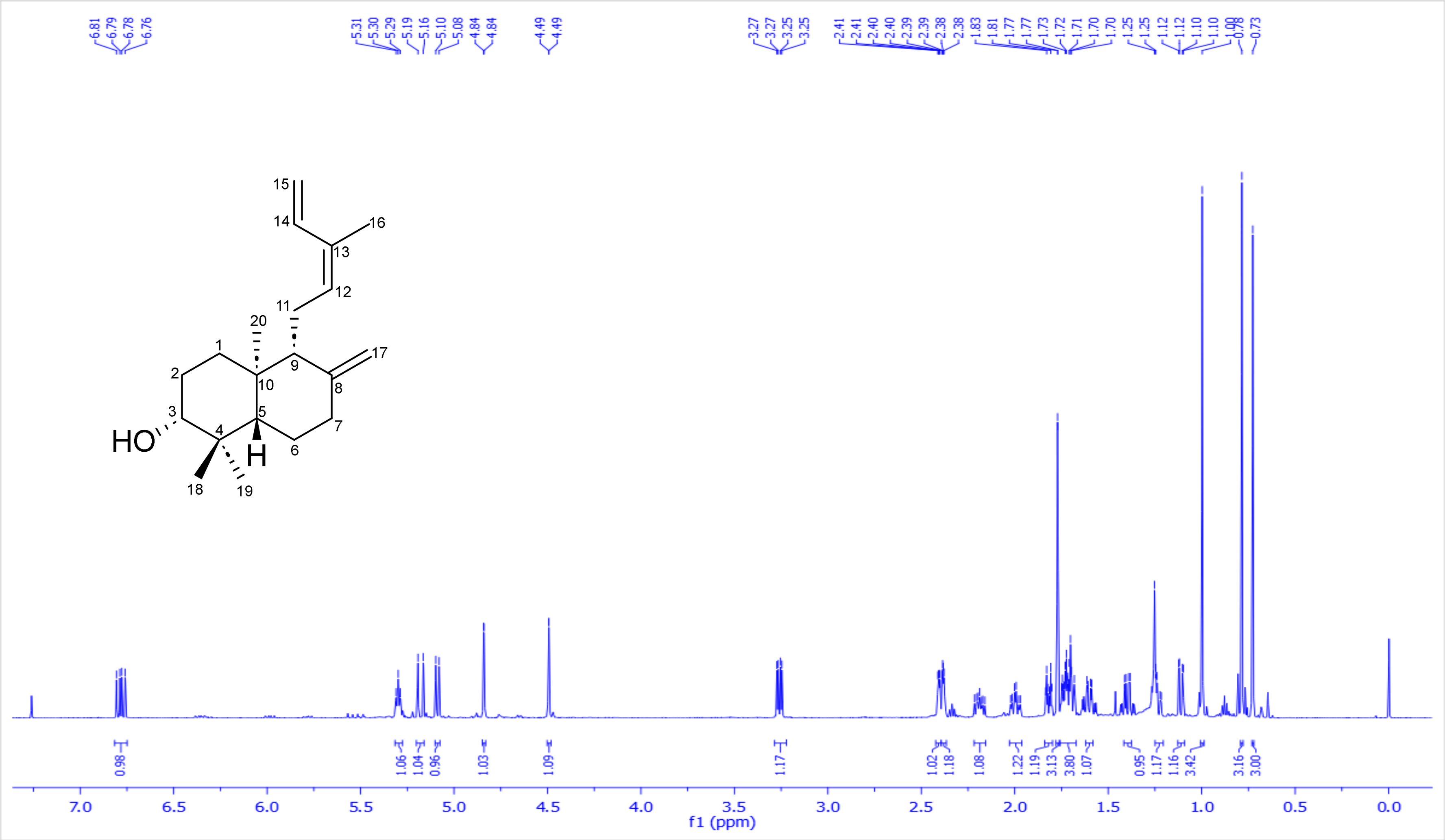

**Figure S22**: ¹H-¹H COSY spectrum of *ent*-3-$\alpha$-hydroxylabda-8(17),12(Z)-14-triene/[(Z)-biformene-3$\alpha$-ol] (**13**) in $CDCl_3$, 600 MHz

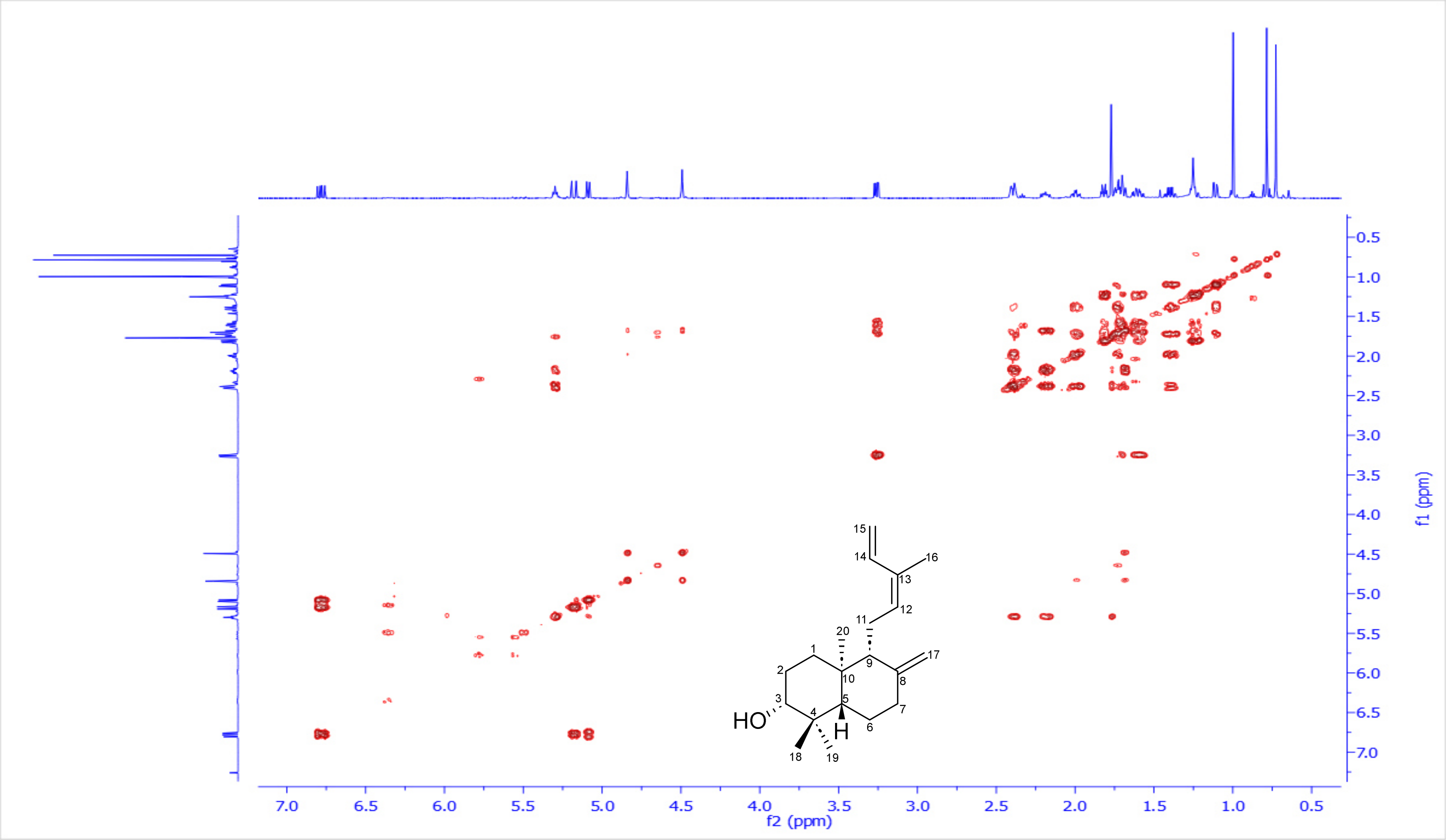

**Figure S23**: $^{1}$H-$^{13}$C HSQC spectrum of *ent*-3-$\alpha$-hydroxylabda-8(17),12(Z)-14-triene/[(Z)-biformene-3$\alpha$-ol] (**13**) in $CDCl_3$, 600 and 150 MHz

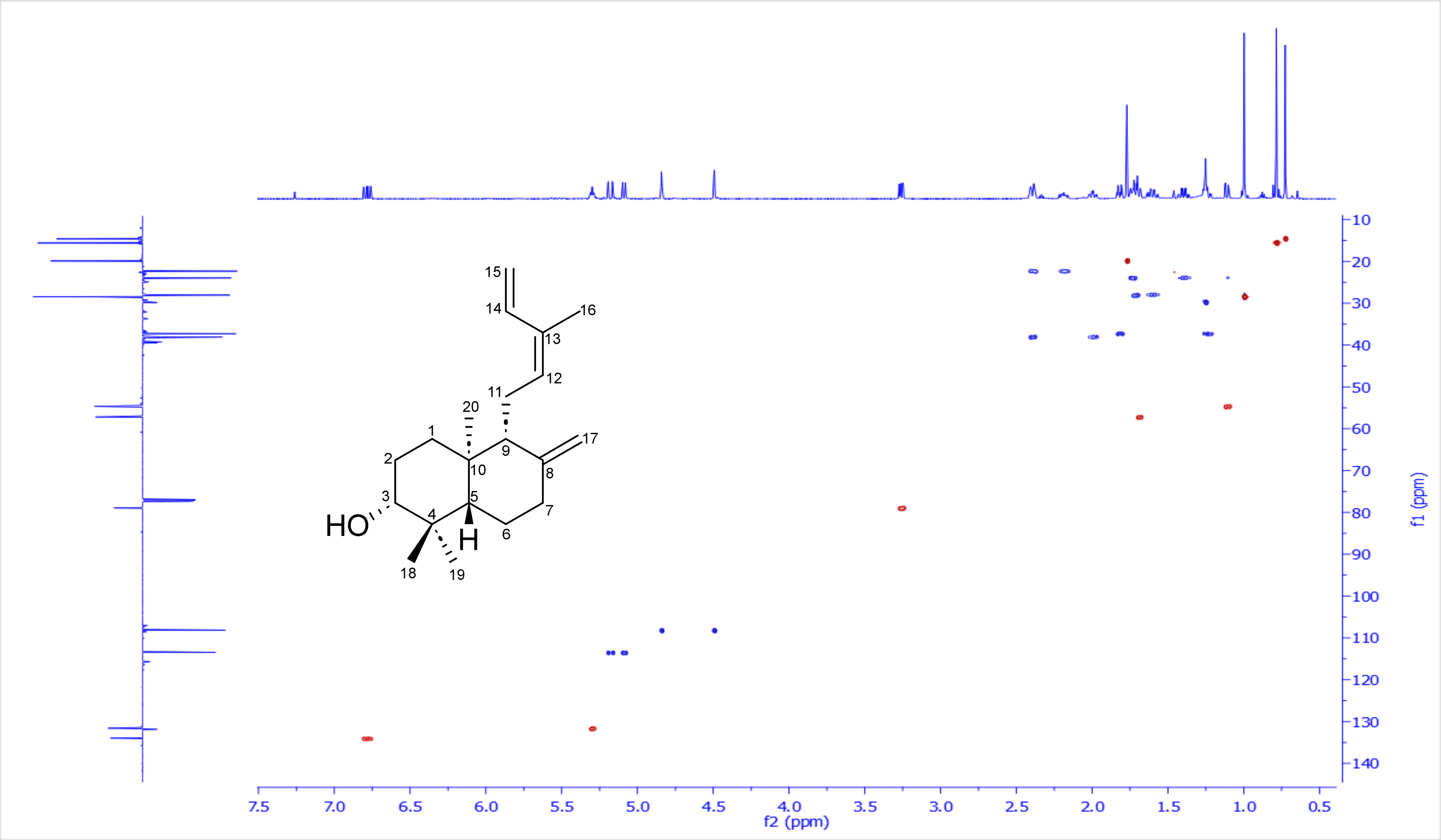

**Figure S24**: $^{1}H$-$^{13}C$ HSQC-TOCSY spectrum of *ent*-3-*α*-hydroxylabda-8(17),12(Z)-14-triene/[(Z)-biformene-3α-ol] (**13**) in $CDCl_3$, 600 and 150 MHz

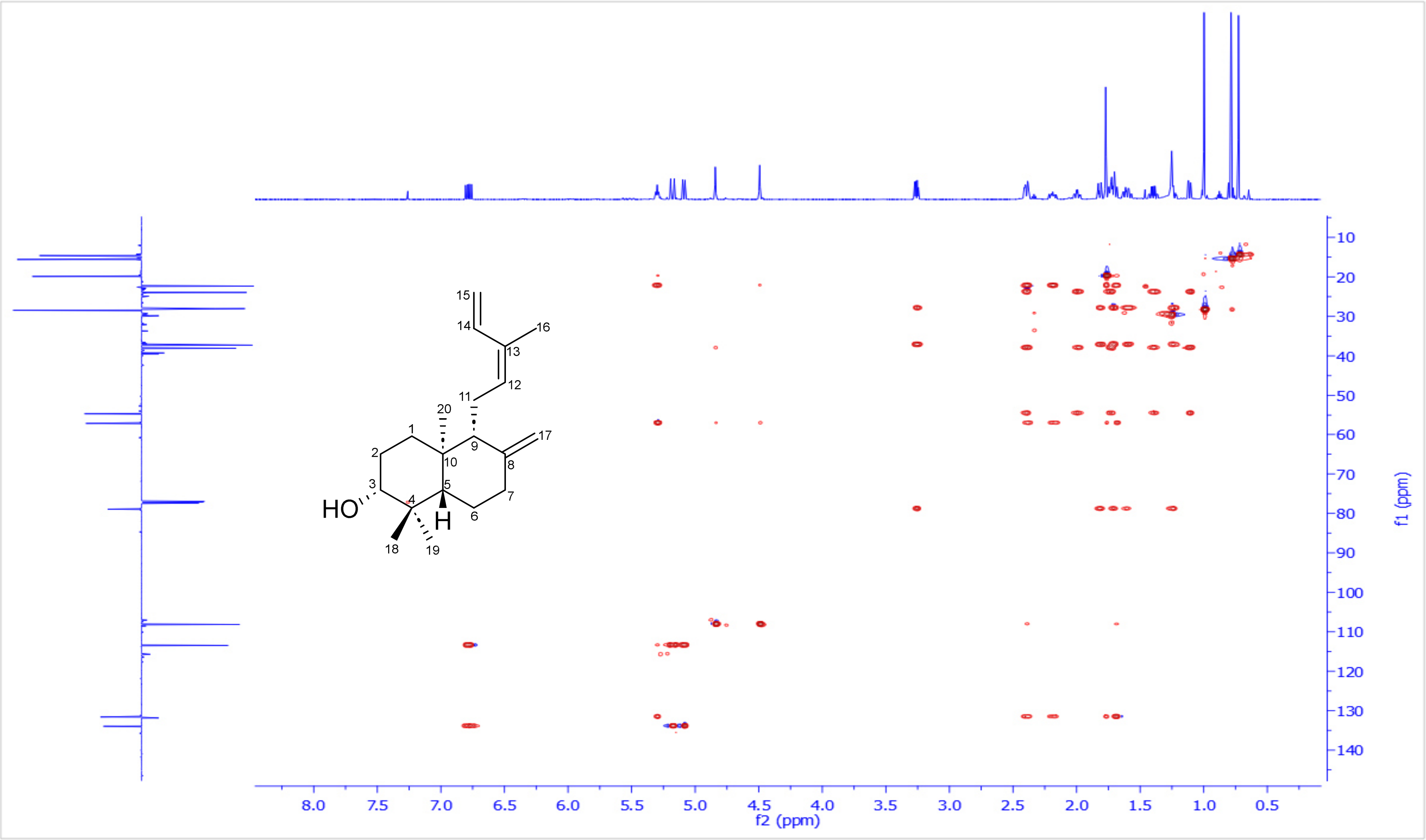

**Figure S25**: $^{1}H$-$^{13}C$ HMBC spectrum of *ent*-3-α-hydroxylabda-8(17),12(Z)-14-triene/[(Z)-biformene-3α-ol] (**13**) in $CDCl_3$, 600 and 150 MHz

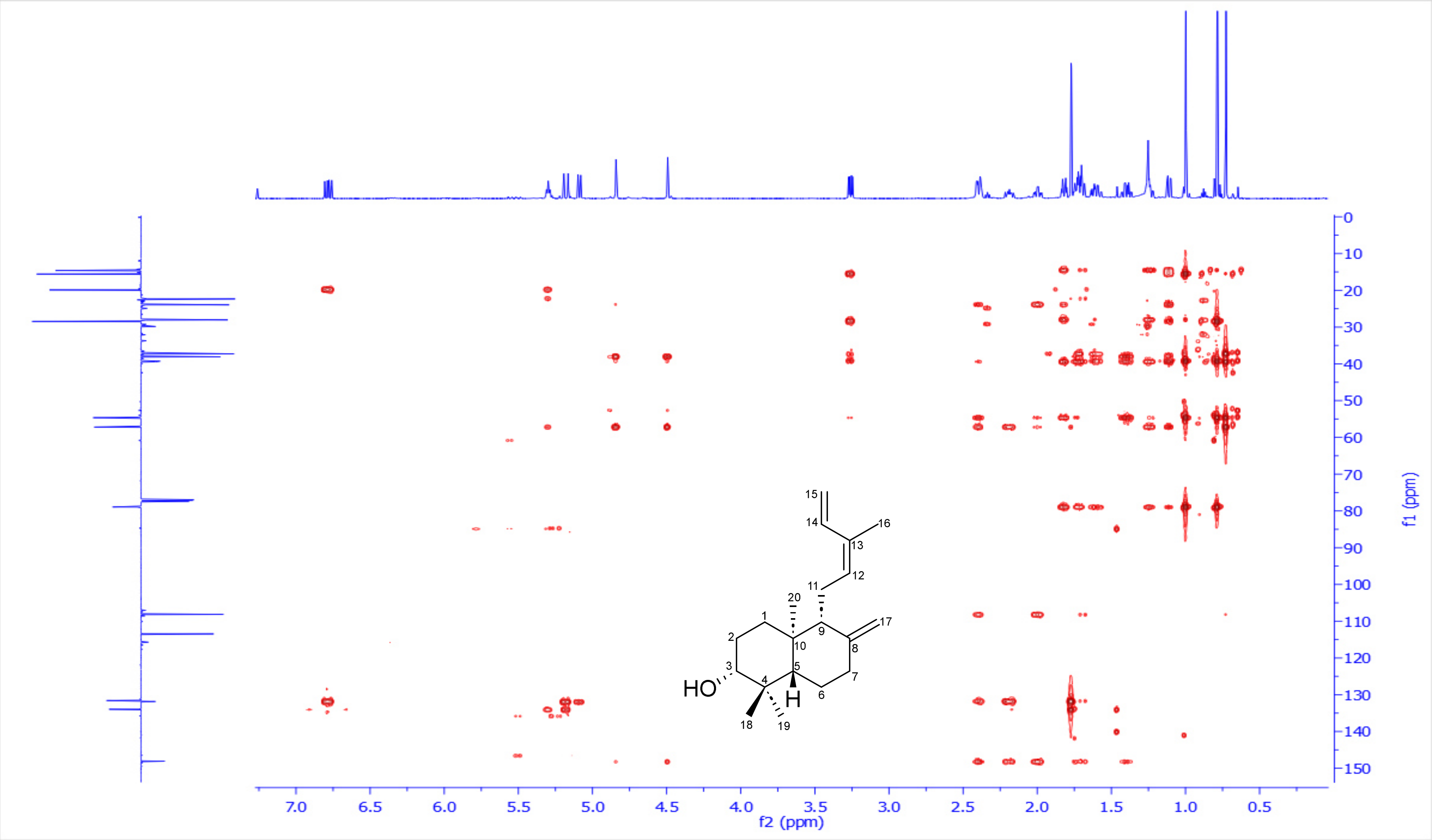

**Figure S26**: $^{1}$H-$^{1}$H ROESY spectrum of *ent*-3-$\alpha$-hydroxylabda-8(17),12(Z)-14-triene/[(Z)-biformene-3$\alpha$-ol] (**13**) in $CDCl_3$, 600 MHz

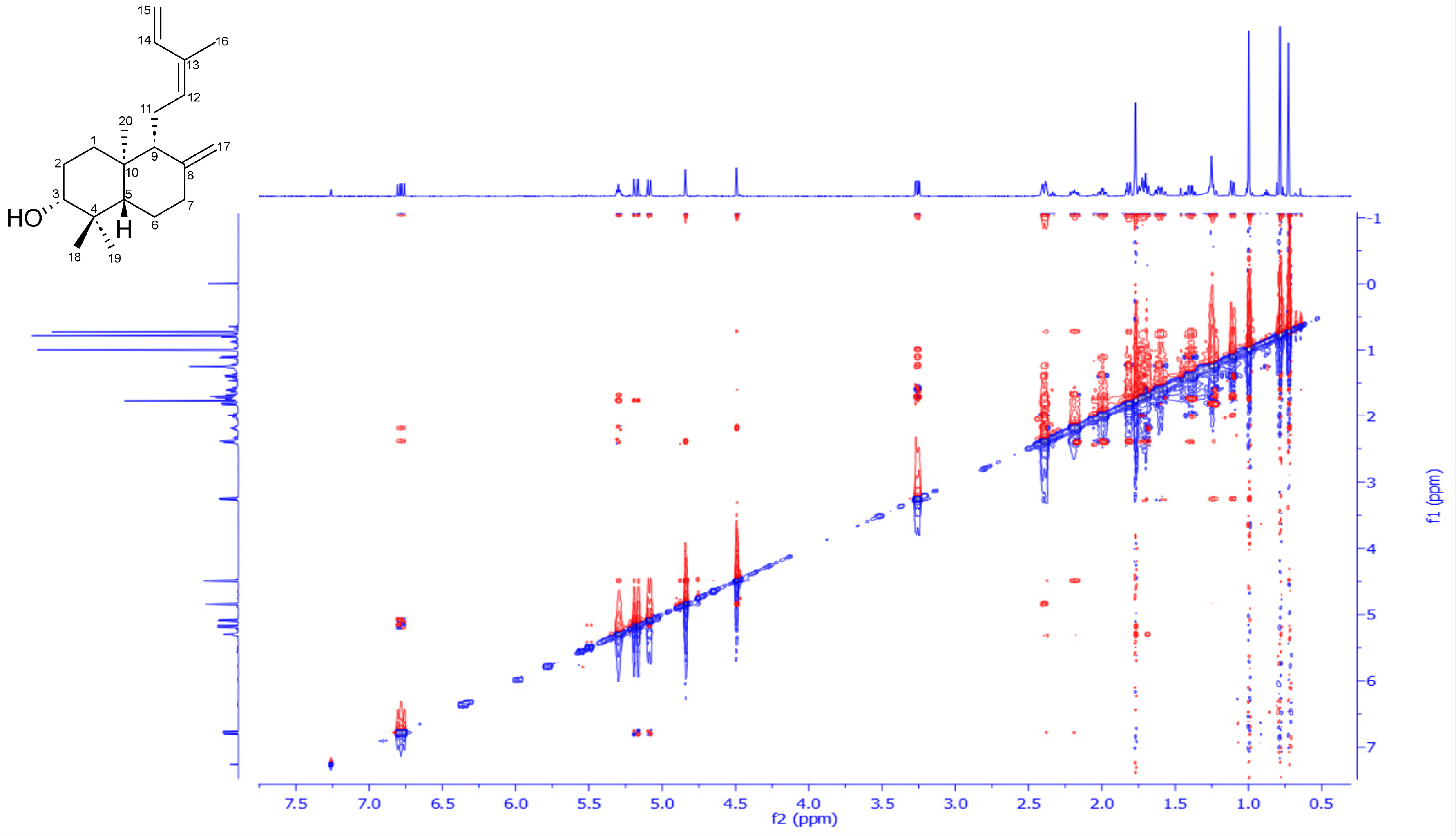

**Figure S27**: DEPTQ-135 spectrum of *ent*-3-$\alpha$-hydroxylabda-8(17),12(Z)-14-triene/[(Z)-biformene-3$\alpha$-ol] (**13**) in $CDCl_3$, 150 MHz

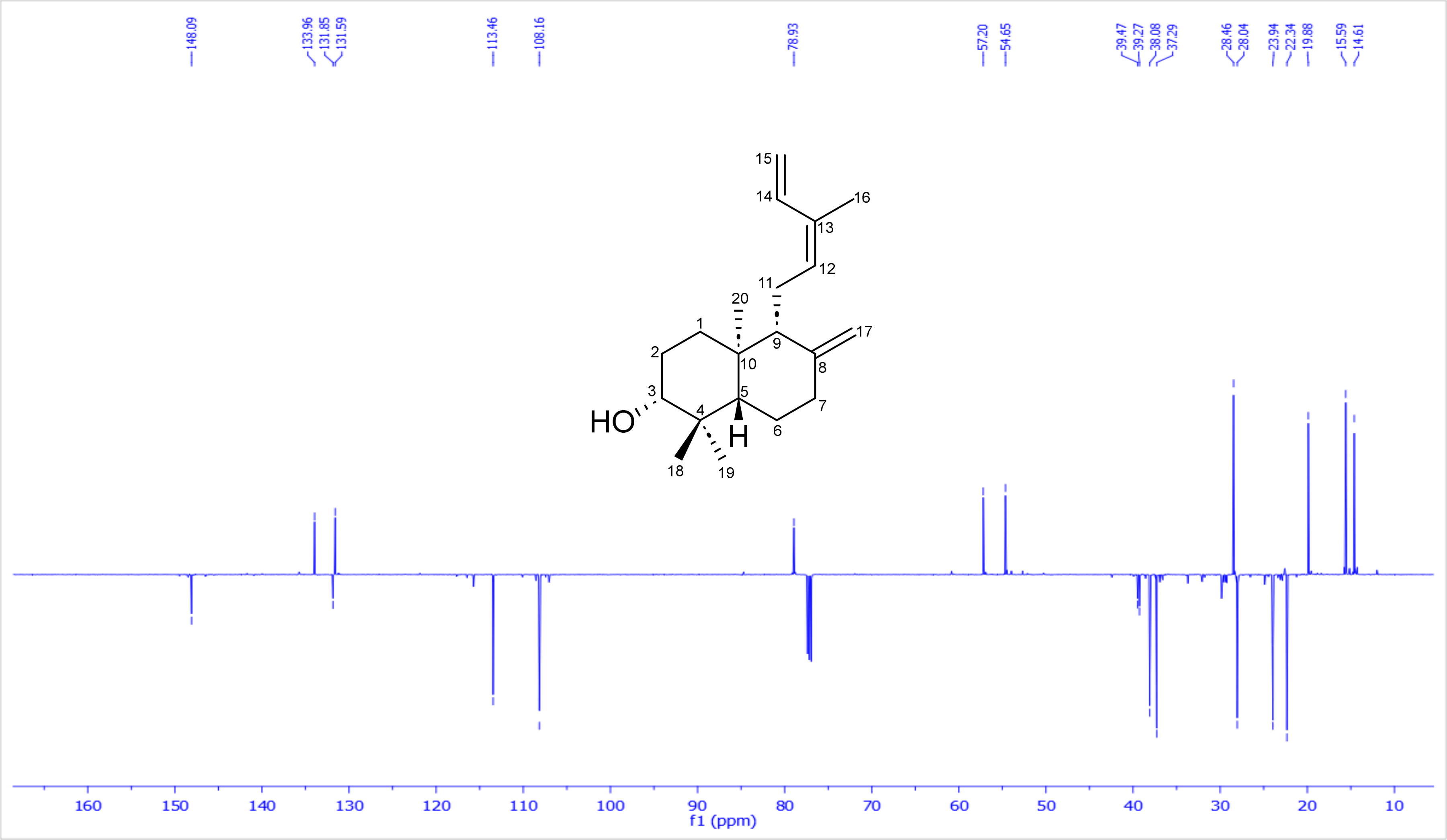

### 4. Extraction, purification and structural characterization of *ent*-15, 19-dihydroxy-labda-8(17),13(*E*)-diene (14)

*TaCPS_B1 and TaCYP701A64_B2* were transiently co-expressed in **75** *Nicotiana benthamiana* plants to afford **80 g** of dry leaves. Subsequently, it was exhaustively extracted using ethyl acetate via speed extractor. The organic extract was collected and dried under reduced pressure to afford an intense green viscus material. Afterwards, it was introduced to three repetitive normal phase flash chromatography using a linear gradient of hexane/ethyl acetate [100/0 up 0/100] along 30 minutes, affording 12 subfractions F1-F12, where all have been monitored by LC-MS. Promising fractions F6 were further purified using repetitive $C_{18}$ semi-preparative HPLC using a short gradient of water/acetonitrile [15/85 up 0/100], 4 mL/min, acidified with 0.1 % FA along 27 min to finally afford *ent*-15,19-dihydroxy-labda-8(17),13(*E*)-diene **(14)** (**0.5** mg, pale yellow powder). Its full structure was resolved based on extensive 1D & 2D-NMR data analysis.

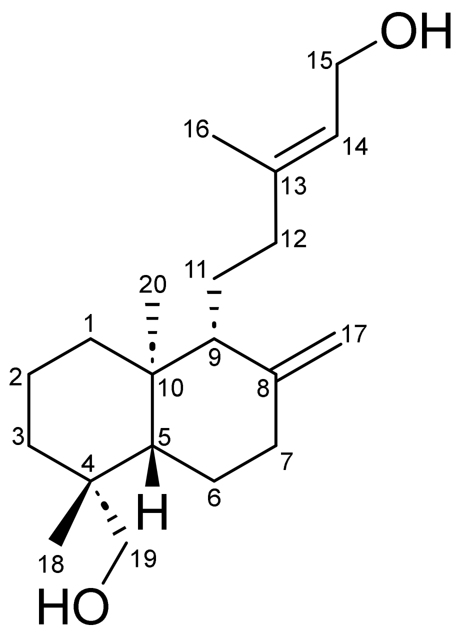

***ent*-15, 19-dihydroxy-labda-8(17),13(*E*)-diene (14)**

**Table 5**: $^1$H, $^{13}$C NMR spectroscopic data for *ent*-15,19-dihydroxy-labda-8(17),13(*E*)-diene **(14)** in $CDCl_3$, 600 and 150 MHz.

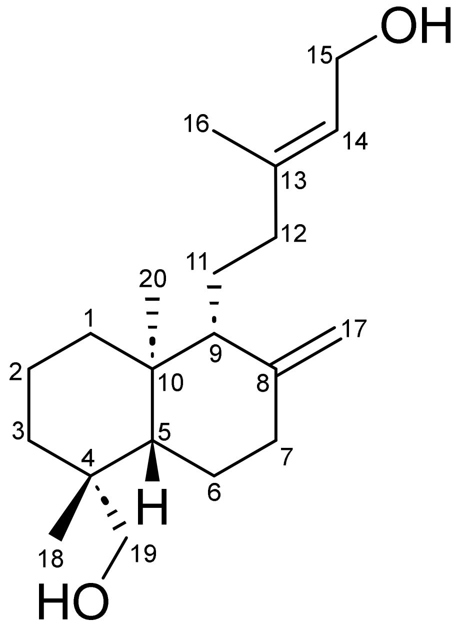

| Position | $\delta_C$, Type | $\delta_H$, mult. (*J* in Hz) |
|---|---|---|
| **1** | 39.2, $CH_2$ | 1.79, *m* |
| | | 1.05, *td* (12.7, 4.9) |
| **2** | 19.2, $CH_2$ | 1.51/1.51, *m* |
| **3** | 35.6, $CH_2$ | 1.82/0.97, *m* |
| **4** | 38.7, Cq | - |
| **5** | 56.5, CH | 1.25, *m* |
| **6** | 24.6, $CH_2$ | 1.81/1.32, *m* |
| **7** | 38.8, $CH_2$ | 2.49, *ddd* (12.6, 3.8, 2.5) |
| | | 1.93, *td* (12.5, 5.2) |
| **8** | 148.2, Cq | - |
| **9** | 56.5, CH | 1.59, *m* |
| **10** | 39.7, Cq | - |
| **11** | 22.0, $CH_2$ | 1.60/1.42, *m* |
| **12** | 38.5, $CH_2$ | 2.15/1.81, *m* |
| **13** | 140.7, Cq | - |
| **14** | 123.2, CH | 5.39, *td* (6.9, 1.0) |
| **15** | 59.6, $CH_2$ | 4.15, *d* (6.5) |
| **16** | 16.5, $CH_3$ | 1.68, *br s* |
| **17** | 106.7, $CH_2$ | 4.82/4.52, *br s* |
| **18** | 27.2, $CH_3$ | 0.98, *s* |
| **19** | 65.3, $CH_2$ | 3.75, *d* (10.7) |
| | | 3.40, *d* (10.9) |
| **20** | 15.5, $CH_3$ | 0.66, *s* |

**Scheme 9**: Key ¹H-¹H COSY (**bold blue**), ¹H-¹³C HMBC (H→C, red arrows) observed for *ent*-15,19-dihydroxy-labda-8(17),13(*E*)-diene **(14)**

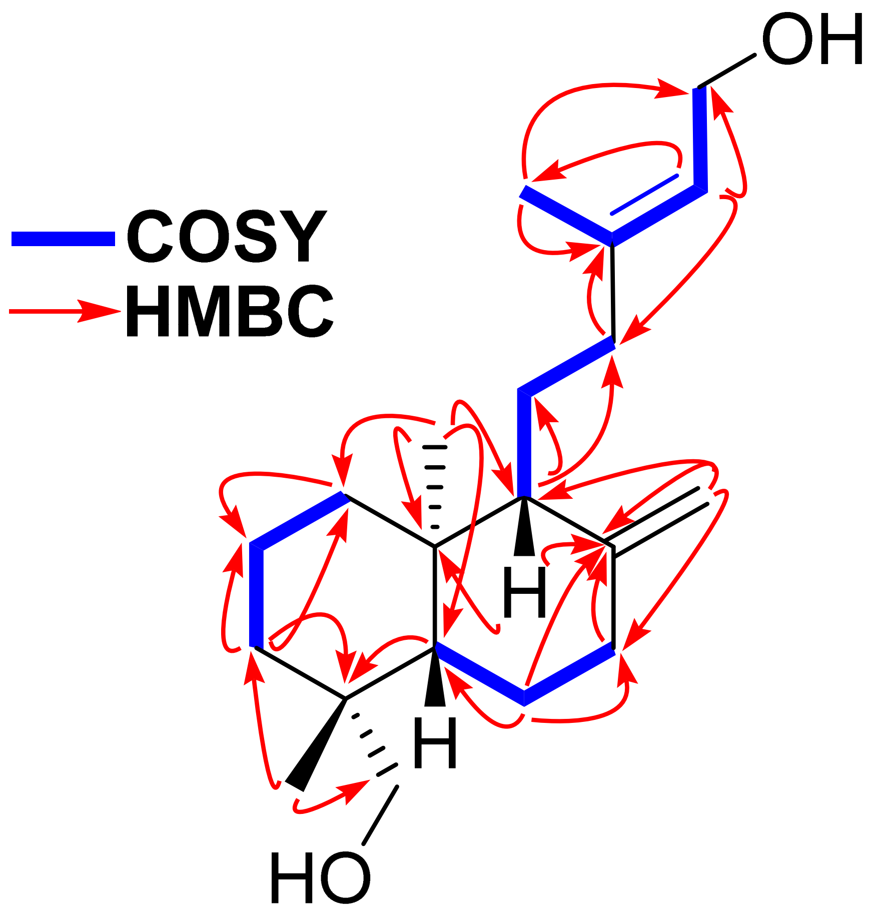

**Scheme 10**: Key ¹H-¹H ROESY (H↔H, green) observed for *ent*-15,19-dihydroxy-labda-8(17),13(*E*)-diene **(14)**

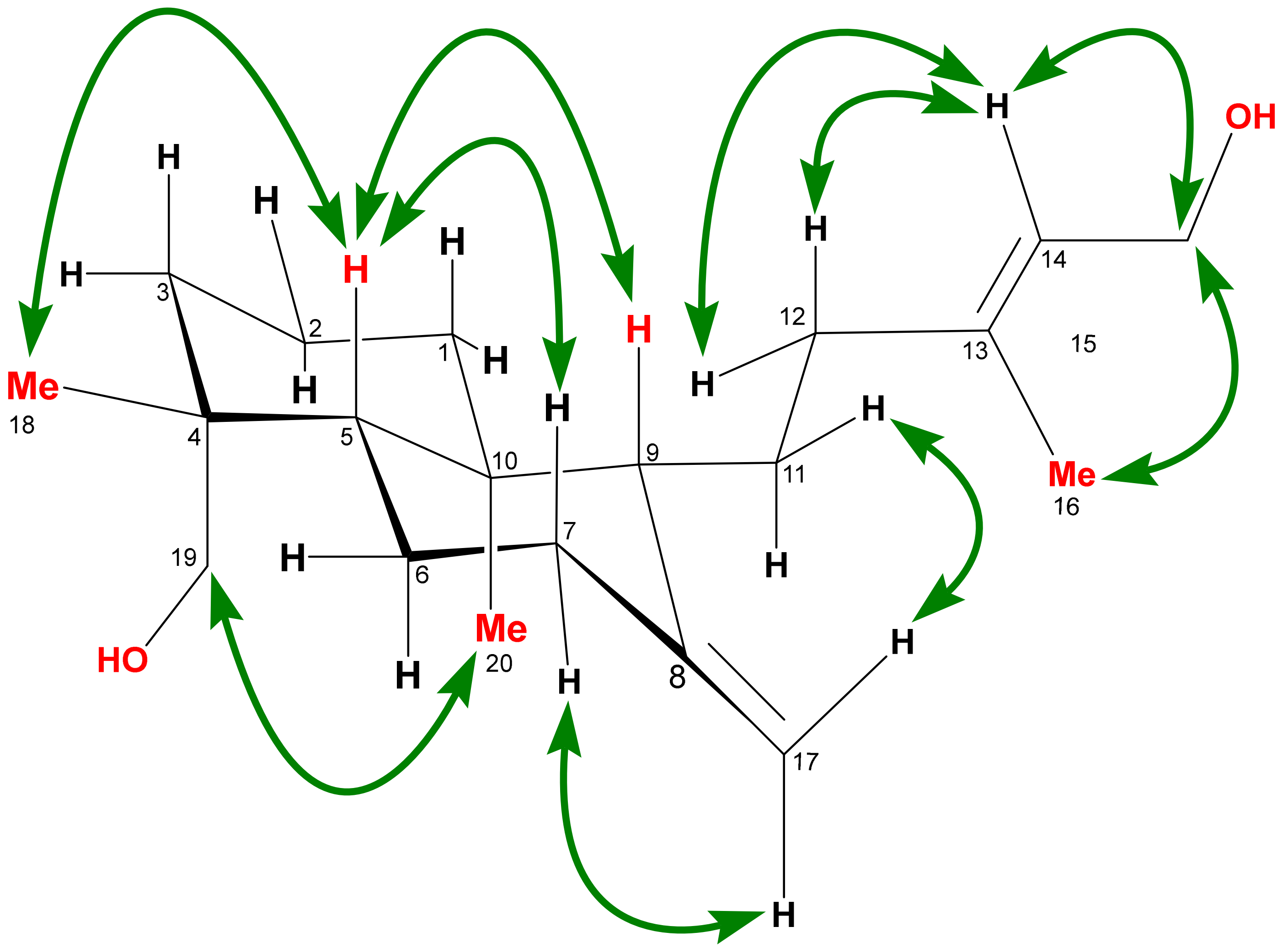

**Figure S28**: $^{1}$H-NMR spectrum of *ent*-15,19-dihydroxy-labda-8(17),13(*E*)-diene **(14)** in $CDCl_3$, 600 MHz

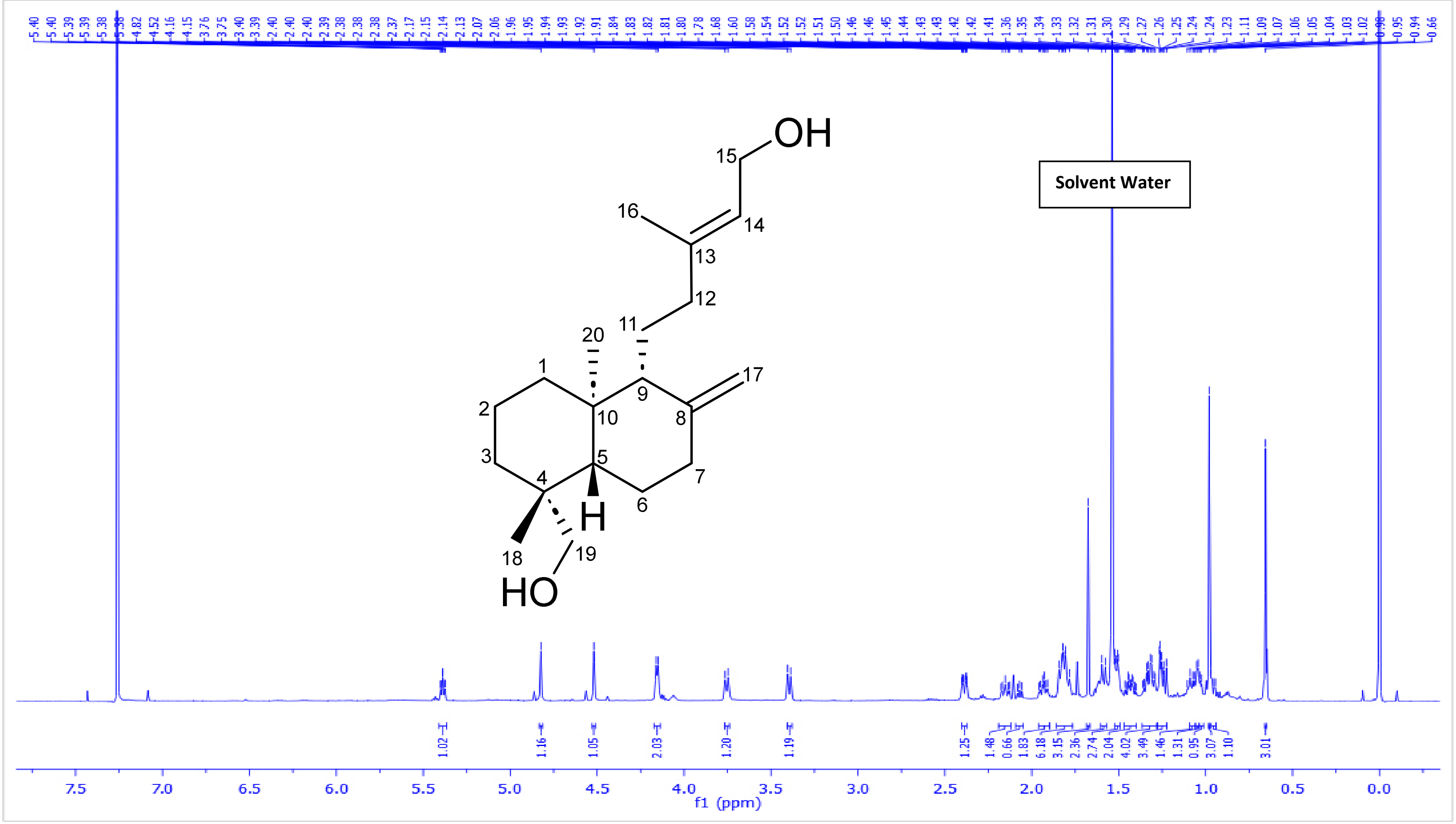

**Figure S29**: $^1H$-$^1H$ COSY spectrum of *ent*-15,19-dihydroxy-labda-8(17),13(*E*)-diene **(14)** in $CDCl_3$, 600 MHz

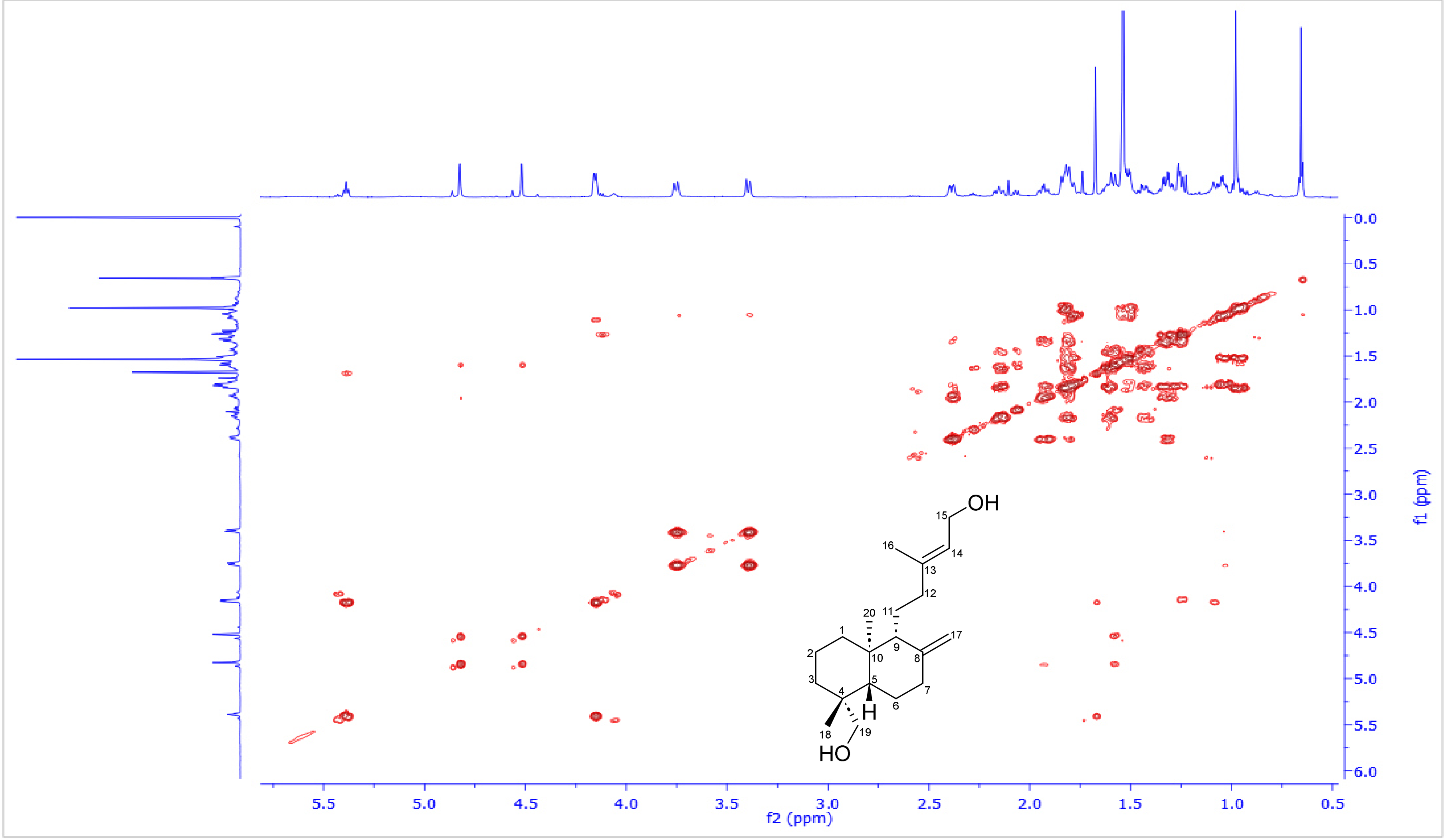

**Figure S30**: $^{1}$H-$^{13}$C HSQC spectrum of *ent*-15,19-dihydroxy-labda-8(17),13(*E*)-diene **(14)** in $CDCl_3$, 600 and 150 MHz

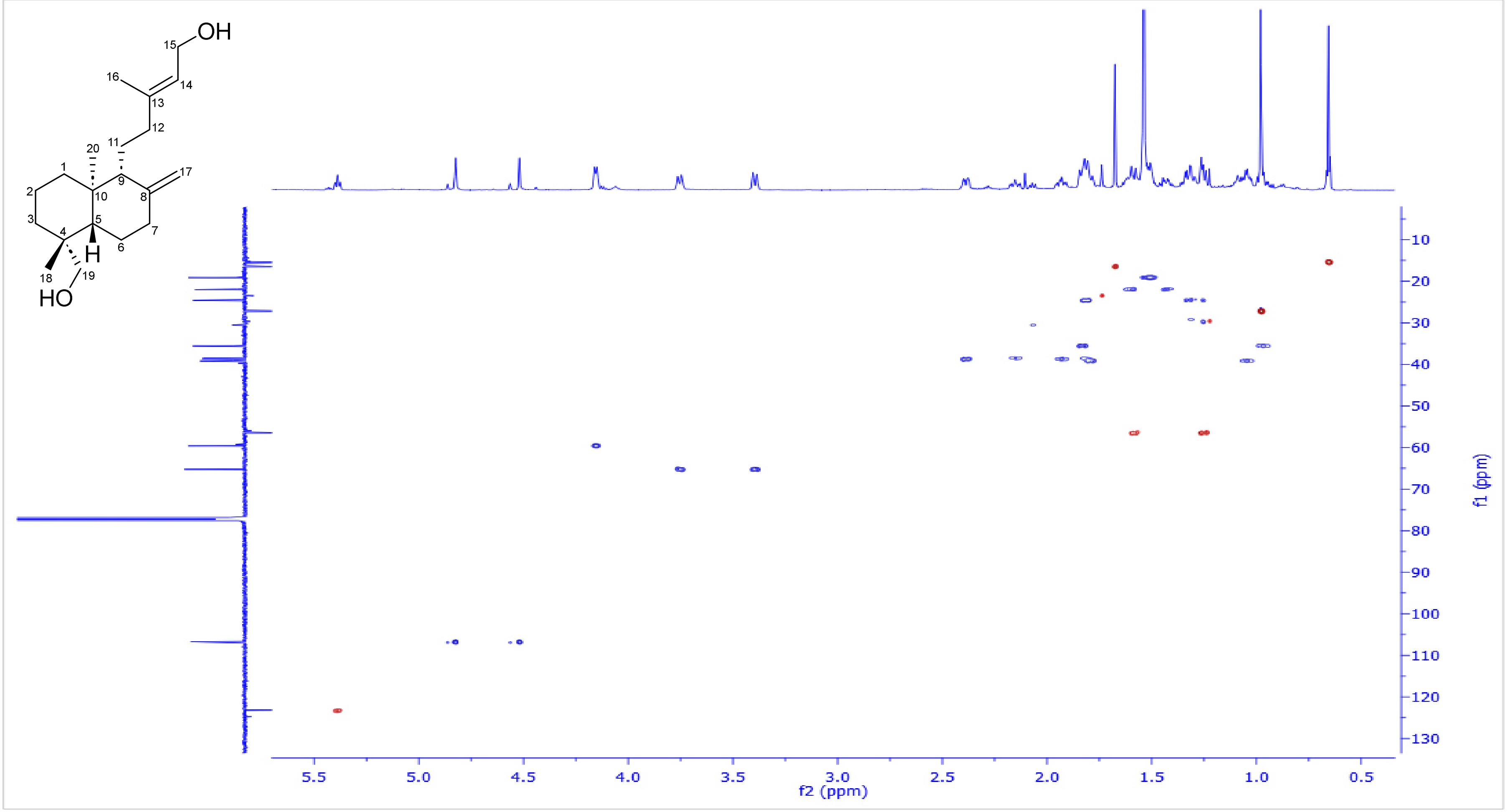

**Figure S31**: $^1$H-$^{13}$C HSQC-TOCSY spectrum of *ent*-15,19-dihydroxy-labda-8(17),13(*E*)-diene **(14)** in $CDCl_3$, 600 and 150 MHz

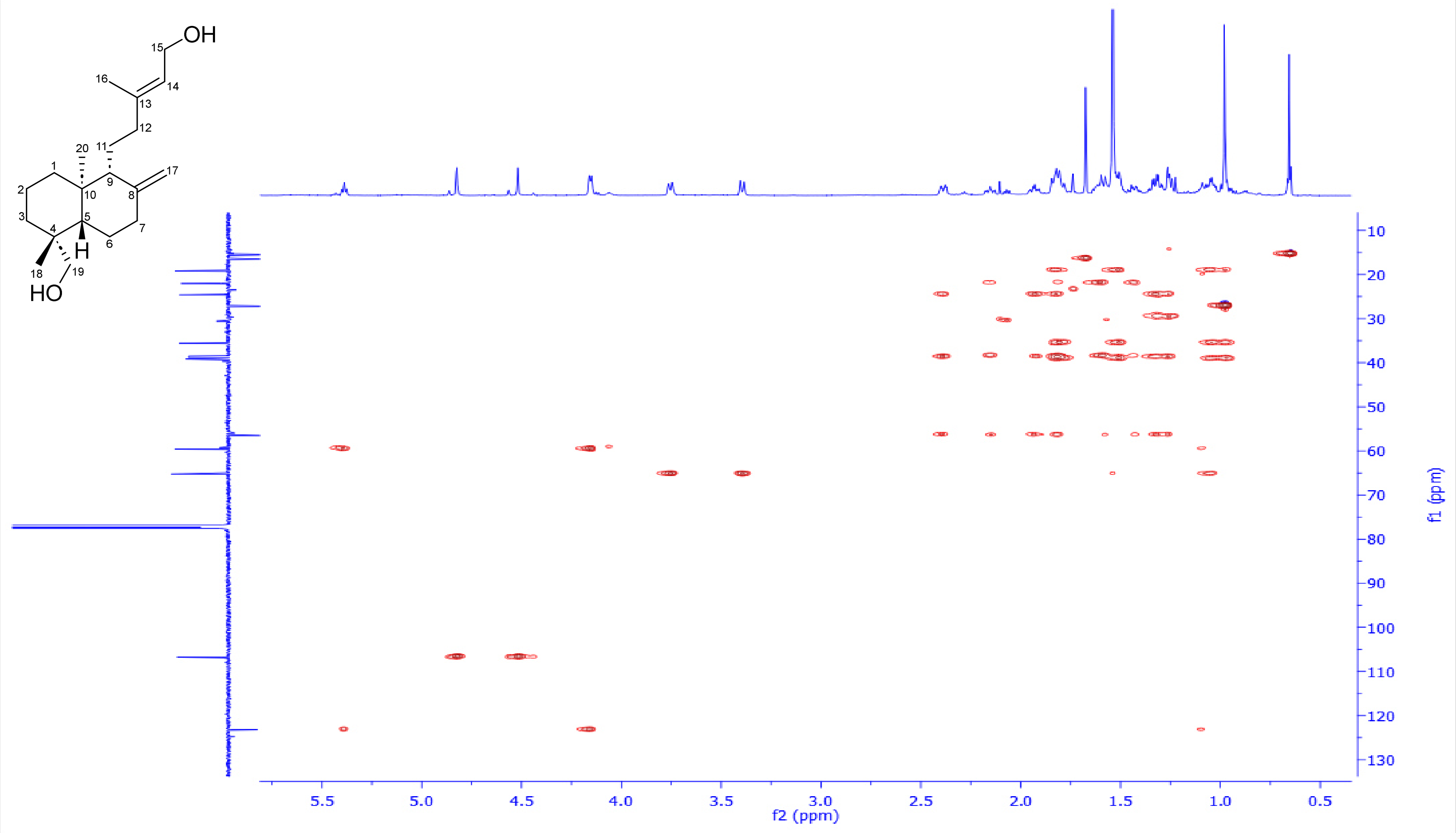

**Figure S32**: $^{1}H$-$^{13}C$ HMBC spectrum of *ent*-15,19-dihydroxy-labda-8(17),13(*E*)-diene **(14)** in $CDCl_3$, 600 and 150 MHz

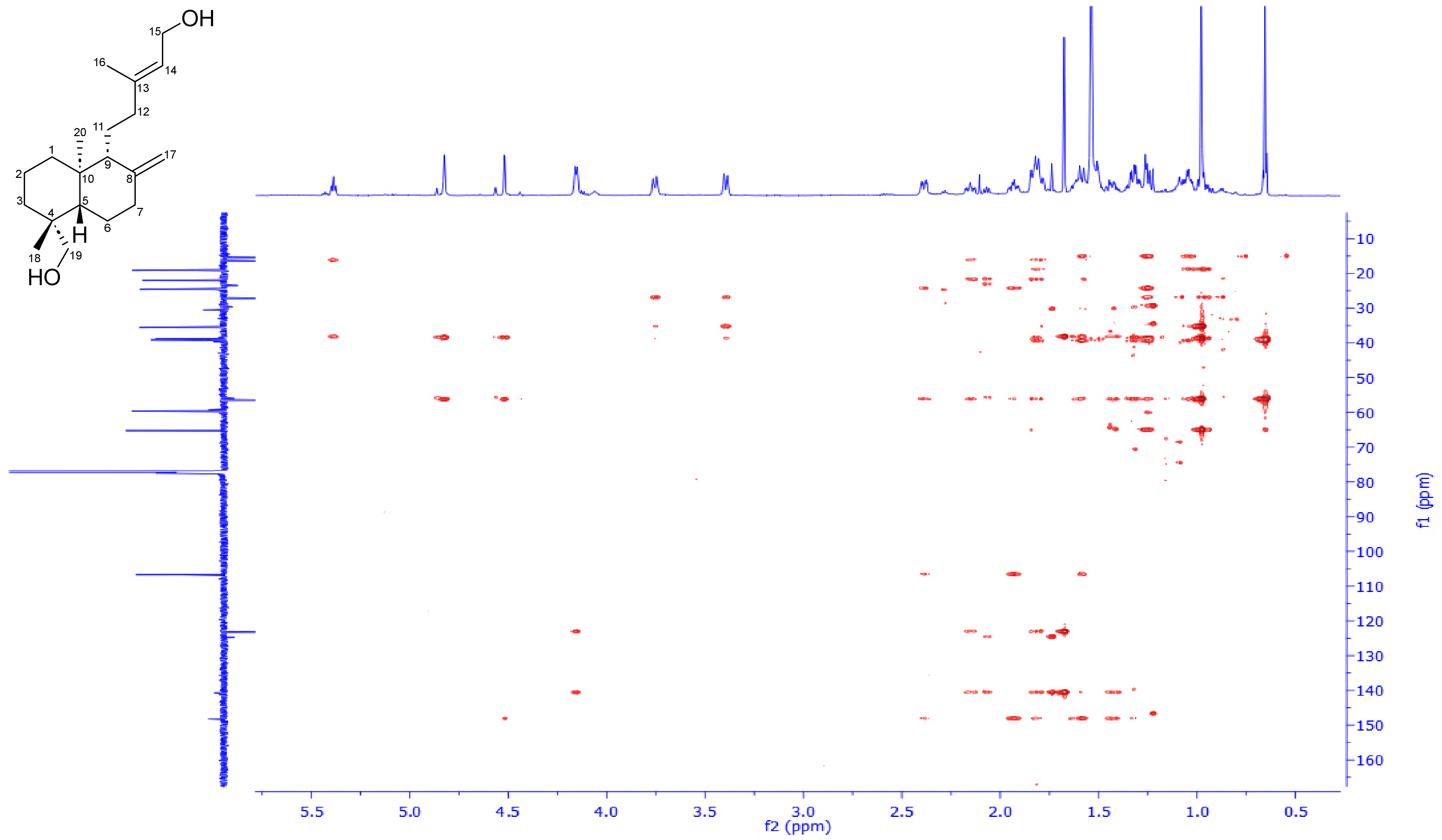

**Figure S33**: $^1$H-$^1$H ROESY spectrum of *ent*-15,19-dihydroxy-labda-8(17),13(*E*)-diene **(14)** in $CDCl_3$, 600 MHz

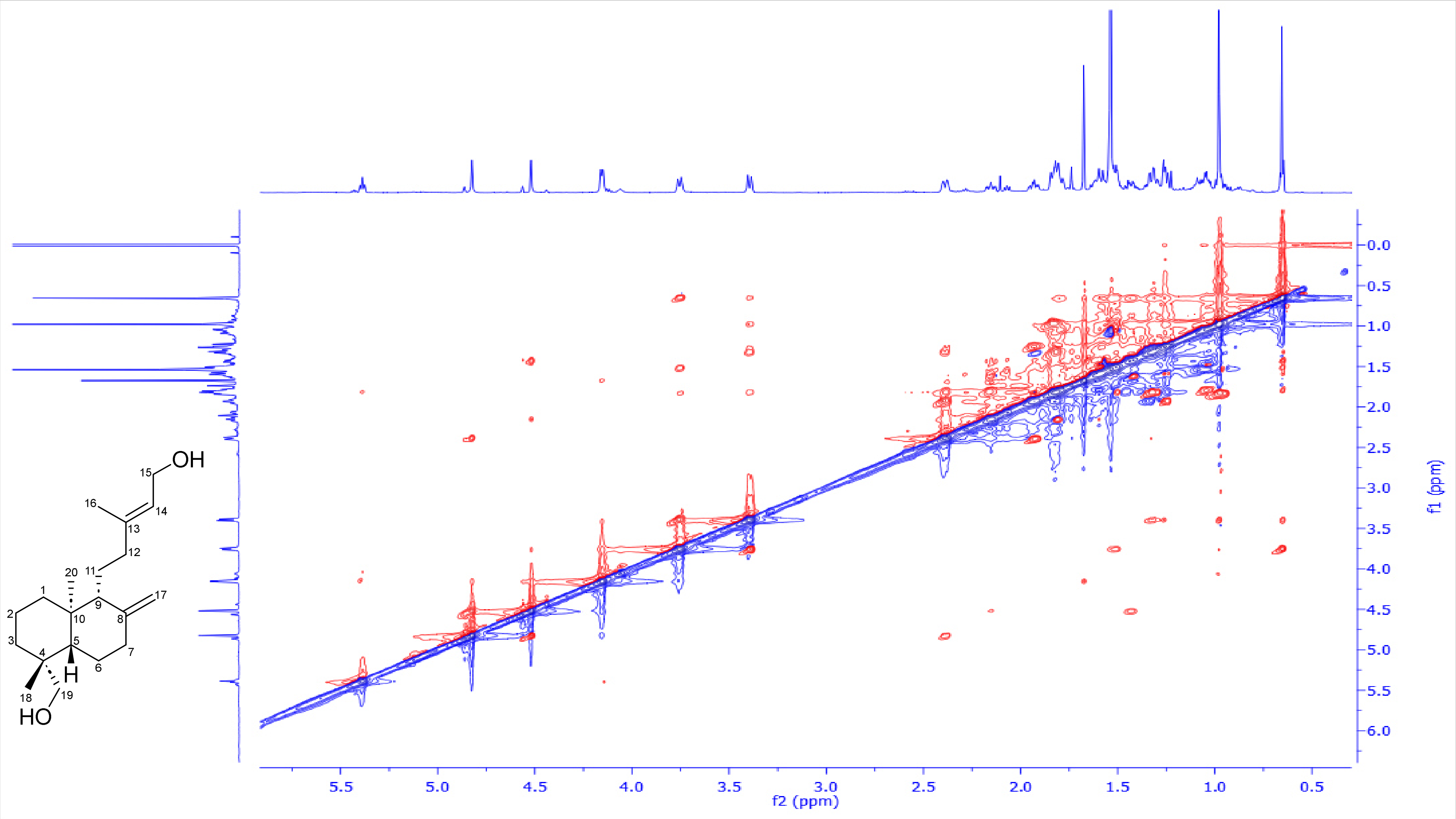

**Figure S34**: DEPTQ-135 spectrum of *ent*-15,19-dihydroxy-labda-8(17),13(*E*)-diene **(14)** in $CDCl_3$, 150 MHz

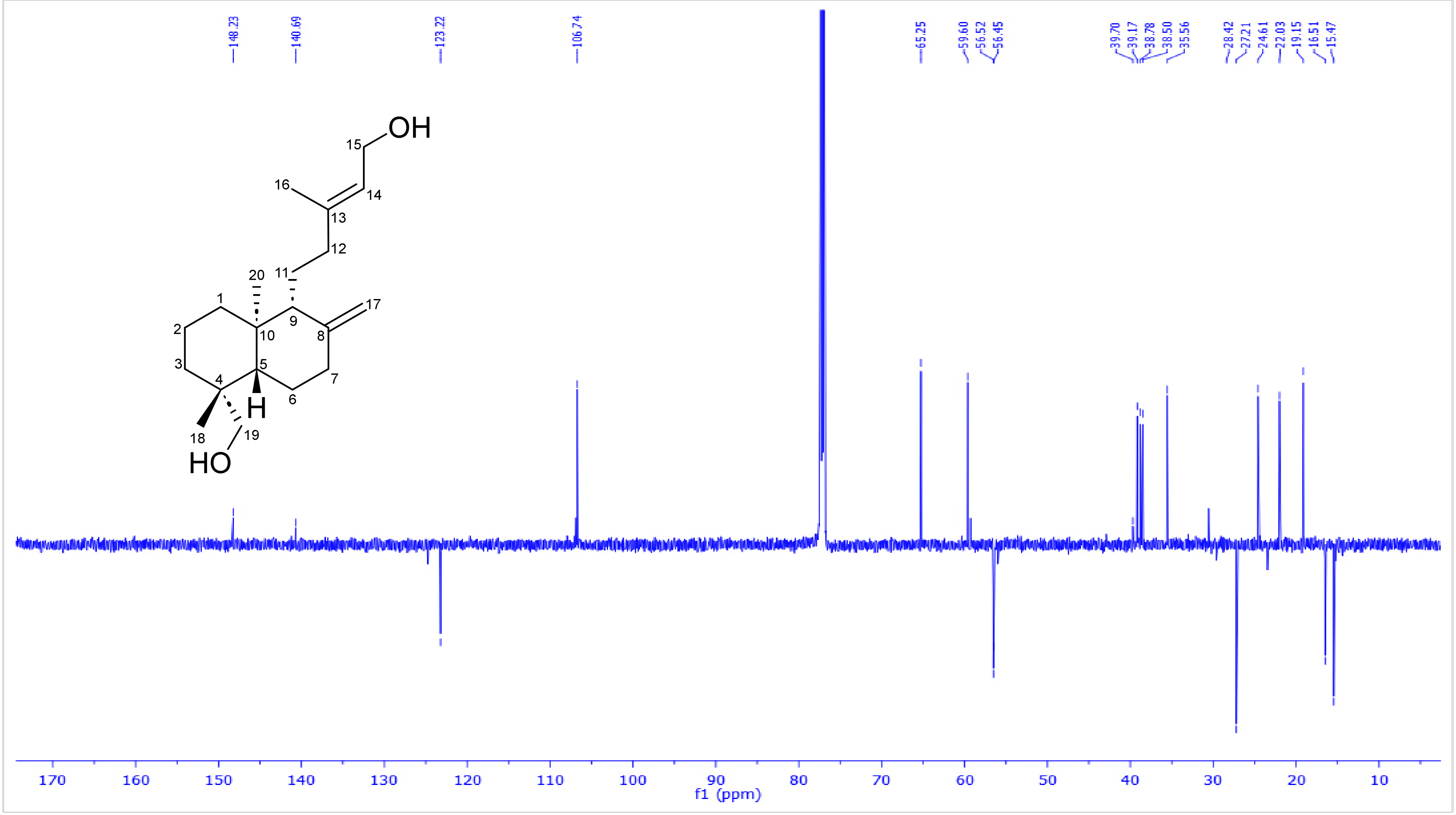

**5. Extraction, purification and structural characterization of *ent*-3α,15-dihydroxy-labda-8(17),13(*E*)-diene (15)**

*TaCPS_B1* and *TaCYP99A39_B2* were transiently co-expressed in **75** *Nicotiana benthamiana* plants to afford **100 g** of dry leaves. Subsequently, it was exhaustively extracted using ethyl acetate via speed extractor. The organic extract was collected and dried under reduced pressure to afford an intense green viscus material. Afterwards, it was introduced to three repetitive normal phase flash chromatography using a linear gradient of hexane/ethyl acetate [100/0 up 0/100] along 30 minutes, affording 12 subfractions F1-F12, where all have been monitored by LC-MS. Promising fractions F3-F4 were further purified using repetitive $C_{18}$ semi-preparative HPLC using a long gradient of water/acetonitrile [15/85 up 0/100], 4 mL/min, acidified with 0.1 % FA along 27 min to finally afford *ent*-3α,15-dihydroxy-labda-8(17),13(*E*)-diene **(15)** (**4** mg, pale yellow powder). Its full structure was resolved based on extensive 1D & 2D-NMR data analysis.

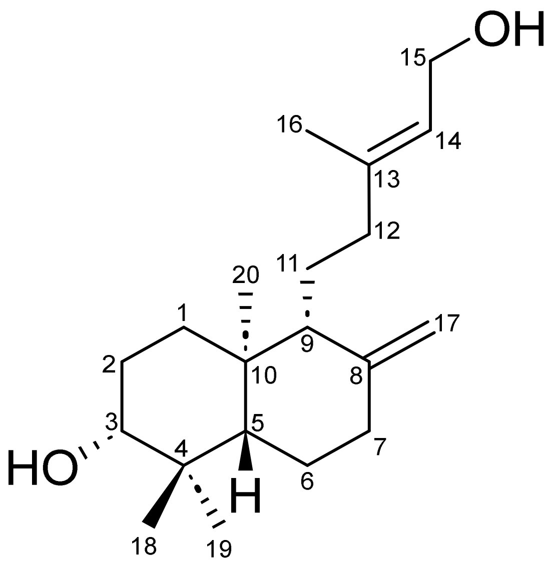

***ent*-3α,15-dihydroxy-labda-8(17),13(*E*)-diene (15)**

**Table 6**: [1]H, [13]C NMR spectroscopic data for *ent*-3$\alpha$,15-dihydroxy-labda-8(17),13(*E*)-diene **(15)** in $CDCl_3$, 600 and 150 MHz.

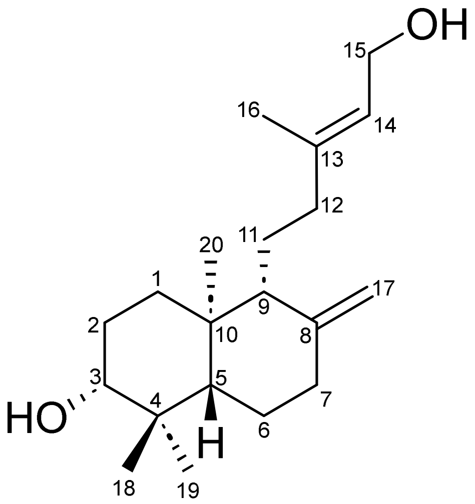

| Position | $\delta_C$, Type | $\delta_H$, mult. (*J* in Hz) |
|---|---|---|
| **1** | 37.2, $CH_2$ | 1.79, *m* |
| | | 1.16, *td* (13.3, 3.7) |
| **2** | 28.1, $CH_2$ | 1.70/1.59, *m* |
| **3** | 79.0, CH | 3.25, *dd* (11.8, 4.4) |
| **4** | 39.3, Cq | - |
| **5** | 54.8, CH | 1.08, dd (12.5, 2.7) |
| **6** | 24.1, $CH_2$ | 1.74, *m* |
| | | 1.39, *dd* (12.9, 4.2) |
| **7** | 38.3, $CH_2$ | 2.40, *ddd* (12.7, 4.0, 2.6) |
| | | 1.96, *td* (13.0, 5.1) |
| **8** | 148.1, Cq | - |
| **9** | 56.2, CH | 1.55, *m* |
| **10** | 39.5, Cq | - |
| **11** | 22.1, $CH_2$ | 1.58/1.46, *m* |
| **12** | 38.5, $CH_2$ | 2.15, *dd* (10.2, 4.1) |
| | | 1.81, *m* |
| **13** | 140.6, Cq | - |
| **14** | 123.3, CH | 5.39, *t* (6.6) |
| **15** | 59.6, $CH_2$ | 4.15, *d* (6.9) |
| **16** | 16.5, $CH_3$ | 1.67, *br s* |
| **17** | 106.9, $CH_2$ | 4.85/4.53, *br s* |
| **18** | 28.4, $CH_3$ | 0.99, *s* |
| **19** | 15.6, $CH_3$ | 0.77, *s* |
| **20** | 14.7, $CH_3$ | 0.69, *s* |

**Scheme 11**: Key $^1$H-$^1$H COSY (**bold blue**), $^1$H-$^{13}$C HMBC (H→C, red arrows) observed for *ent*-3α,15-dihydroxy-labda-8(17),13(*E*)-diene **(15)**

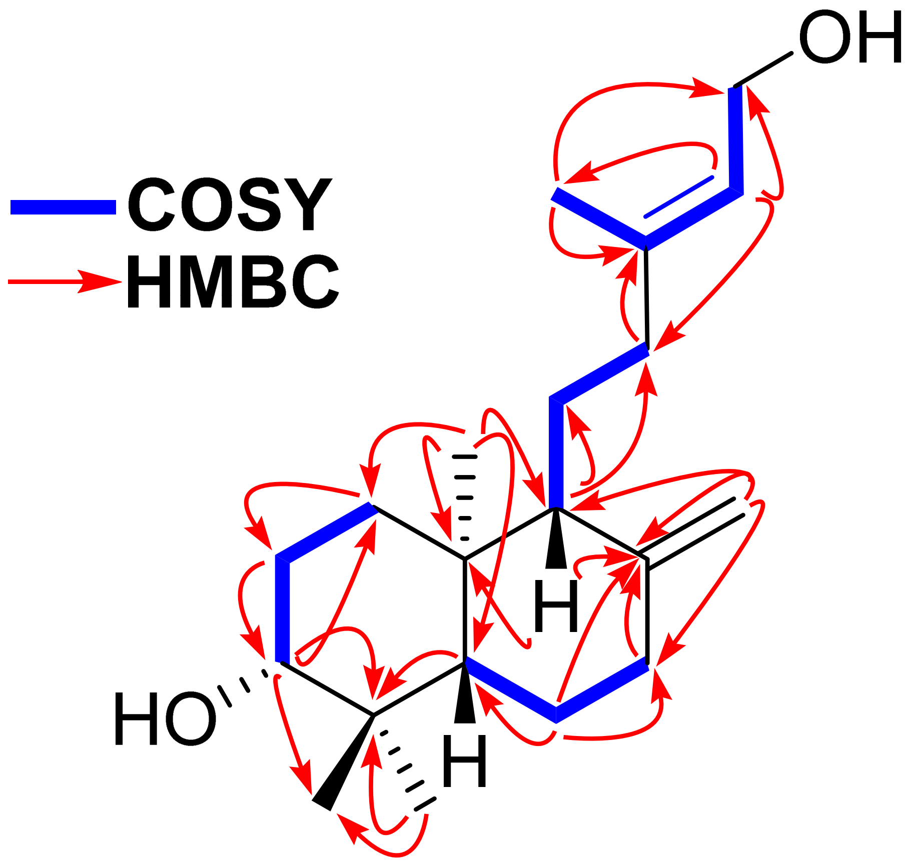

**Scheme 12**: Key ¹H-¹H ROESY (H↔H, green) observed for *ent*-3α,15-dihydroxy-labda-8(17),13(*E*)-diene **(15)**

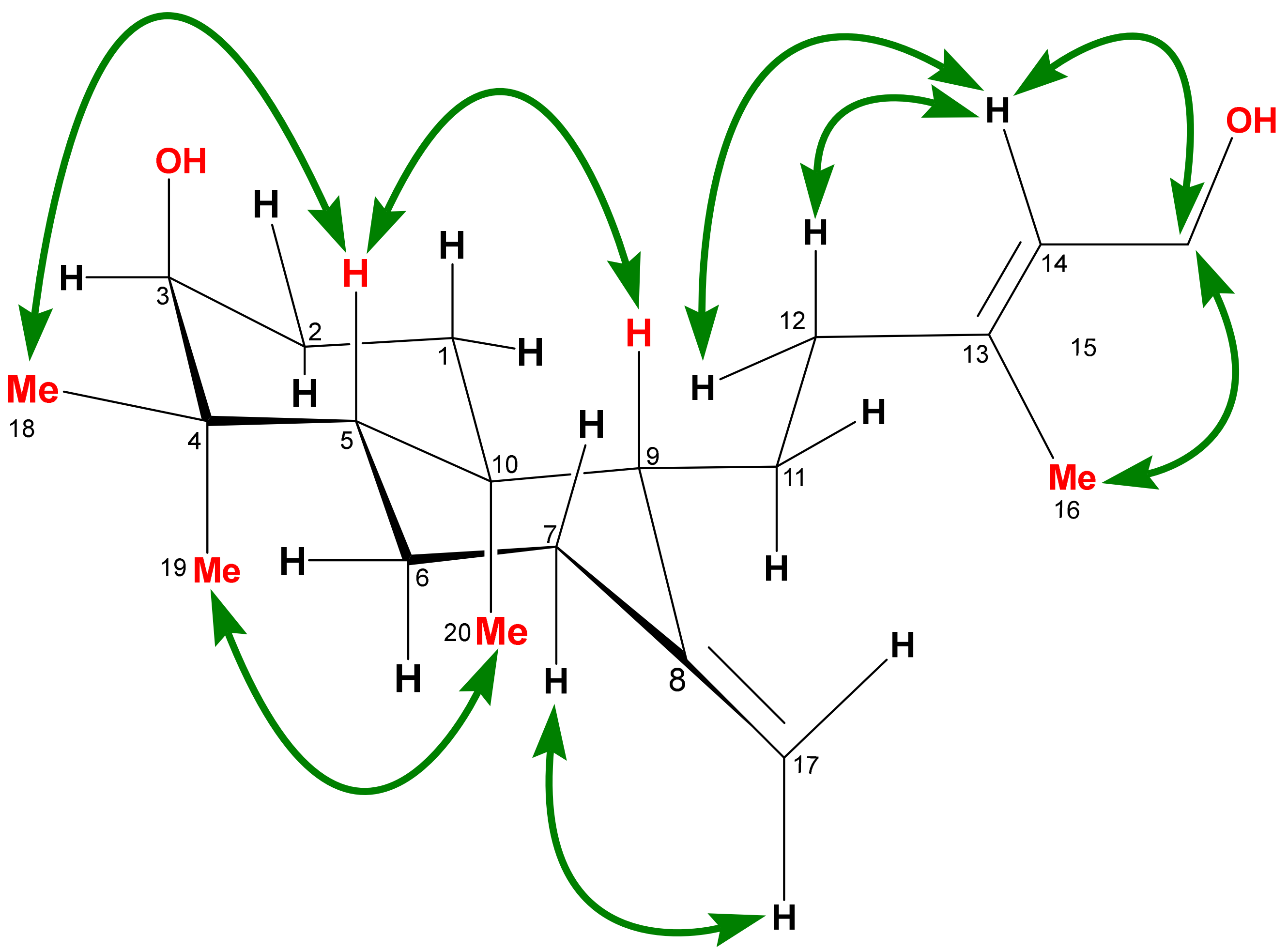

**Figure S35**: $^1$H-NMR spectrum of *ent*-3α,15-dihydroxy-labda-8(17),13(*E*)-diene **(15)** in $CDCl_3$, 600 MHz

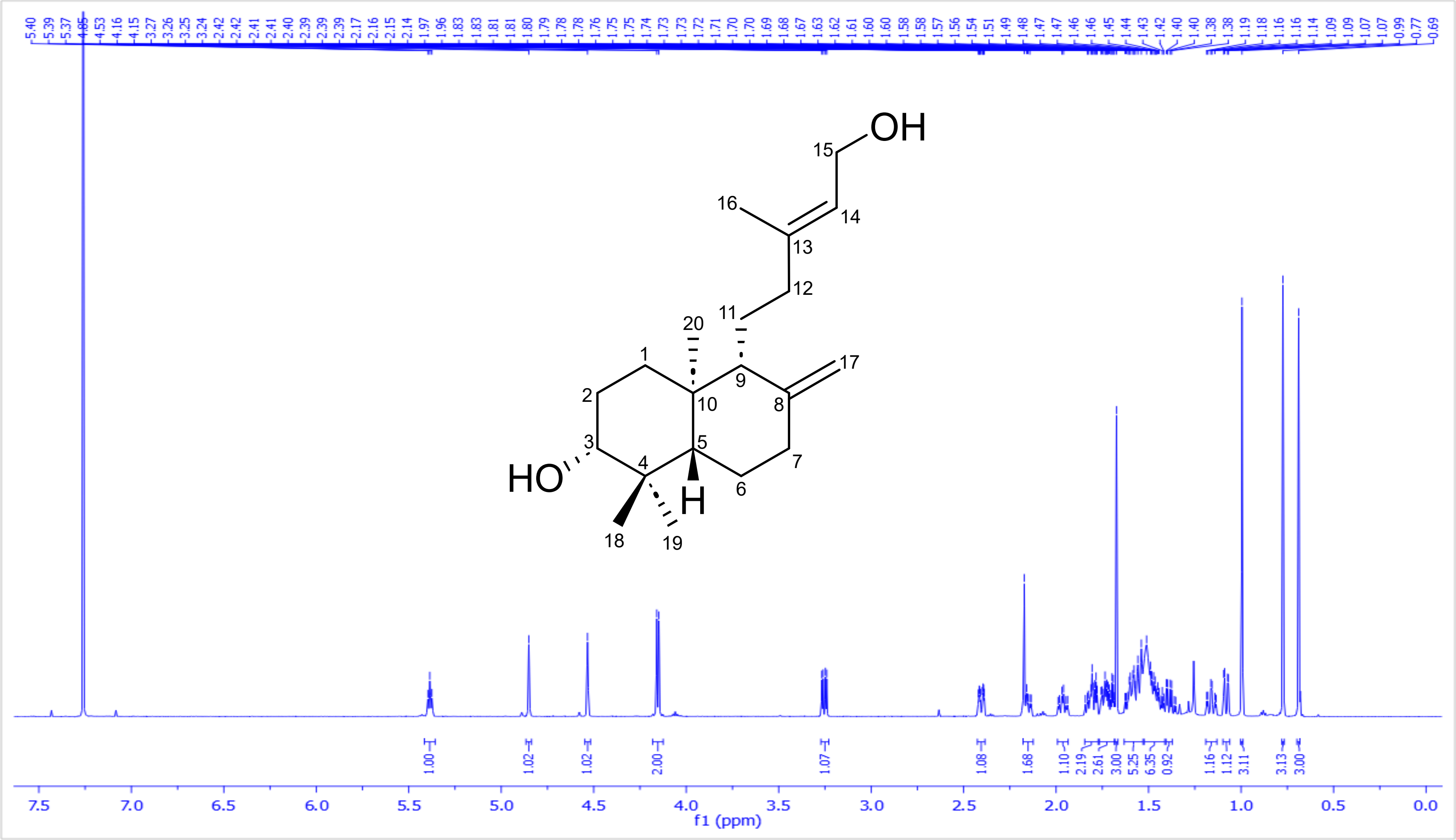

**Figure S36**: [1]H-[1]H COSY spectrum of *ent*-3α,15-dihydroxy-labda-8(17),13(*E*)-diene **(15)** in CDCl3, 600 MHz

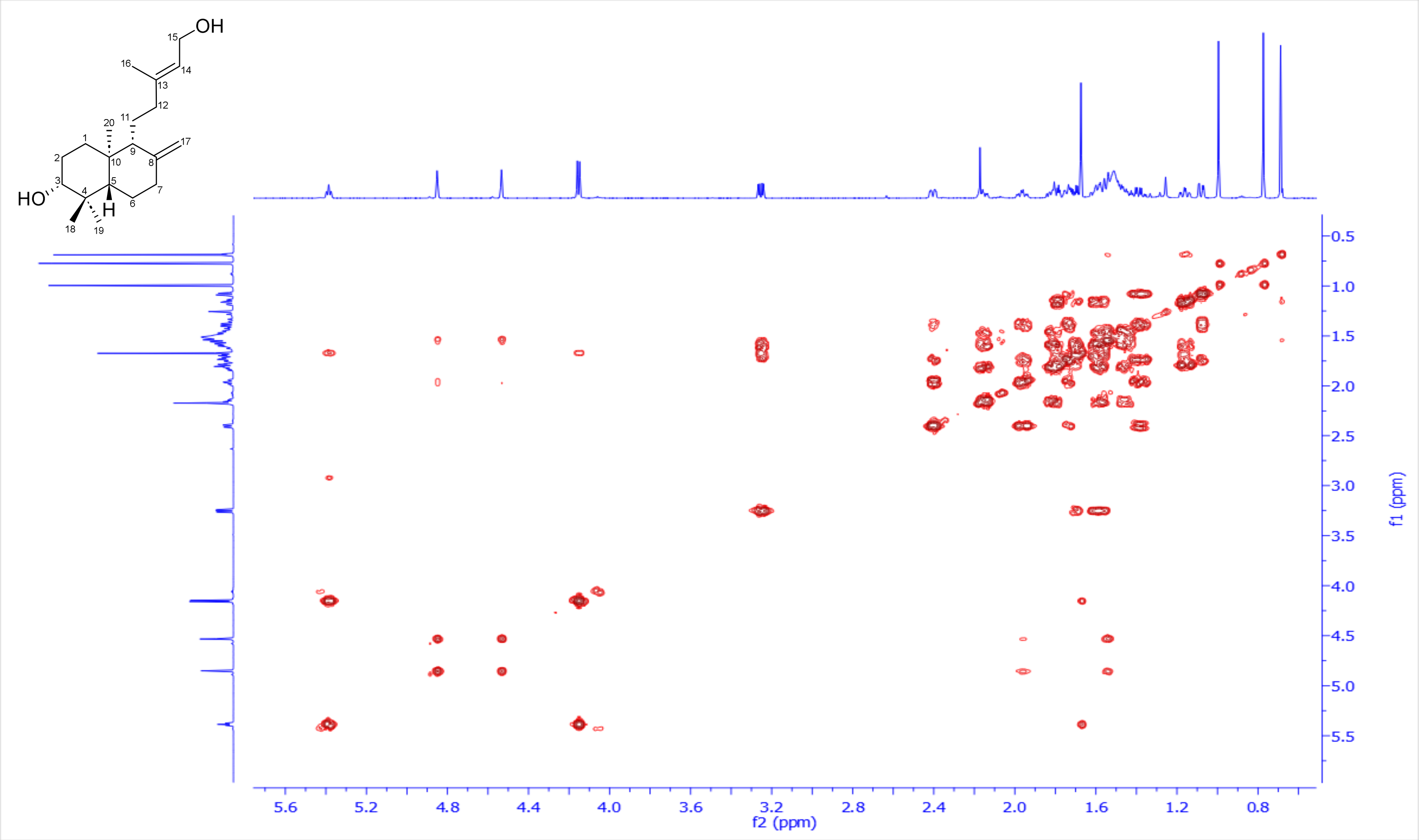

**Figure S37**: $^{1}H$-$^{13}C$ HSQC spectrum of *ent*-3α,15-dihydroxy-labda-8(17),13(*E*)-diene **(15)** in $CDCl_3$, 600 and 150 MHz

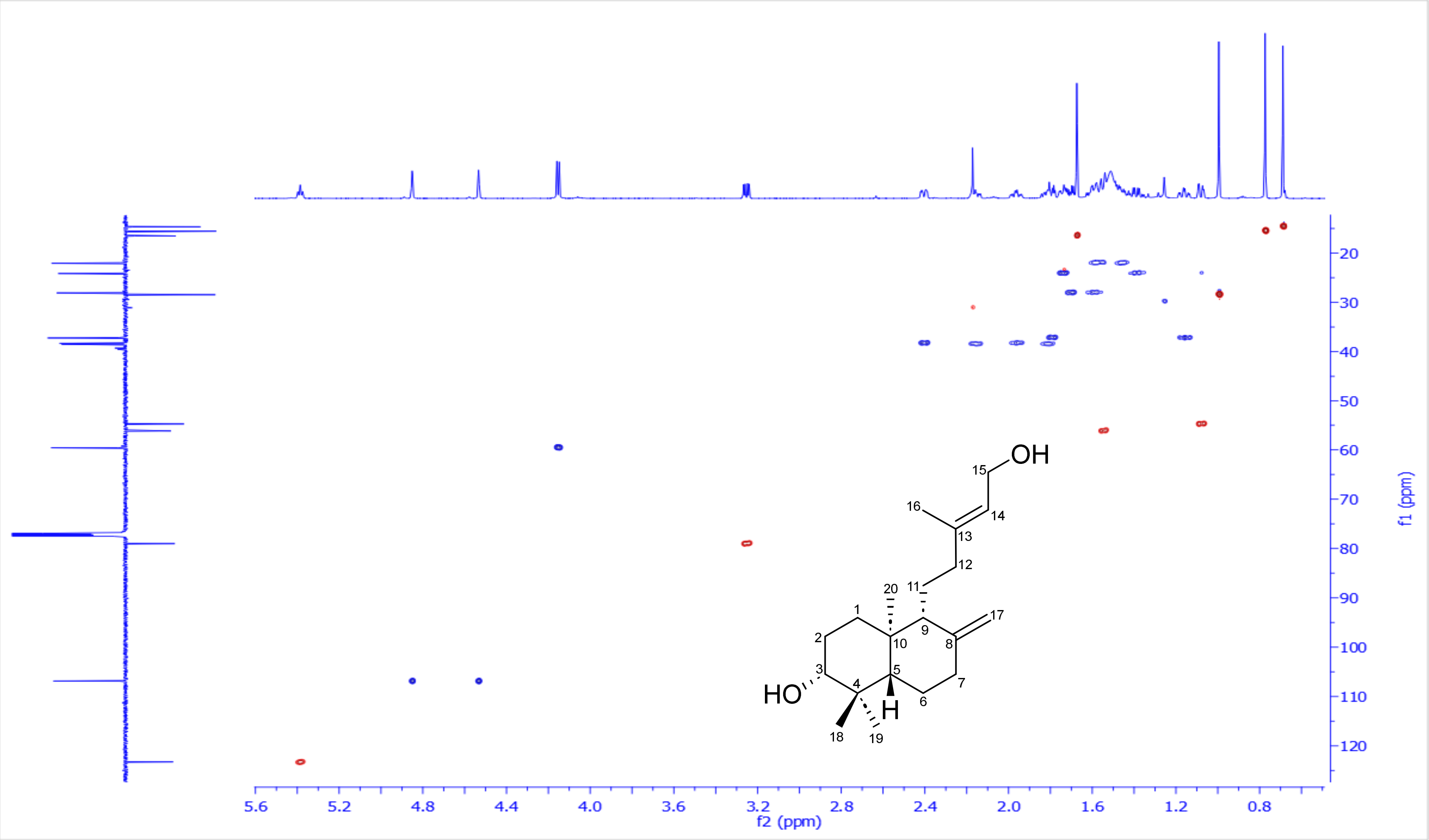

**Figure S38**: $^{1}H$-$^{13}C$ HSQC-TOCSY spectrum of *ent*-3α,15-dihydroxy-labda-8(17),13(*E*)-diene **(15)** in $CDCl_3$, 600 and 150 MHz

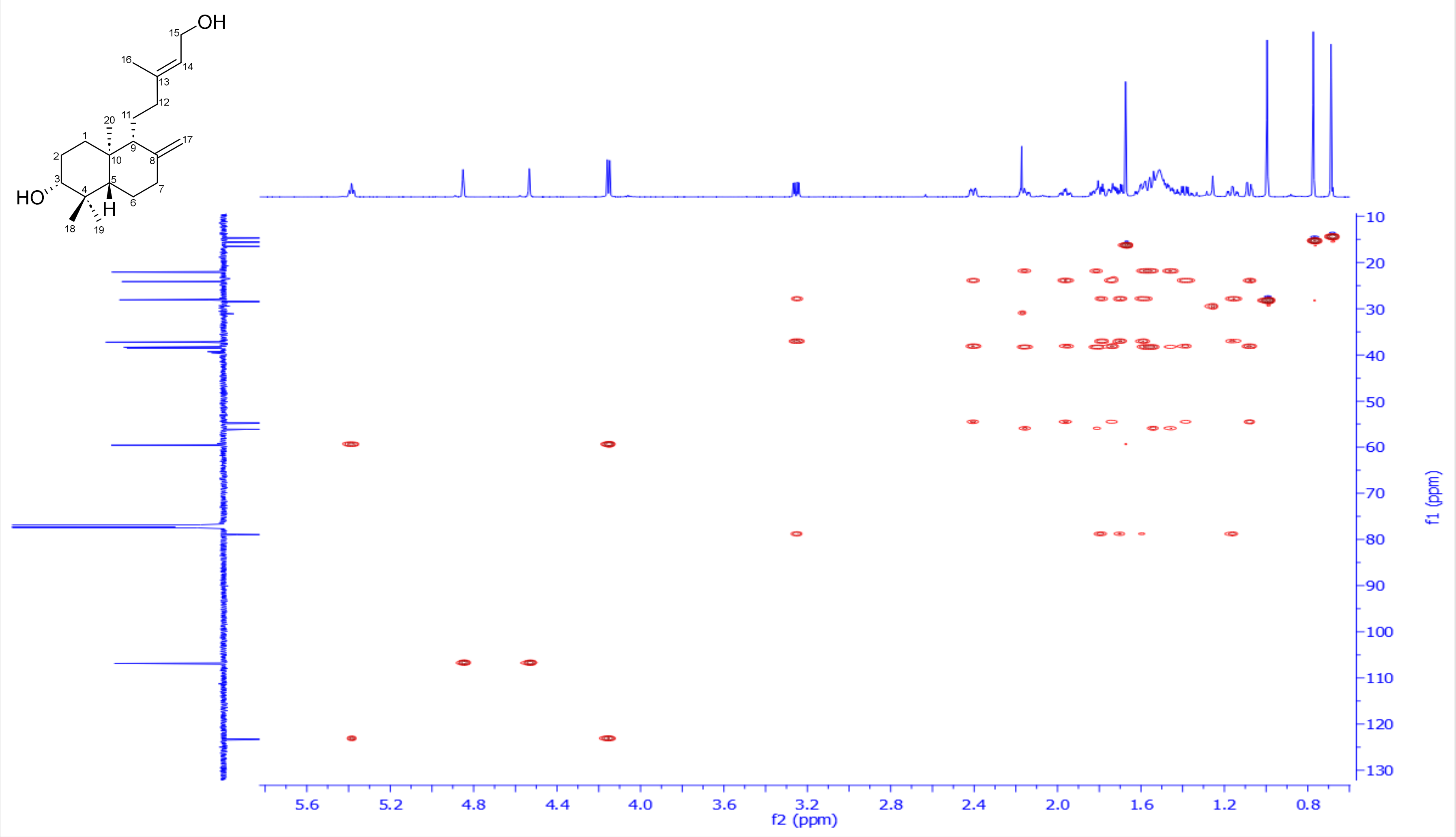

**Figure S39**: $^1$H-$^{13}$C HMBC spectrum of *ent*-3α,15-dihydroxy-labda-8(17),13(*E*)-diene **(15)** in $CDCl_3$, 600 and 150 MHz

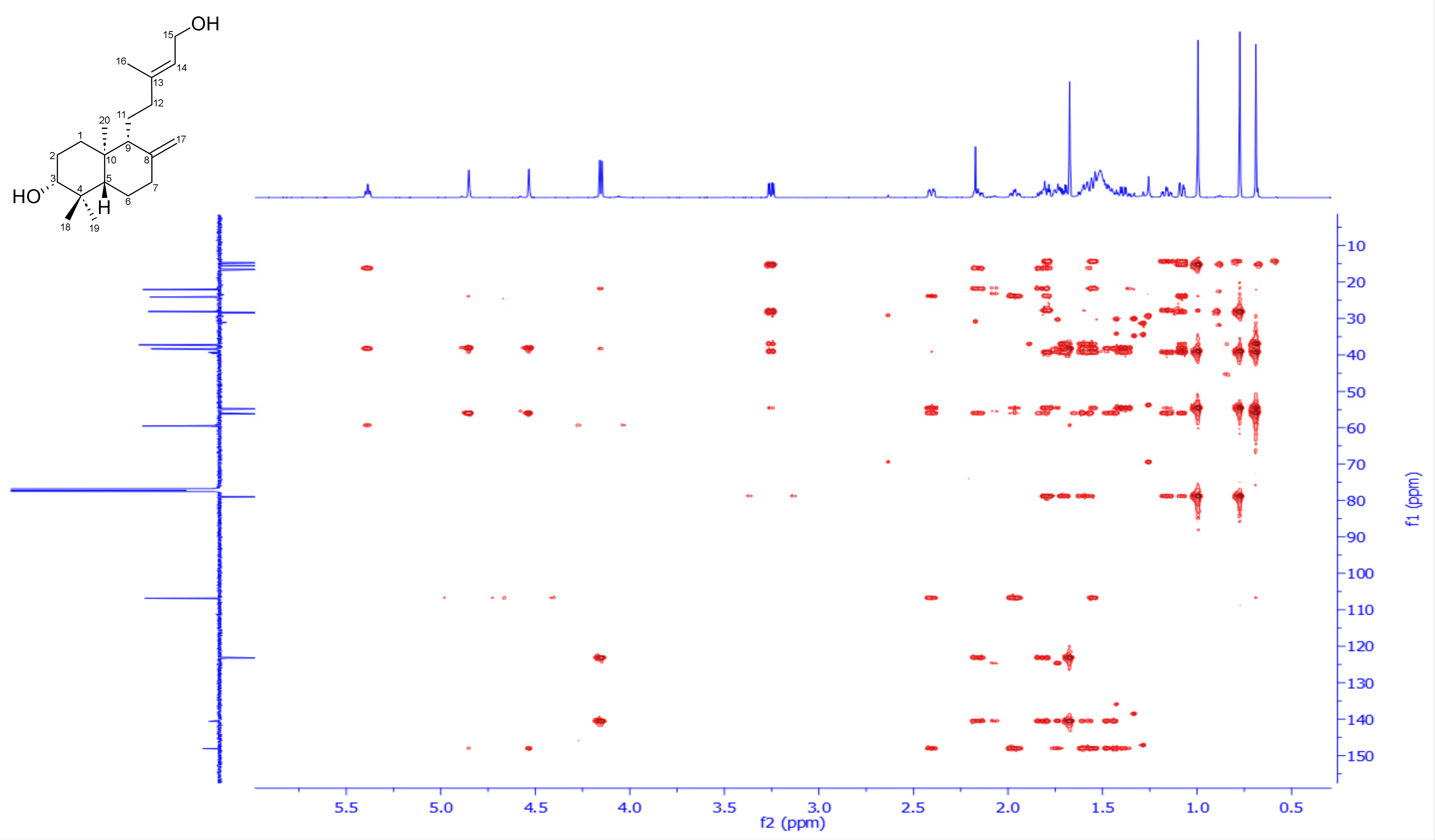

**Figure S40**: $^1$H-$^1$H ROESY spectrum of *ent*-3α,15-dihydroxy-labda-8(17),13(*E*)-diene **(15)** in $CDCl_3$, 600 MHz

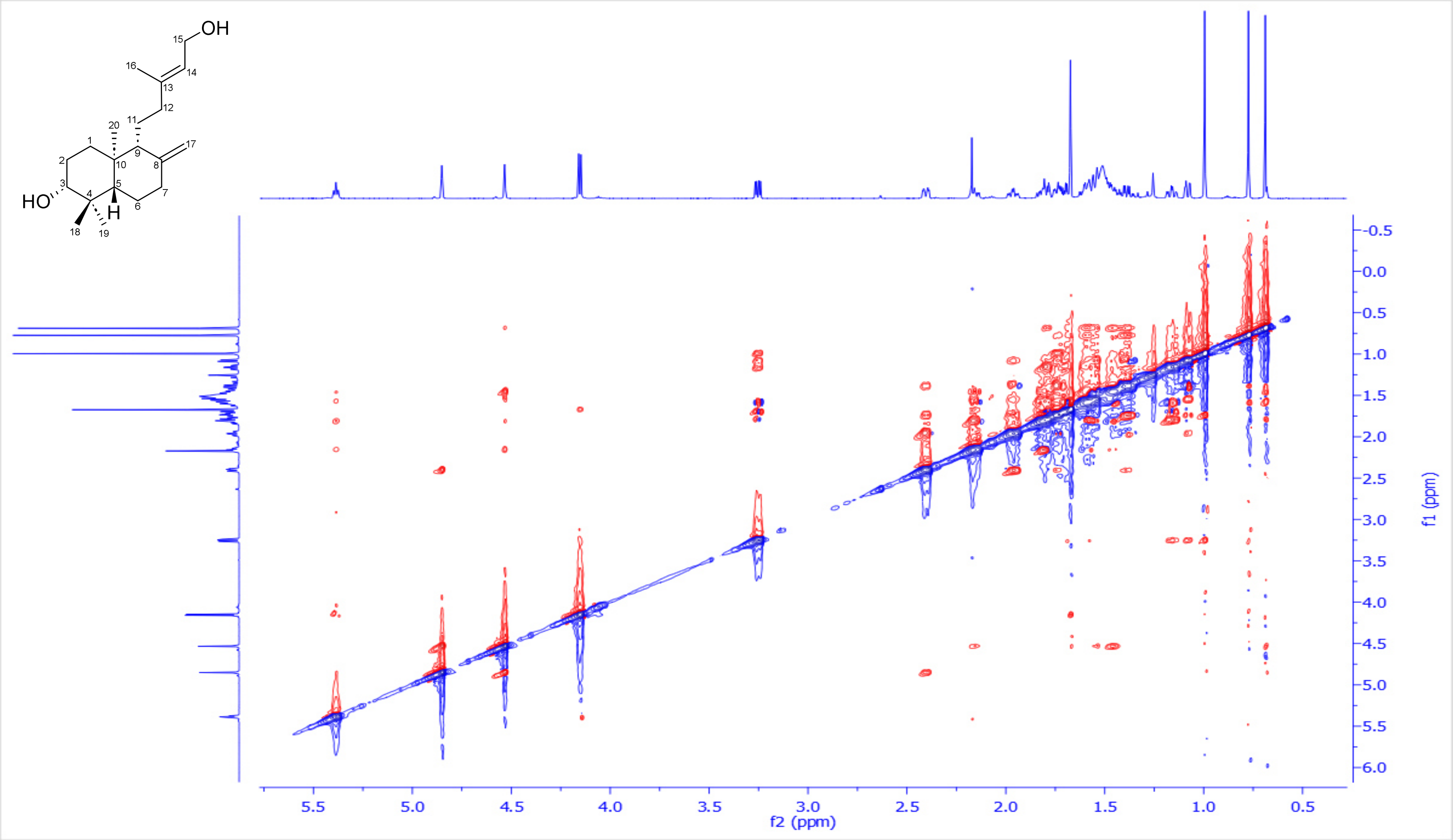

**Figure S41**: DEPTQ-135 spectrum of *ent*-3α,15-dihydroxy-labda-8(17),13(*E*)-diene **(15)** in $CDCl_3$, 150 MHz

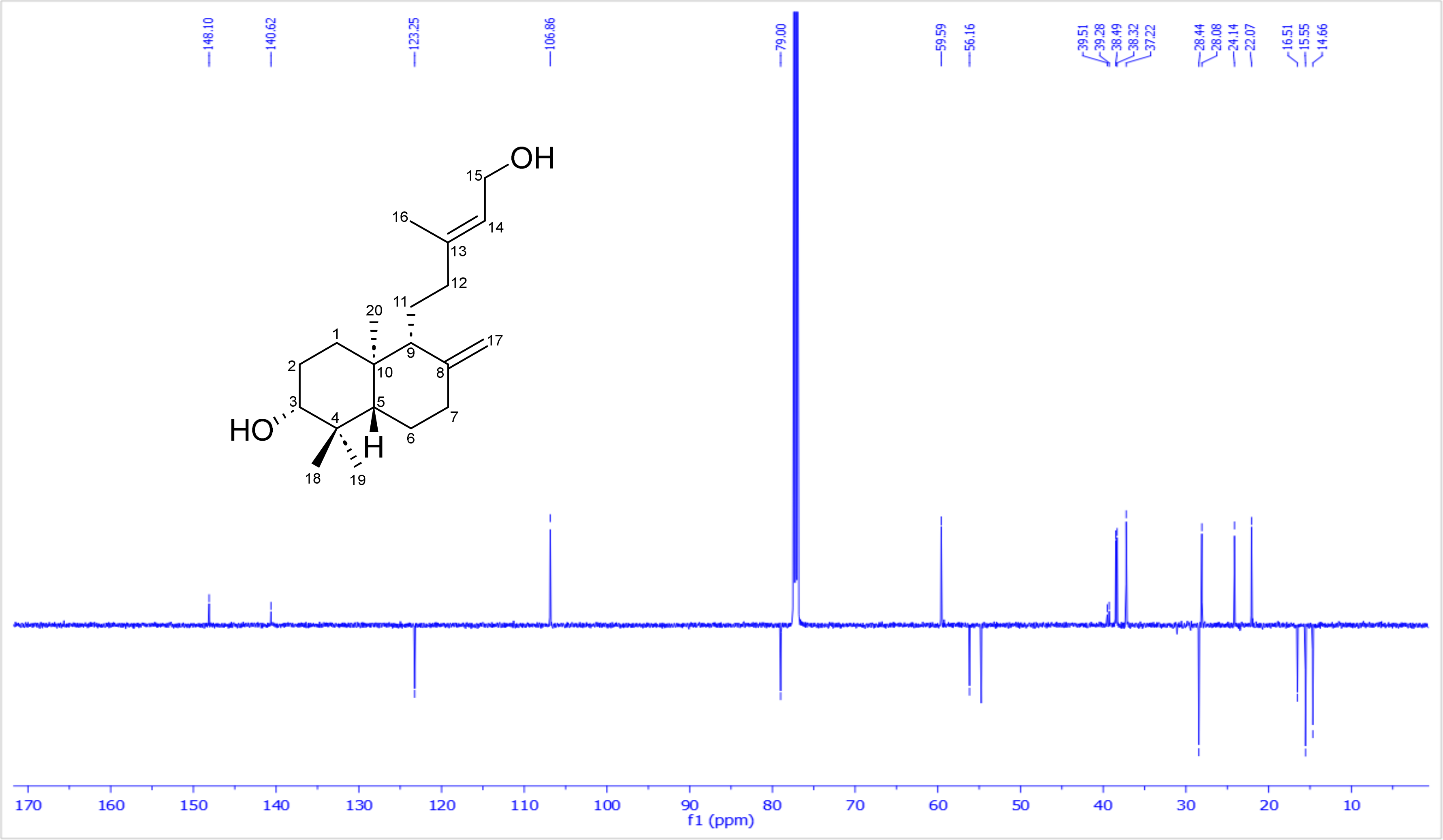
